# Pathogenic Bacteria Remotely Remodel Host Membrane Mechanics via Extracellular Vesicles to subvert Phagosome Maturation and Promote infection

**DOI:** 10.64898/2025.12.17.694930

**Authors:** Debraj Koiri, Lydia Mathew, Ayush Panda, Divit Solanki, Assirbad Behura, Shobhna Kapoor, Mohammed Saleem

**Author notes:** Equal contribution (Lydia Mathew, Ayush Panda).

## Abstract

Bacterial extracellular vesicles (BEVs) are known to enhance infection susceptibility in vivo, yet the mechanistic basis for this remote preconditioning of host cells is unknown. Here, we discover an evolutionarily conserved, lipid-driven physical mechanism by which pathogenic bacterial EVs systemically arrest phagosome maturation in bystander host cells. Using live-cell fluorescence lifetime imaging, *in vitro* reconstitution and micromanipulation, we show that EVs from diverse pathogens - *Mycobacterium tuberculosis*, *Klebsiella pneumoniae*, and *Staphylococcus aureus* - fuse with host plasma, phagosomal, and lysosomal membranes. This fusion increase membrane order and perturbs early phagosomal maturation. Transcriptomic profiling confirms a broad downregulation of phagosome maturation genes while upregulation of lysosomal stress responsive genes. Crucially, in vitro reconstitution shows that EVs, and their purified lipids alone, are sufficient to induce phase separation and increase membrane order, directly inhibiting phago-lysosomal fusion. Our findings establish a paradigm in which pathogens exploit EVs not merely as delivery vehicles, but as tools to remotely rewire host cell membrane mechanics to hijack phagosome maturation and promote infection - a strategy that moves beyond canonical effector-based models of pathogenesis.

## Introduction

Phagosome maturation, a fundamental host defense mechanism, mediates pathogen degradation through phagolysosome formation ^1^.This involves the sequential fusion of phagosomes with lysosomes tightly orchestrated via membrane trafficking and organelle acidification ^2,3^. Several pathogens have evolved sophisticated strategies to disrupt phagosome maturation, enabling their survival and replication within macrophages ^4–6^. Following the initial discovery that *Mycobacterium tuberculosis* inhibits phagosome-lysosome fusion^7^, numerous studies have identified the bacterial effectors responsible for this survival strategy. For example the ESX-1 secretion system ^8,9^, secreted lipid virulence factors^10–13^ and secreted phosphatases^14,15^ are all known to interfere with host signalling. An abnormal retention of early endosomal markers like Rab5 and the exclusion of late endosomal markers like Rab7 and the v-ATPase complex is also known to result in an aberrant phagosome ^16^. The failure of phagosome lysosome fusion is attributed to the reduced accumulation of phosphatidylinositol-3-phosphate (PI3P) on the phagosomal membrane, which diminishes the recruitment of effector proteins essential for lysosomal fusion ^17^. Mycobacterial lipids like lipoarabinomannan (LAM) and enzymes such as the phosphatases SapM and PtpA are also known to contribute to this process ^18,19^. Despite decades of pursuit our understanding of the mechanism of how these effectors, particularly the Mycobacterial lipids, are trafficked to and function at the host-pathogen interface to orchestrate phagosome maturation arrest is still an intensive and unresolved area of research ^20^. *Mycobacterium tuberculosis* (Mtb) secretes nanoscale extracellular vesicles (MEVs) that function as delivery vehicles, transporting a payload of bacterial lipids, proteins, and nucleic acids directly into host cells^21^. Previous research has established that these vesicles play a significant immunomodulatory role, such as by inhibiting T-cell activation ^22–24^. However, the fundamental question of how MEVs interact with and alter host intracellular compartments, their precise intracellular fate and mechanism of action has remained unknown.

We discovered that pathogenic bacterial extracellular vesicles (BEVs) across bacterial kingdom (i.e, *Mycobacteriuam tuberculosis, S. aureus* and *K. pneumonia*) spontaneously fuse with the host organellar membranes and remodel their mechanical parameters. Using *Mycobacterium tuberculosis* as a primary model, we determined that extracellular vesicles upon fusion subsequently enrich their lipids in the phago-lysosomal membranes dramatically increasing the order. This mechanical alteration delays the recruitment of the key GTPase Rab5, a master regulator that initiates phagosome maturation. This elevated membrane order persists even after the phagosome fuses with a lysosome. Lipidomic analysis confirmed that this effect is driven by the mycobacterial extracellular vesicle (MEV)-mediated enrichment of mycobacterial lipids within host membranes, predominantly inositol derivatives like Lipoarabinomannan (LAM), Phthiocerol Dimycocerosate (PDIM), and Phosphatidylinositol Mannosides (PIMs). *In vitro* reconstitution demonstrates that the purified MEV lipid fraction is alone sufficient to increase membrane order and drive phase separation creating a mechanical barrier that opposes the fusion of phagosomes with lysosomes. Further, MEV lipids also elevate lysosomal membrane anisotropy evident from lifetime imaging, disrupting lysosomal homeostasis and further impairing its ability to fuse. Surprisingly, the physical manipulation of host membranes is complemented by transcriptional reprogramming of the host cell’s phagosome maturation and membrane repair machinery to sustain the blocked state of maturation. Together, Our study discovers a novel, evolutionarily conserved paradigm that bacterial pathogens exploit extracellular vesicles not merely as delivery vehicles, but as tools to remotely manipulate the physical parameters of host organelle membranes, thereby hijacking a central pillar of innate immunity i.e. phagosome maturation.

## Results

### Mycobacterial Extracellular Vesicles (MEVs) Spontaneously Fuse With The Host Plasma Membrane

We first cultured *M.tb* H37Ra under minimal nutrient conditions to induce vesicle production. Transmission electron microscopy (TEM) revealed distinct stages of Mycobacterial extracellular vesicles (MEVs) biogenesis, including membrane protrusion and bud formation from the bacterial surface (Fig. 1A). We then isolated and purified MEVs, adopting a vesicle isolation protocol previously established for *M. tuberculosis* H37Rv ^22^, involving sequential differential centrifugation followed by OptiPrep density gradient ultracentrifugation. Vesicle-enriched fractions from the culture supernatants of H37Ra grown under minimal nutrient conditions revealed discrete, bilayered vesicular structures ranging from ∼ 120 to 300 nm in diameter revealed by nanoparticle tracking analyzer and dynamic light scattering consistent with the known size distribution of MEVs (Fig. S1 A-C, Supplementary movie 1). Likewise, the biochemical signatures of the MEVs such as the derivatives of Lipoarabinomannan (LAM), Phthiocerol Dimycocerosate (PDIM), and Phosphatidylinositol Mannosides (PIM) were also identified using lipidomics (Fig.S13B). MEVs exhibited consistently higher anisotropy than host cell membranes, indicating that MEVs are intrinsically more ordered and rigid as evident from their fluorescence anisotropy through DPH/TMA-DPH probes as a function of time (Fig. S1 H-I).To visualize the initial interaction between *M.tb* H37Ra extracellular vesicles *(MEVs)* and host macrophages in real-time, PMA-differentiated THP-1 macrophages were treated with PKH67-labeled MEVs and monitored using live-cell fluorescence microscopy (Fig. 1B). We ruled out any dye related artifacts such as dye aggregates by removal of the free dyes through size exclusion chromatography and rigorous washing to remove residual dye as previously reported ^25^. Further, we confirmed the same by treating PKH67 dye with macrophage that indicated a diffused signal of the dye (Fig.S2A). Further, we also confirmed that fluorescence labelling of the MEVs did not significantly change its size or zeta potential (Fig.S2B-C). Within 30 minutes of treatment, a subset of MEVs accumulated and likely fused with the macrophage plasma membrane, seen as a distinct spread of the fluorescent signal at the cell’s periphery (Fig. 1B, S3A). Quantitative analysis of membrane-associated fluorescence revealed ∼ 40% rise in the fluorescence intensity corresponding to docking or fusion events at the host plasma membrane (Fig. 1C). To further validate MEV - host membrane fusion, we employed a membrane fusion assay based on R18 dye fluorescence dequenching. R18 is a self-quenching lipophilic dye that exhibits increased fluorescence upon dilution in the lipid bilayer of a target membrane during fusion events ^26^. R18-labeled MEVs were incubated with large unilamellar vesicles (LUVs) mimicking the host plasma membrane composition, and fluorescence intensity was monitored over time. The fluorescence of the R18 dye increased over time because the dye molecules spread out as the membranes fused (Fig. S3B). To reconfirm membrane fusion and assess lipid mixing between MEVs and host membranes, a kinetic fluorescence resonance energy transfer (FRET)-based lipid mixing assay was employed. In this assay, donor (NBD-PE) and acceptor (Rh-PE) fluorophores were incorporated into host plasma membrane-mimicking LUVs. Upon fusion with unlabeled MEVs, dilution of the donor-acceptor pair leads to a concomitant increase in donor fluorescence, indicating lipid mixing ^27,28^. Consistent with the R18 dequenching results, we observed a time-dependent increase in NBD fluorescence intensity following MEV addition, suggesting that MEVs actively fuse and exchange lipids with the model host membrane (Fig. 1D). In contrast, LUVs incubated in the absence of MEVs exhibited no appreciable change in donor fluorescence, confirming that the observed signal increase was specific to MEV-mediated fusion.

**Figure 1.**
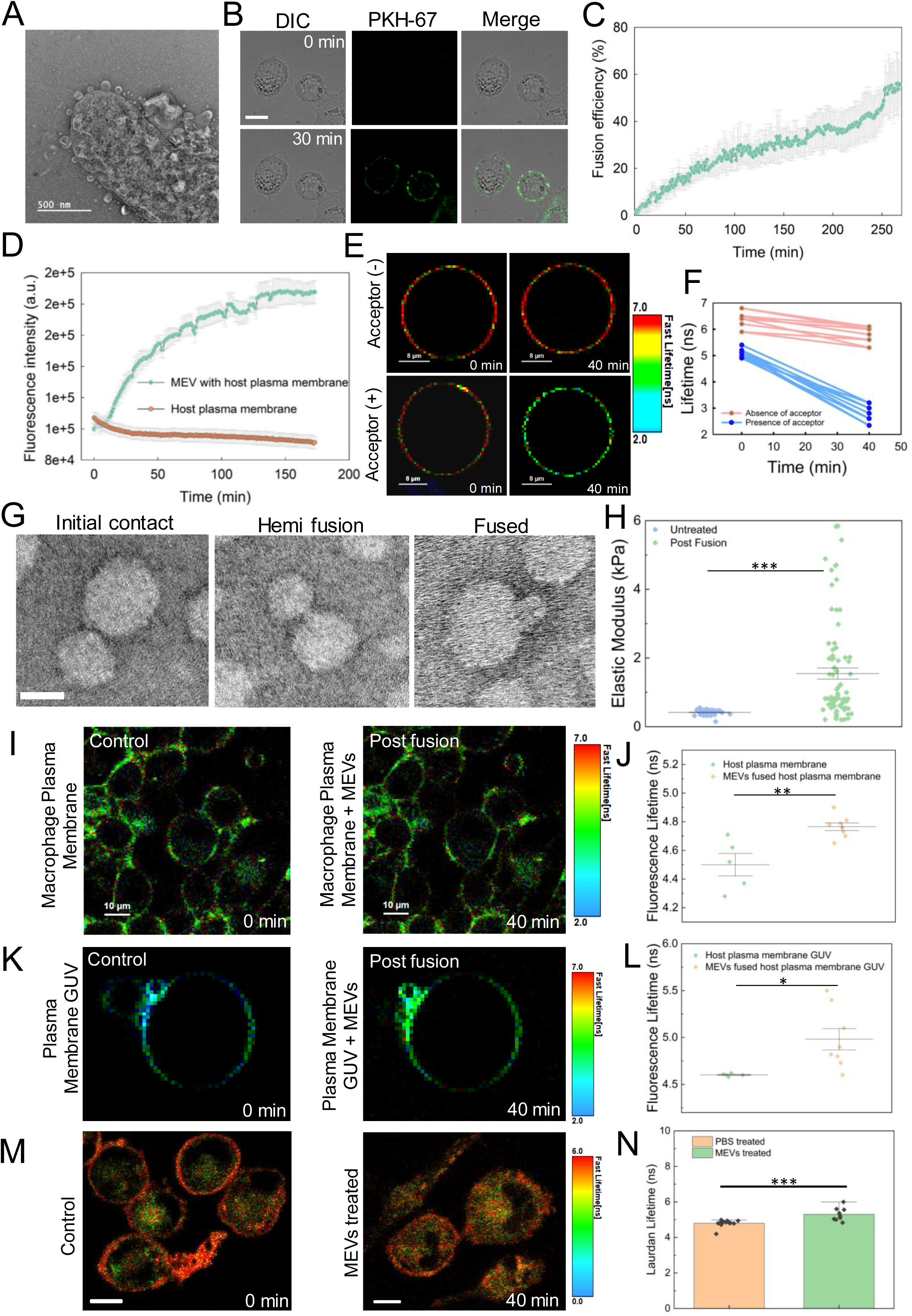
Mycobacterial extracellular vesicles (MEVs) spontaneously fuse with the host plasma membrane and remodel the membrane mechanics. (a) Transmission electron micrographs representing distinct stages of mycobacterial extracellular vesicles (MEVs) formation and release from the bacterial envelope, highlighting membrane protrusion, budding, and detachment events. (b) Time-lapse confocal imaging of dTHP-1 macrophages incubated with PKH-67 labelled MEVs (green). Scale bar, 10μm. Images are representative of five independent experiments (n ∼250 cells). (c) Fusion kinetics were evaluated by monitoring the time-dependent increase in fluorescence intensity, reflecting lipid and/or content mixing between MEVs and the host plasma membrane. Nonlinear regression fits illustrate the progression of fusion over time. Data points represent mean ± S.D. from three independent experiments. (d) Förster resonance energy transfer (FRET) assay was performed with plasma membrane LUVs labelled with 1% NBD-PE and 1% Rh-PE. Donor fluorescence intensity was monitored as a function of time in the presence and absence of MEVs. Data points are shown as the means ± S.D. of three independent measurements. All experiments are carried out in PBS pH 7.4 at 37°C. (e&f) Fluorescence lifetime imaging microscopy-based Förster resonance energy transfer (FLIM-FRET) assay was performed with the plasma membrane GUV labelled with NBD-PE and the MEVs were labelled with R18. Donor Fluorescence Lifetime was monitored over a period of 40 minutes in the presence and absence of acceptor. Images are the representative of five independent experiments (n ∼35 GUVs). Data points are shown from three independent experiments. All experiments are carried out in PBS pH 7.4 at 37°C. Scale bar, 8μm. (g) Transmission electron micrographs representing distinct stages of MEV fusion with the membrane. Scale bar, 100nm. (h) Mean elastic modulus values of dTHP-1 macrophage surfaces measured by atomic force microscopy (AFM) show an increase in cellular stiffness following MV fusion relative to untreated (only PBS) macrophages. Data are presented as mean ± SEM from ≥ 60 cells per condition. Fluorescence lifetime imaging microscopy (FLIM) analysis of membrane biophysical changes during MEV fusion. (i) FLIM images of Flipper-Tr labelled live macrophages before and after MEV fusion. (j) Quantification of Flipper-Tr fluorescence lifetimes in macrophages as shown in (i). (k) FLIM images of Flipper-Tr -incorporated plasma membrane–mimicking GUVs before and after exposure to MEVs. (l) Corresponding fluorescence lifetime measurements of GUVs as shown in (k). Data points in (j) and (l) represent mean ± S.D. from each independent experiment. (m) FLIM images of Laurdan lifetime incorporated in Thp-1 derived macrophages in the presence of MEVs and untreated (only PBS treated). Scale bar: 10 μm. (n)Corresponding fluorescence lifetime measurements of macrophages as shown in (m). Data points represent mean ± S.D. from each independent frame (n∼5 cells at each frame). (**p<0.01, ***p<0.001 in one-way ANOVA).

Since intensity-based FRET measurements can sometimes be prone to artifacts such as photobleaching and donor dye dilution during membrane fusion, we employed fluorescence lifetime imaging microscopy (FLIM)-FRET to more accurately assess MEV fusion. R18-labeled MEVs were incubated with NBD-PE labeled giant unilamellar vesicles (GUVs) mimicking the host plasma membrane. NBD-PE fluorescence lifetime was quantified over 40 minutes in the presence and absence of the acceptor. A significant decrease in NBD lifetime was observed in the presence of R18-labeled MEVs, indicating efficient FRET and confirming membrane fusion between MEVs and the GUVs (Fig. 1E-F). To visualize the ultrastructural intermediates of MEV fusion, we performed negative-stain transmission electron microscopy following incubation of MEVs with plasma membrane-mimicking LUVs (Fig. 1G). Distinct morphological stages of membrane fusion were observed, including initial membrane contact, hemifusion, and complete fusion. Quantitative analysis after 40 minutes of incubation revealed that approximately 70% of MEVs were at the initial contact stage, ∼45% exhibited hemi fusion, and ∼30% had progressed to full fusion with the LUVs (Fig. S3C). Collectively, these results indicate that spontaneous fusion of the MEVs with the plasma membrane, represents a key initial mode of interaction at the host-MEV interface.

### MEV Fusion Increases the Host Plasma Membrane Order

To assess how MEV fusion impacts membrane mechanics, we first employed fluorescence lifetime imaging microscopy (FLIM) using Flipper-TR, a molecular rotor sensitive to changes in membrane order (lipid packing) and widely used to indirectly report apparent tension ^29–32^. THP-1-derived macrophages were treated with MEVs and labelled with 100 nM Flipper-TR, and fluorescence lifetime was monitored over 40 minutes. A marked increase in probe lifetime from 4.5 ns to 4.8 ns was observed following MEV treatment, indicating a significant elevation in plasma membrane order associated with fusion mediated reorganization (Fig. 1I,J). To validate these findings in a controlled minimal system, we used plasma membrane-mimicking GUVs doped with Flipper-TR. Consistent with the in vivo observations, MEV fusion led to an increase in lifetime from 4.6 ns to 5.0 ns (Fig. 1K,L). Furthermore, to understand whether the observed effect is due to composition, membrane tension or both, we fused DOPC LUVs (similar in size to MEVs) with homogeneous DOPC GUVs, this fusion increased the Flipper-TR lifetime, which, in this homogeneous system, indicates a reduction in membrane tension, consistent with previous reports ^33^. In striking contrast, in heterogeneous system (i.e, DOPC LUVs fusion with phase separated GUVs), a lifetime increase indicates an elevation in membrane tension, not a reduction ^33^(Fig. S1D-E). Indeed, the fusion of smaller MEVs with the GUVs transfers their intrinsic outer-leaflet asymmetry and induces outer-leaflet expansion of the GUV driving increased spontaneous tension as demonstrated previously^34–36^. To compare the magnitude of MEV-induced membrane remodeling, we compared it against the effect of MβCD-mediated cholesterol depletion, which yielded a 1 ns shift in Flipper-TR lifetime (Fig. S3D-E), in contrast 0.3-0.5 ns change observed for MEV fusion. Regulation of cell membrane mechanical parameters is essential for numerous cellular processes, including endocytosis, signalling, and vesicular trafficking^32,37–39^. To assess the effect of altered membrane order on the diffusion behaviour of lipids, we monitored the fluorescence recovery after photobleaching (FRAP) of plasma membrane-mimicking giant unilamellar vesicles (GUVs) labelled with Atto-PE. GUVs were immobilized and incubated with MEVs for 40 minutes to allow fusion to take place, and fluorescence recovery was monitored post-bleaching (Fig. S4 A,B). In comparison to host plasma membranes, MEV-fused GUVs exhibited about ∼20% lower fluorescence recovery, indicative of reduced lateral lipid diffusion. The reduction in lipid diffusion could be attributed to the nanoscale lipid phase separation accumulating line tension in the heterogenous environment of phagosome mimicking membrane ^40^. To verify this, we performed FRAP on homogeneous DOPC GUVs and phase-separated PC:SM:CHOL GUVs. Homogeneous GUVs showed rapid, near-complete recovery consistent with free lateral diffusion. Phase-separated GUVs showed significantly reduced recovery despite having higher Flipper-TR lifetimes, confirming that a substantial fraction of lipids become confined within slowly exchanging surrounding ordered (Lo) domains (Fig. S4 C,D).

Furthermore, we showed that heterogenous phase-separated membranes have inherently higher baseline Flipper-TR lifetimes indicating higher membrane order than homogeneous DOPC membranes (Fig. S1F,G). Additionally, Laurdan two-photon FLIM in MEV-treated macrophages confirms increased lipid packing/order (increased Laurdan lifetime) compared to the control (PBS treated) macrophages (Fig. 1M,N).

These findings suggest that MEV fusion leads to tighter lipid packing and phase separation within the membrane, thereby impairing lipid mobility and altering membrane order. These results collectively demonstrate that MEV incorporation into the membrane leads to a consistent increase in membrane order, both in live cells and in reconstituted membranes.

To further dissect the mechanical consequences of MEV fusion on host membranes, we evaluated how MEV incorporation alters the membrane’s compressibility modulus by employing Langmuir lipid monolayers. This approach enables precise measurement of the surface pressure (π) versus molecular area (A) isotherms, which reflect the thermodynamics of lipid interaction and packing density ^41,42^. Plasma membrane mimicking monolayers were exposed to increasing concentrations of MEVs, and the resulting π - A isotherms were measured. We observed a significant rightward shift in the isotherms with MEV addition (Fig. S4E), suggesting that MEV integrates into the monolayer, thereby increasing the average molecular area and expanding the film.

To quantitatively assess membrane compressibility, we calculated the compressibility modulus (Cs⁻¹) from the π-A isotherms using the equation:

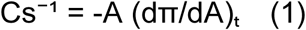

This modulus reflects the membrane’s resistance to lateral deformation. Importantly, we focused our analysis at a surface pressure range of ∼30-35 mN/m, which closely mimics the lateral pressure found in biological bilayers, where lipid packing, hydrophobic free energy, and area per molecule resemble native membranes^41,42^. At this bilayer-equivalent pressure, we found that the compressibility modulus decreased significantly with increasing MEV concentration (Fig. S4F), indicating that MEV fusion renders the membrane more laterally compressible. In addition, to compare the magnitude of MEV-induced changes in compressibility modulus, we compared it against addition of 10 mol% cholesterol, which yielded similar change in compressibility modulus corresponding to 5 mN/m (Fig. S4H). Next, as demonstrated by the in vitro data that showed higher membrane rigidity and increased compressibility of the plasma membrane post-fusion with MEVs, we explored the effect of fusion on the cell cortical stiffness using atomic force spectroscopy. In line with in vitro assays, fusion of MEVs with PMA-differentiated THP-1 macrophages cells lead to increased elastic modulus indicative of a stiffer plasma membrane (Fig. 1H, S4G). Collectively the findings suggest a complex remodelling of the host plasma membrane in which MEV fusion enhances membrane order and stiffness in the bilayer.

### MEV Fusion Triggers Actin Fragmentation and Increases Their Phagocytosis

To confirm the enrichment of the MEVs in the phagosome either in fused or engulfed state, PMA-differentiated THP-1 macrophages were treated with Anti-LAM antibody to detect the surface markers (i.e., LAM) on the MEV^43^. This revealed that the MEVs were efficiently internalized and fully phagocytosed within 2 hours. (Fig. 2A-B). The increase in phagocytosis of MEVs was also reflected in the rise in surface expression of TLR2, as detected by flow cytometry and in line with previous observation ^22^ (Fig. S5C,D). To further investigate the mode of MEV internalisation, we performed pharmacological inhibition experiments using methyl-β-cyclodextrin (MβCD) and cytochalasin D. MβCD disrupts cholesterol-rich membrane domains and impairs raft-dependent endocytosis, whereas cytochalasin D inhibits actin polymerisation and consequently blocks phagocytic uptake. Treatment with either inhibitor significantly reduced MEV internalisation compared with untreated controls. Notably, cytochalasin D produced a pronounced reduction in MEV uptake, indicating a strong dependence on actin-mediated phagocytic processes. These findings suggest that MEV internalisation occurs predominantly through phagocytosis and/or actin-dependent endocytic pathways depending on the number of MEVs engulfed (Fig. S5A-B). To determine whether MEVs reside within membrane-bound intracellular compartments, we performed immunofluorescence staining for EEA1, a well-established marker of early endosomes. Our confocal microscopy analysis revealed significant colocalization of internalized MEVs with EEA1-positive compartments, indicating that MEVs are trafficked through membrane-bound endocytic vesicles following cellular uptake (Fig. S8D). We then questioned whether the the MEV internalization and fusion impacts the host cytoskeleton dynamics and their interaction with membranes to regulate the phagocytic kinetics? To investigate this, we monitored the actin network in the MEV treated macrophages. We observed an increased abundance of small actin puncta over time suggesting actin fragmentation (Fig. 2C-D). The smaller fragments of actin indicate multiple nucleating actin facilitating high rate of actin turnover essential for phagocytosis. Concomitant with actin puncta generation, quantification of actin anisotropy demonstrated that actin is more misaligned within the actin puncta observed within the cell (Fig. 2E). Anisotropic architecture is known to directly influence membrane bending ^44^.

**Figure 2.**
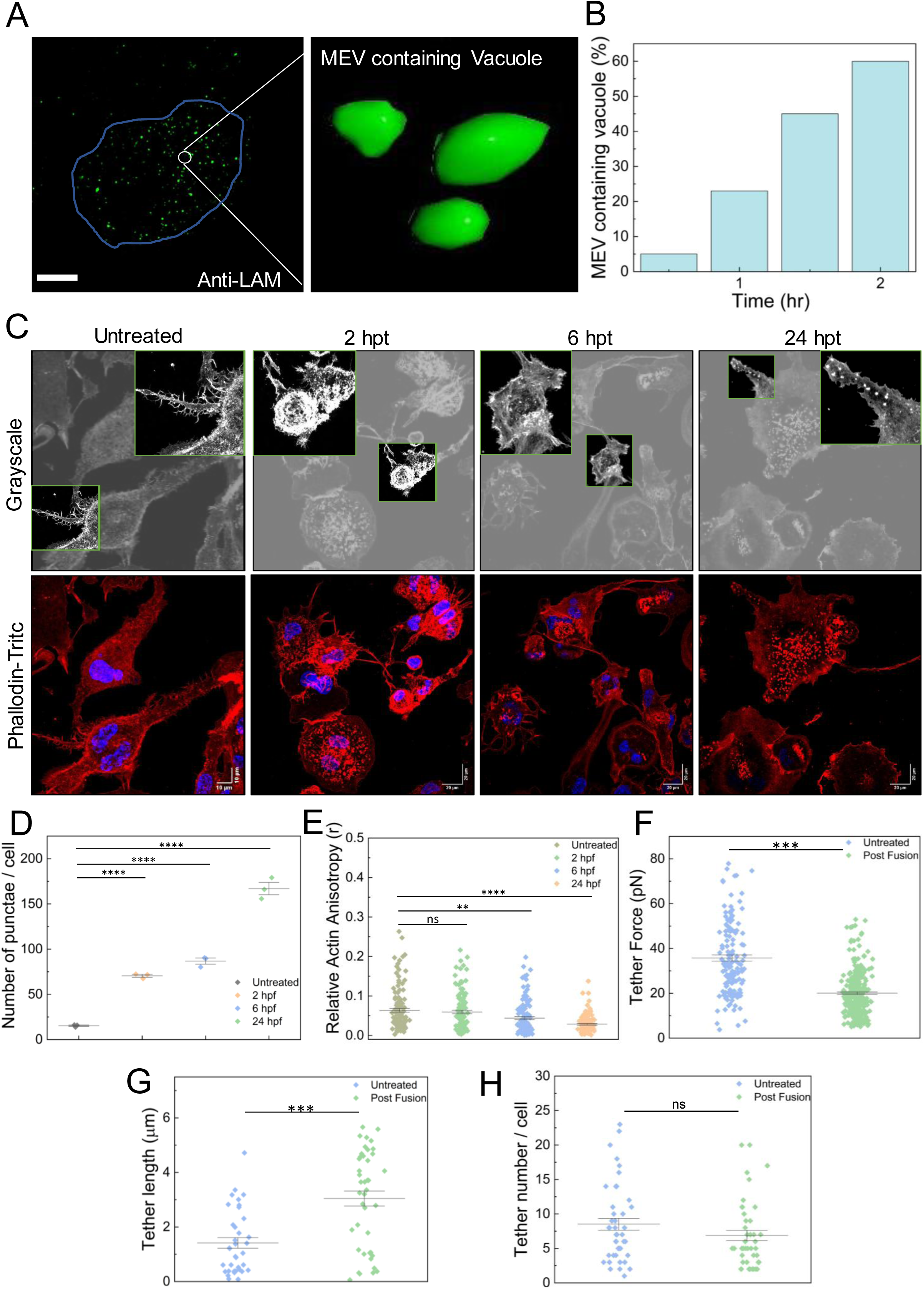
Temporal mapping of MEV uptake, cytoskeletal remodelling, and alterations in membrane–cortex mechanics. **(a)** Representative confocal micrographs showing internalized MEVs at indicated time point, detected by immunofluorescence staining using an anti-LAM antibody. **(b)** Quantification of the percentage of internalized MEVs over time, demonstrating progressive uptake by macrophages. Data points represent three independent experiments. **(c)** Representative confocal micrographs of macrophages at indicated time points following MEV treatment, showing progressive accumulation of punctate structures (red). Nuclei are stained with DAPI (blue). **(d)** Quantification of puncta per cell demonstrates a time-dependent increase upon MEV addition compared to untreated (only PBS) controls**. (e)** Quantitative analysis of actin anisotropy reveals misaligned actin network organisation and cytoskeletal remodelling following MEV fusion relative to control cells (only PBS treated). Scale bars, 20 µm. Data are presented as mean ± SEM; statistical significance determined by one-way ANOVA. **(f)** Atomic force microscopy-based tether force measurements indicate altered membrane-cytoskeleton coupling in MEV-treated cells, reflected by changes in the force required to extract membrane tethers**. (g-h)** Quantification of tether length and tether number further demonstrates changes in membrane deformability and cortical tension upon MEV interaction. Data represent mean ± SEM from ∼60 cells per condition. (**p<0.01, ***p<0.001 in one-way ANOVA).

Confocal and 3D iso-surface imaging revealed a time-dependent reorganisation of the actin cytoskeleton in macrophages following treatment of MEV. Under control conditions, cells displayed well-defined cortical actin and elongated filopodia, indicative of a stable, spread morphology (Fig. S6A). At 2 h post treatment, actin condensation at the cell cortex becomes apparent, accompanied by a noticeable reduction in cell spreading. Quantitative analysis showed a significant increase in the volume of actin per cell area at 2 h post treatment which reduced to the level of untreated (PBS-only control) cells at later time points (Fig. S6B). This suggests only a transient change in actin volume upon MEV treatment with dTHP-1 cells. Coupled with actin punctate generation, which otherwise is expected to decrease the elastic modulus, the data indicate that MEV treatment rigidifies the cell cortex (Fig. 1H). Next, we analysed tether dynamics post MEV treatment. Tethers report alterations in the membrane reservoir and curvature. While, tether force is associated with membrane stiffness, bending rigidity, and cortical interactions, tether length and number refer to changes in membrane resistance to bending. Upon MEV treatment, a decrease in tether force was observed (Fig. 2F, S6C), indicating facile pulling of tethers and hence a weaker membrane–cytoskeleton coupling. The fragmentation of actin filaments into puncta possibly remodels the interactions between the plasma membrane and the actin network. The low tether force enables high tether extension, reflected in increased tether length in dTHP-1 upon fusion (Fig. 2G). Concomitantly, as the total membrane reservoir remains the same, flow of membrane lipids to reach higher tether lengths restricts the number of tethers per cell to be formed and is observed as reduced tether number (Fig. 2H, S6D). Together, MEVs may trigger TLR2-mediated phagocytic activation, further facilitated by actin fragmentation, potentially initiating downstream phagosome formation and maturation processes.

### MEVs Interfere With The Host Phagosome Maturation Pathway By Enriching Their MEV-Derived Lipids

We next investigated the impact of MEVs on phagosome maturation and to this end we tracked the recruitment dynamics of the early endosomal marker Rab5. THP-1-derived macrophages were transfected with mCherry-tagged Rab5 (mCh-Rab5), which allowed real-time visualization of early endosome association through single-cell time-lapse imaging. Upon treatment with MEVs derived from *M. smegmatis* and *M. tuberculosis* H37Ra labelled with PKH67, Rab5 colocalization with the phagosomal membrane was significantly delayed, first appearing around 8 and 9hours post-treatment, respectively (Fig. 3B-C, Supplementary movie 4-7). This was further confirmed through maximum intensity projection images taken at end time points, which revealed Rab5 accumulation around MEV-containing phagosomes (Fig. 3B-C). In contrast, macrophages treated with *E. coli*-derived EVs showed early recruitment of Rab5 to the phagosomal membrane, with colocalization detectable as early as 3 hours post-internalization. This was visible in the maximum intensity projection images, which demonstrated robust Rab5 signal surrounding the EV-containing compartments at this earlier time point (Fig. 3A, Supplementary movie 2-3). A similar delay in Rab5 recruitment was also quantitatively evident in fixed-cell imaging of macrophages expressing mCh-Rab5 at various time points post-MEV treatment revealed by pearson’s correlation coefficient, corroborating the live-cell imaging results (Fig. S7A-B). These observations suggest that MEVs delay the recruitment of Rab5 to the maturing phagosome, potentially through spontaneous fusion with the phagosomal membrane that disrupts normal trafficking and maturation.

**Figure 3.**
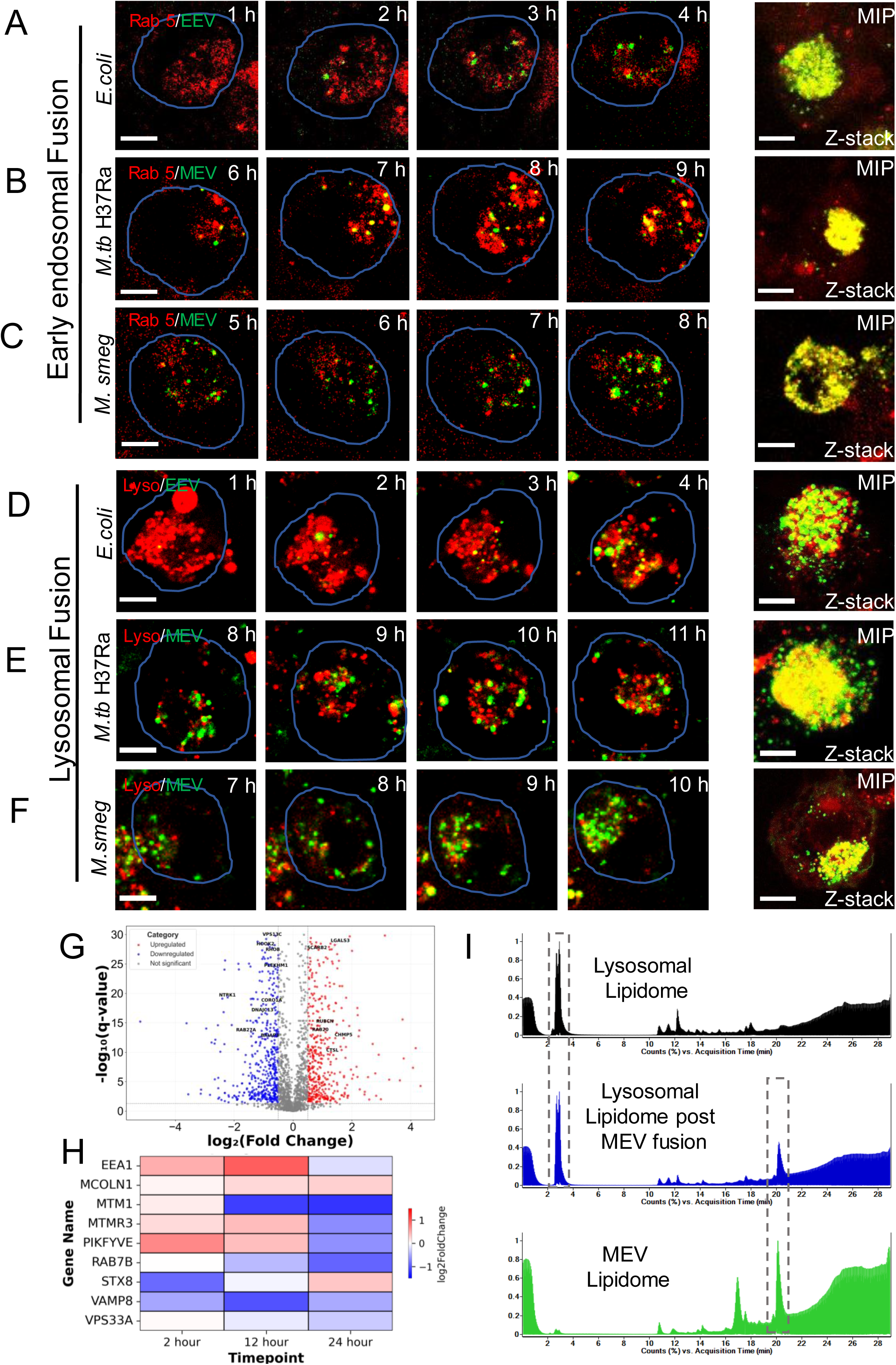
MEV fusion delays the kinetics of phagosome maturation. Live-cell time-lapse confocal imaging of early endosome-phagosome fusion following internalisation of different bacterial extracellular vesicles (Green). **(a)** *Escherichia coli* extracellular vesicles (EEVs), **(b)** *Mycobacterium tuberculosis* H37Ra extracellular vesicles (MEVs), and **(c)***Mycobacterium smegmatis* extracellular vesicles (MEVs) were internalised by macrophages stably expressing Rab5 an early endosomal marker (Red). Sequential imaging reveals dynamic fusion of Rab5-positive early endosomes with the EV-containing phagosomes. Corresponding maximum-intensity projections were acquired at the final time point from nearby cells to minimise photobleaching. The time interval between frames was 60 minutes. Scale bar, 10 µm. Each condition represents ∼70 cells. Live-cell time-lapse confocal imaging of lysosome-phagosome fusion following early endosomal fusion events. **(d)** *Escherichia coli* extracellular vesicles (EEVs), **(e)** *Mycobacterium tuberculosis* H37Ra extracellular vesicles (MEVs), and **(f)** *Mycobacterium smegmatis* extracellular vesicles (MEVs) (green) were monitored for subsequent fusion with lysosomes (red). The time-series imaging demonstrates the progressive recruitment and fusion of lysosomes with EV-containing phagosomes. Maximum-intensity projections at the final time point were collected from adjacent cells to reduce photobleaching. The time interval between frames was 60 minutes. Scale bar, 10 µm. Each condition represents ∼83 cells. **(g)** Volcano plot showing differentially expressed genes in macrophages 12 hours after MEV treatment. Significantly upregulated genes (n = 377, red), downregulated genes (n = 428, blue), and unchanged genes (n = 1216, grey) are indicated. Threshold was log2FC > 0.5 (or < −0.5) and P_value_ < 0.05. **(h)** Heatmaps showing temporal gene expression profiles of macrophage genes associated with the phagosome maturation pathway at different time intervals (2,12,24 hours) following MEV treatment. **(i)** Lipidomics chromatograms comparing the lysosomal lipidome, MEV lipidome, and lysosomal lipidome following MEV fusion.

To further understand the molecular basis underlying the observed delay in phagosomal maturation induced by MEVs, we performed transcriptomic profiling of macrophages exposed to MEVs and compared them with untreated (PBS-only control) macrophages at different time points. RNA-sequencing analysis revealed striking transcriptional reprogramming with approximately ∼ 1,000 genes differentially expressed at 12 hours relative (Fig. 3G, S8A). An unbiased ranking based on log2FC revealed that the top-most upregulated genes (that showed ∼ 4-5 fold change) mostly affect cellular pathways, such as disrupting ion homeostasis, cellular stress responses, membrane lipid transport, hypoxia-dependent metabolic adaptation, and growth factor-mediated cytoskeletal remodelling (Fig S9A,10A). Also, the top most downregulated genes (that showed ∼ 3-5 fold change) are mostly affecting the cellular pathways such as immune differentiation programs, mitochondrial function, DNA replication, mitotic spindle dynamics, and cell-cycle progression, potentially contributing to a reduced proliferative and metabolically altered macrophage state (Fig . S9A,10B). Importantly, we also discovered suppression of several key regulators of endosomal maturation and fusion among the genes exhibiting significant downregulation (∼ 2-3 fold) at 12 hours (Fig. 3H). These included MTM1, a phosphoinositide phosphatase critical for endosomal homeostasis^45^; Rab7B, a small GTPase that promotes transition from early to late endosomes and lysosomal biogenesis^46,47^; VAMP8, a SNARE protein required for lysosomal fusion events^48,49^; and PIKFYVE, the lipid kinase responsible for generating PI(3,5)P₂, which regulates endolysosomal trafficking and membrane dynamics ^50^. The reduced expression of these factors indicates a coordinated manipulation of the molecular machinery essential for phagosome progression towards lysosomal fusion.

In contrast, several genes associated with early endosomal function and compensatory trafficking pathways were upregulated at 12 hours (Fig. 3H). These included EEA1, a canonical early endosome tethering protein that stabilizes PI(3)P-enriched membranes^51^; MCOLN1, a lysosomal Ca²⁺ channel that regulates vesicular trafficking and membrane fission events^52^; MTMR3, a phosphatase that counterbalances PI3P signaling^53^; STX8, a SNARE component linked to endosome-endosome fusion^54^; and VPS33A, a Sec1/Munc18 (SM) family protein that coordinates SNARE-mediated membrane fusion. Upregulation of these transcripts suggests that macrophages attempt to counteract MEV-induced trafficking defects by reinforcing early endosomal tethering and fusion machinery. Together, transcriptomic signatures are in line with the delayed Rab5 recruitment and indicate an impaired lysosome-phagosome fusion. This also reveals a dual impact of MEVs on the endolysosomal system, i.e, suppression of genes essential for phagosome maturation and lysosomal fusion, coupled with compensatory upregulation of early endosomal regulators by the host (Supporting information 2).

To further investigate the effect of MEVs on phagosome-lysosome fusion, we monitored lysosomal acquisition in THP-1 - derived macrophages treated with EVs from *E. coli*, *M. smegmatis*, and *M. tb* H37Ra. Colocalization of LysoTracker, a fluorescent probe that accumulates in acidic organelles, with PKH67-labeled EV-containing compartments was monitored over time using live-cell imaging. In macrophages treated with *E. coli* EVs, robust LysoTracker colocalization was observed as early as 4 hours post-internalization, as evident from the maximum intensity projection images (Fig. 3D, Supplementary movie 8-9). In contrast, LysoTracker signal colocalization was markedly delayed in cells treated with MEVs from *M. smegmatis* and H37Ra, appearing only around 10 and 11hours post-treatment, respectively (Fig. 3E-F, Supplementary movie 10-13). A comparable delay in phagosome-lysosome fusion was also observed in fixed-cell imaging using LysoTracker-stained macrophages at various time points following MEV treatment, further corroborating the live-cell imaging (Fig. S11).To further investigate whether the MEVs alter autophagy, we quantified the percentage of LC3B-positive cells by immunofluorescence following MEV treatment and compared with untreated cells and cells treated with bafilomycin A1, a well-established modulator of autophagic flux. We observed a significant increase in LC3B-positive cells in MEV-treated macrophages relative to untreated controls. As expected, bafilomycin A1 treatment also resulted in robust LC3B accumulation (Fig S8B-C). These findings independently support the transcriptomic prediction that autophagy-related pathways are altered following MEV exposure. Additionally, fixed-cell imaging of THP-1 macrophages infected with *E. coli*, live GFP–*M. smegmatis*, or heat-killed *M. smegmatis*, shows lysosomal colocalization at 4 h, 48 h, and 30 h, respectively, with the pronounced delay in live *M. smegmatis* highlighting MEV-mediated impairment of phagosome–lysosome fusion (Fig. S12). These observations suggest that EVs from mycobacteria significantly delay phagosome acidification and fusion with lysosomes, further supporting their role in disrupting normal phagosome maturation pathways.

This was further corroborated by visualizing the fusion dynamics where the macrophages were preloaded with fluorescent-dextran (red) to label lysosomes and subsequently challenged with FITC-latex beads (green) to generate classical phagosomes (Fig 4A-B). In PBS-treated cells, bead-containing phagosomes progressively acquired the lysosomal dextran signal over time, resulting in extensive colocalization by 120 minutes (Fig 4A and inset). This observation is consistent with normal phagosome maturation and efficient phagolysosome formation. In contrast, in MEV-treated cells, bead-containing phagosomes remained largely devoid of lysosomal dextran throughout the imaging period, exhibiting minimal colocalization even at the final time point (Fig 4B and inset). Thus, MEV exposure markedly impaired the transfer of lysosomal contents to phagosomes, indicating a defect in phagosome-lysosome fusion. Next we wondered whether the MEVs directly interact and remodel existing bacteria containing phagosome. To this end, we followed the intracellular fate of internalized MEVs (PKH-26 labelled, red) within infected macrophages that already contained bacteria-laden phagosomes (GFP *M.smeg,* green). Remarkably, MEVs consistently gathered around and wrapped these pre-existing phagosomes, pointing to a direct physical engagement of the target membrane (Fig 4C, inset from multiple plains), enabling them to actively remodel phagosomal membrane architecture and physicochemical parameters. Importantly, to test the biological relevance of MEV in facilitating bacterial infection, BALB/c mice were pre-treated with MEVs and, 48 hours later, challenged intranasally with GFP-expressing *Mycobacterium smegmatis* (Fig 4D). Lungs were collected on day 9 post-infection, and bacterial burden was assessed by quantifying GFP fluorescence. Strikingly, MEV-pretreated mice exhibited significantly higher GFP signals in the lungs compared with PBS-treated controls, indicating enhanced bacterial persistence following MEV exposure. These in vivo findings provide independent functional evidence that MEV-mediated host cell remodelling creates an intracellular environment that is more permissive to mycobacterial survival. This is in line with a previous report that showed that the spreading of a mycobacterial cell-surface lipid into host epithelial membranes promotes infectivity^55^ .

**Figure 4.**
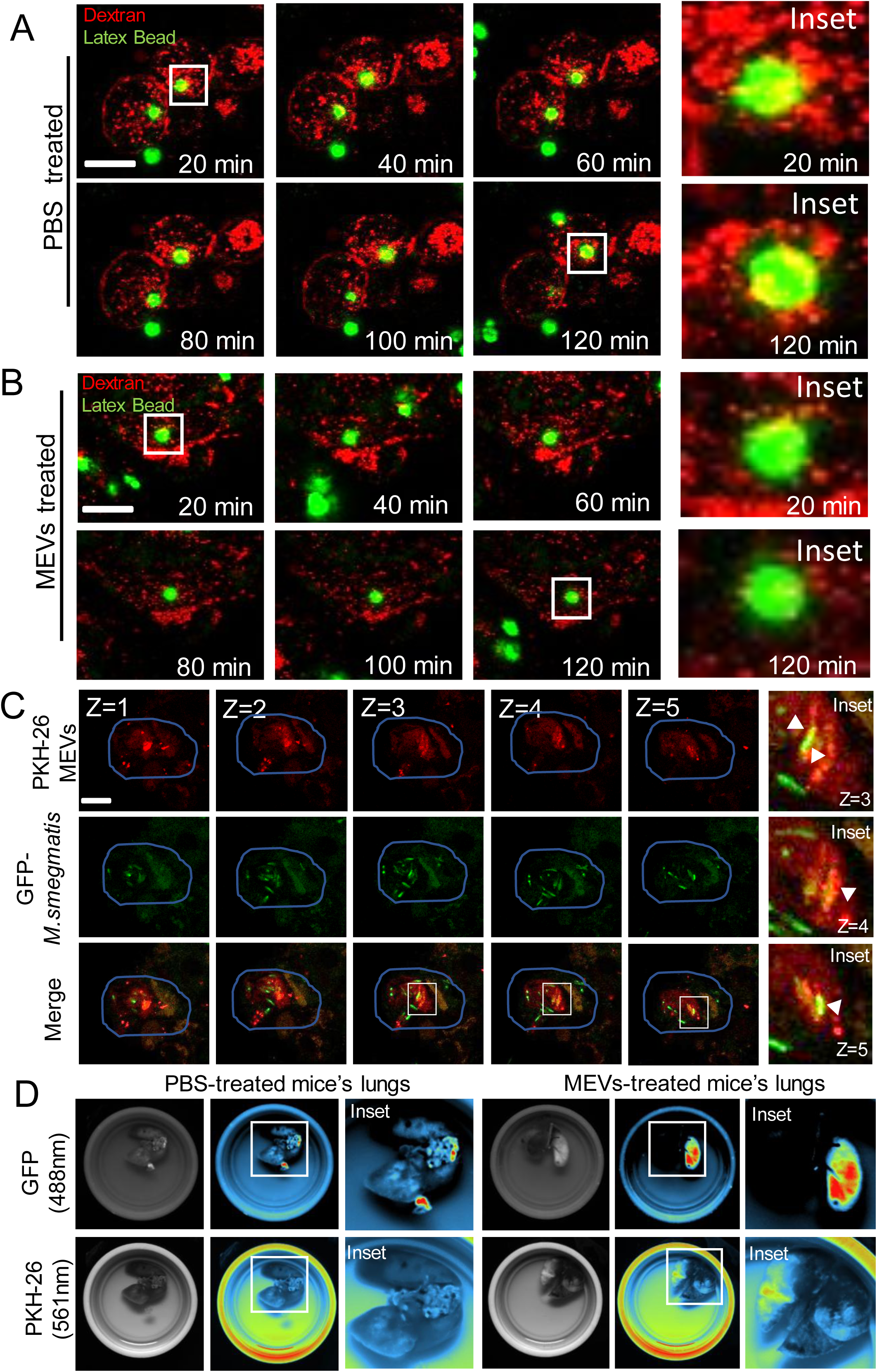
MEV-Mediated Remodelling of Incoming and Established Phagosomes Facilitates Bacterial Persistence In Vivo. Live-cell time-lapse confocal imaging of early phagosome-lysosome fusion following internalisation of different latex beads (Green) in the **(a)** PBS-treated and **(b)** presence of MEVs. Sequential imaging reveals dynamic fusion of phagosomes with the dextran-loaded lysosomes (red). The time interval between frames was 10 minutes. Scale bar, 10 μm. Each condition represents ∼50 cells**. (c)** Live-cell confocal Z-stack imaging of preformed bacteria-containing phagosomes (Green) and internalised MEVs (Red) fusion. Imaging reveals that the internalised MEVs remodelled the existing phagosomal membrane. Scale bar, 10 μm. These are representative images from three independent experiments. (n∼65 cells). **(d)** Representative images of lungs isolated from PBS-treated and MEV-treated BALB/c mice showing the fluorescence intensity of GFP-expressing M. smegmatis. The fluorescence signal reflects bacterial persistence within the lung tissue, with differences in intensity indicating altered bacterial burden following MEV treatment.

We hypothesized that the enrichment of the MEV lipid components in the phago-lysosomal membrane compartments could be responsible for the observed delay in fusion. The identification of specific lipidic changes will enable better rationalization of the changes induced in host membranes upon fusion. The lipidomic profiling of purified lysosomes, MEVs and purified lysosomes from MEV-treated macrophages was performed by extracting lipids followed by mass spectroscopy optimized for mycobacterial lipids as previously described^56–59^. Firstly, mycobacterial lipids identified from the MEV lipids were classified into six major lipid classes based on MtbLipidDB (i.e, glycerophospholipids, saccharolipids, glycerolipids, fatty acyls, prenols and polyketides) and in decreasing levels of abundance (Fig. 3I, Supporting information 1).

In glycerophospholipids, the predominant lipids identified were acylated versions of phosphatidylinositol mannosides (PIMs) such as AC1PIM1, AC1PIM2, AC2PIM2 and AC2PIM3 (Fig. S13A-B). These lipids are highly specific to mycobacterial species and their high abundance is mainly attributed to them being the primary membrane-forming lipids. Notably, the enrichment of ACPIMs in the inner plasma membrane leaflet implies that other membrane-derived heterogenous MEV sub-populations may exist, a possibility that merits further exploration. However, it is important to note that ACPIMs are also found to enrich in the outermembrane of the cell envelope during the late stages of secretion ^58^. Other abundant lipids within glycerophospholipids were cardiolipins (CL), lysophosphatidylinositol (LPI), and standard phospholipids such as PA, PG, PI and PE. Within saccharolipids, the major contributors were glycopeptidolipids I and IIB among other lipid as shown in Fig S13B. Saccharolipids are generally known to facilitate infection and foster evasion of phagocytosis and promote membrane permeability. Mycolic fatty acids are long-chain (C60-90) fatty acids specific to mycobacterial strains and were identified in the fatty acyl lipid class (Fig. S13B) and among them, mycosanoic acid was the most abundant. We could detect both alpha and methoxy mycolic acids within the bacterial lipidome, albeit with low abundances. Next, mannosyl phosphomycoketides, and PDIM B constituted the major components within prenol and polyketides class (Fig. S13B), though their absolute abundance was low. These lipids are known to act as carrier lipids. Lipids such as mono-, di-, and tri-acyl glycerol (MAG, TAG and DAG) constituted the glycerolipids but are not selective to mycobacterial lipidome as these are also found in mammalian lipidome (Supporting information 1), and they mainly act as storage lipids. After quantifying MEV lipidome, we specifically looked for signature of mycobacterial lipids within lysosomes purified from MEV infected host cells as a result of MEV fusion. Class-wise lipid composition of untreated (PBS-only control) lysosomes is used as the baseline for comparison with MEV-fused lysosomes (Fig. S13C). As demonstrated in (Fig. S13D), we could detect all major lipids (class wise) within the fused fraction confirming membrane fusion as well as insertion of bacterial lipids within the host’s lysosomal membrane. Based on these, we investigated changes within the host cellular lysosomal lipidome post integration of specific bacterial lipids which could alter membrane properties and thus influence the rate of fusion. We evaluated two main factors: alkyl chain length and degree of unsaturation. Upon fusion with MEVs, the average chain length of lipids within lysosomal membrane increased (Fig. S13E) and is mainly attributed to the integration of long-chained bacterial lipids. The dynamics of the increased chain length owing to their defective packing may contribute to free void volume resulting in disorderliness of the overall membrane. The observed decrease in degree of unsaturation (> 4 double bonds) upon fusion reflects a local ordering in the membrane (Fig. S13F). Together, we think that the fusion of MEVs though might order the membrane locally, however, hydrophobic mismatch between lysosomal and MEV lipids would induce lipid segregation leading to overall disorderliness. It is important to note that the lipidomic profiling provides a semi-quantitative indication of lipid remodelling rather than absolute molar concentrations or direct comparisons across lipid classes.

### MEV-derived Lipids Are Sufficient to Disrupt Lysosomal Homeostasis in Host Macrophages

Having established that MEVs fully fuse with the lysosomal compartment, we next sought to determine whether they perturb lysosomal homeostasis and which components can minimally drive the same. Alterations in key membrane biophysical properties - such as polarity, viscosity, order and tension - are strongly associated with the progression of various pathological states ^60–62^. Many intracellular biochemical processes are highly sensitive to local viscosity, changes in organelle-specific viscosity can serve as early indicators of cellular dysfunction ^63^. To assess lysosomal viscosity, we employed JIND-Mor, a molecular rotor-based fluorescent probe that selectively accumulates in lysosomes and reports changes in viscosity through alterations in its fluorescence lifetime and highly pH tolerant ^64^. Differentiated THP-1 macrophages were incubated with 0.2 µM JIND-Mor for 15 minutes before imaging, and lysosomal viscosity was monitored 12 hours post-treatment with EVs derived from *M. smegmatis* and *M. tuberculosis* H37Ra. This time point was selected based on previously observed trafficking and colocalization time of MEVs with lysosomes (Fig. 3E-F), ensuring that viscosity measurements reflect MEV post-fusion lysosomal dynamics. We observed a significant increase in the fluorescence lifetime of JIND-Mor in macrophages treated with MEVs. Specifically, the average lifetime measured in cells treated with EVs from *M. smegmatis* was approximately ∼3.0 ns, while EVs from *M. tuberculosis* H37Ra yielded a slightly higher lifetime of ∼3.1 ns (Fig. 5B,C, E). In contrast, untreated (PBS-only control) macrophages exhibited a baseline JIND-Mor fluorescence lifetime of ∼2.75 ns (Fig. 5A,E). This increase in fluorescence lifetime indicates an elevation in lysosomal viscosity following MEV treatment. Since JIND-Mor lifetime positively correlates with microviscosity, these results suggest that fusion of MEVs with the lysosomal membrane leads to a more viscous intra-lysosomal environment, potentially reflecting altered lipid packing or changes in luminal content dynamics. Interestingly, to delineate the specific contribution of MEV lipids to lysosomal viscosity changes, we reconstituted synthetic vesicles composed solely of lipids extracted from MEVs (hereafter referred to as MDLVs, MEV-derived lipid vesicles), maintaining a size distribution comparable to native MEVs. Upon treating macrophages with these MDLVs and subsequently monitoring lysosomal viscosity using JIND-Mor, we observed a striking increase in fluorescence lifetime, reaching approximately ∼3.4 ns (Fig. 5D-E). This enhancement exceeds that observed with intact MEVs, indicating that the lipid components alone are sufficient to induce significant alterations in lysosomal microviscosity. Specificity was confirmed by *E. coli* EV treatment, which despite identical uptake did not replicate the JIND-Mor lifetime increase seen with MEVs, ruling out non-specific vesicle internalisation effects (Fig S14A-B). Further, to illustrate the magnitude of changes in lysosomal viscosity, cholesterol was selectively trafficked to lysosomes. Cholesterol-loaded lysosomes exhibited a significant increase in viscosity compared to controls as a result of its established role in enhancing membrane packing/order ^65^. This control therefore provides a benchmark for interpreting the viscosity changes observed following MEV treatment (Fig S14C-D).

**Figure 5.**
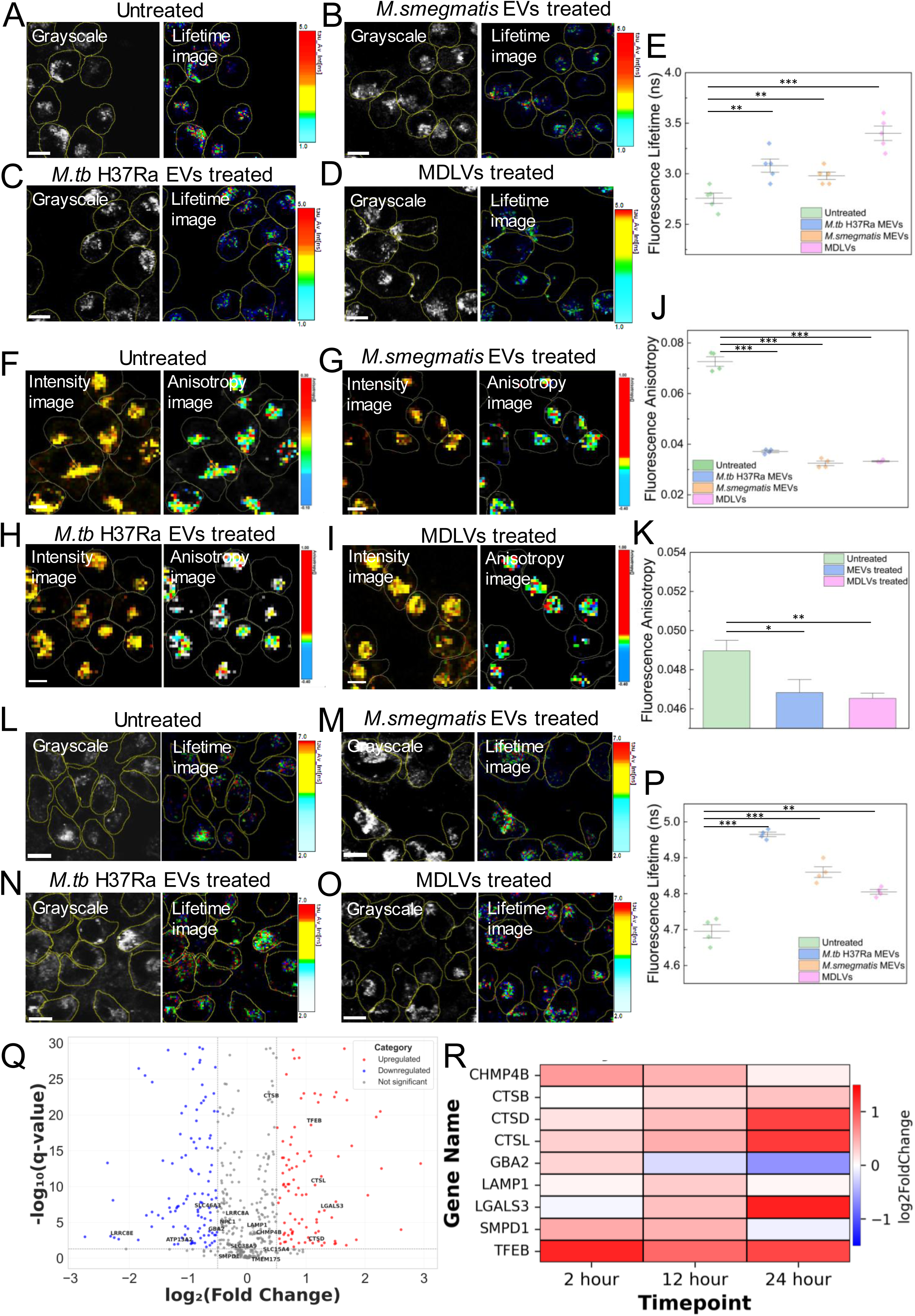
MEVs and MDLVs mediated disruption of lysosomal homeostasis. Fluorescence lifetime imaging microscopy (FLIM) analysis of lysosomal viscosity in macrophages treated with different MEVs. **(a)** Untreated (only PBS) macrophages, **(b)** *Mycobacterium smegmatis* EV–treated macrophages, **(c)** *Mycobacterium tuberculosis* H37Ra EV-treated macrophages, and **(d)** MDLV-treated macrophages stained with Jind-Mor, a lysosomal viscosity sensor. All measurements were acquired 12 hours post-treatment, revealing MEV-dependent alterations consistent with lysosomal fusion. Images are representative of five independent biological replicates (∼400 cells). **(e)** Quantification of fluorescence lifetimes corresponding to the conditions shown in **(a-d).** Each data point represents the mean lifetime from an independent biological replicate. (*p* < 0.05, one-way ANOVA). FLIM analysis of lysosomal membrane anisotropy in macrophages treated with different MEVs. **(f)** Untreated (only PBS) macrophages, **(g)** *Mycobacterium smegmatis* EV-treated macrophages, **(h)** *Mycobacterium tuberculosis* H37Ra EV-treated macrophages, and **(i)** MDLV-treated macrophages stained with Jind-Mor. All measurements were acquired 12 hours post-treatment, revealing MEV-dependent alterations consistent with lysosomal fusion. Images are representative of five independent biological replicates (∼400 cells). **(j)** Quantification of fluorescence lifetimes corresponding to the conditions shown in **(f-i).** Each data point represents the mean lifetime from an independent biological replicate. (*p* < 0.05, one-way ANOVA). **(k)** Fluorescence anisotropy measurements of isolated and purified lysosomes under different treatment conditions, untreated macrophages, *Mycobacterium tuberculosis* H37Ra EV-treated macrophages, and MDLV-treated macrophages using 1 µM DPH as a membrane-order probe. All measurements were conducted in PBS (pH 7.4) at 37 °C. The data represent three independent biological replicates. FLIM analysis of lysosomal membrane tension in macrophages treated with different MEVs. **(l)** Untreated (only PBS) macrophages, **(m)** *Mycobacterium smegmatis* EV-treated macrophages, **(n)** *Mycobacterium tuberculosis* H37Ra EV-treated macrophages, and **(o)** MDLV-treated macrophages stained with Lyso-Flipper, a lysosomal membrane tension sensor. All measurements were acquired 12 hours post-treatment, revealing MEV-dependent alterations consistent with lysosomal fusion. Images are representative of five independent biological replicates (∼475 cells). **(p)** Quantification of fluorescence lifetimes corresponding to the conditions shown in **(l-o).** Each data point represents the mean lifetime from an independent biological replicate. (*p* < 0.05, one-way ANOVA). **(q)** Volcano plot showing differentially expressed genes in macrophages 24 hours after MEV treatment. Significantly upregulated genes (n = 90, red), downregulated genes (n = 106, blue), and unchanged genes (n = 282, grey) are indicated. Threshold was log2FC > 0.5 (or < −0.5) and P_value_ < 0.05. **(r)** Heatmaps showing temporal gene expression profiles of macrophage genes associated with the Lysosomal stress at different time intervals (2,12, 24 hours) following MEV treatment. (**p<0.01, ***p<0.001 in one-way ANOVA).

Lysosomal membrane order (or fluidity) is a critical factor in maintaining lysosomal homeostasis and function ^66^. To assess how MEVs influence lysosomal membrane dynamics, we quantified fluorescence anisotropy of JIND-Mor localized to the lysosomal membrane, extrapolating data from Fig. 5A-E. Compared to untreated (PBS-only control) macrophages, which displayed a baseline anisotropy of ∼0.073 (Fig. 5F, J), MEV-treated cells exhibited a marked reduction in lysosomal membrane anisotropy. Specifically, macrophages treated with MEVs from *M. smegmatis* showed an average anisotropy of ∼0.032, while treatment with MEVs from *M. tuberculosis* H37Ra yielded a slightly higher value of ∼0.038 (Fig. 5G-H, J). These results indicate a significant membrane disorderliness upon MEV fusion. Consistent with this, macrophages treated with MDLVs composed solely of MEV-derived lipids also demonstrated a decreased anisotropy of ∼0.034 (Fig. 5I-J). To confirm this further, we isolated and purified the lysosomes as described ^66^ from MEV-treated and MDLV-treated macrophages and found the fluorescence anisotropy of DPH to decrease by ∼ 4% (Fig. 5K). Lipid membrane packing and tension are essential for fusion-fission dynamics and maintenance of organelle integrity, thereby playing a pivotal role in cellular homeostasis^34–37^. To investigate whether MEVs modulate lysosomal membrane order, we employed the lysosome-targeted mechanosensitive probe Lyso-Flipper. Untreated (PBS-only control) macrophages exhibited a baseline Lyso-Flipper fluorescence lifetime of ∼4.7 ns (Fig. 5L). Upon treatment with MEVs derived from *M. smegmatis*, the lifetime increased to ∼4.85 ns, while MEVs from *M. tuberculosis* H37Ra produced a similar increase to ∼4.98 ns. Notably, MDLVs also elevated the lysosomal membrane order, with a lifetime of ∼4.8 ns (Fig. 5 L-P). To further strengthen the analysis, EEV-treated cells did not exhibit a comparable Lyso-Flipper lifetime increase despite vesicle uptake, confirming the response is specific to pathogenic bacterial vesicles and their unique lipid composition, not a general effect of internalisation (Fig S14E-F). This suggests that MEVs and specifically their lipid constituents are sufficient to induce measurable increases in lysosomal membrane order, potentially contributing to altered lysosomal mechanics and disrupted homeostasis.

We next asked whether these changes translate into transcriptional reprogramming of host macrophages. To address this, we performed RNA sequencing of untreated (PBS-only control) and MEV-treated macrophages at different time intervals. At the 12-hour post treatment, MEV exposure induced a broad transcriptional response with approximately 300 genes differentially expressed compared to untreated (PBS-only control) macrophages (Fig. 5Q). Pathway enrichment analysis of these early transcriptional changes revealed significant enrichment in categories associated with lysosomal stress, lysosomal membrane damage and lipid metabolic processes, consistent with the cellular response to altered membrane homeostasis (Fig. 5Q). However, 24 hours post-treatment, the transcriptional profile consolidated into a focused set of genes strongly linked to lysosomal function and stress signaling. We observed marked upregulation of cathepsins (*CTSD*, *CTSL*, *CTSB*), reflecting an attempt to enhance proteolytic activity in response to impaired lysosomal degradation (Fig. 5R). In addition, *LGALS3* (Galectin-3), a known sensor of lysosomal membrane damage ^67^, was significantly upregulated, indicating the engagement of damage surveillance pathways. Importantly, *TFEB* the master regulator of lysosomal biogenesis and autophagy^68^ was also upregulated, suggesting the initiation of a compensatory transcriptional program to restore lysosomal capacity (Fig. 5R).

Interestingly, several genes central to lysosomal integrity and lipid homeostasis were downregulated (Fig. 5R). These included *GBA2* (β-glucosidase 2), a key enzyme in glycosphingolipid metabolism ^69^; *CHMP4B*, a component of the ESCRT-III complex essential for membrane repair and intraluminal vesicle formation ^32^; and *SMPD1*, encoding acid sphingomyelinase, which regulates membrane curvature and lipid turnover ^70^. The suppression of these genes suggests that MEV-derived lipids may impair the cell’s ability to restore lysosomal stability and lipid balance, thereby compounding the mechanical stress imposed at the membrane level. Together, these transcriptomic findings reveal that MEV-induced alterations in lysosomal membrane mechanics are paralleled by activation of lysosomal stress-adaptive programs (cathepsins, TFEB, galectins) and simultaneous suppression of repair pathways. This transcriptional dichotomy suggests that while macrophages attempt to compensate for lysosomal dysfunction, MEV-derived lipids create a sustained imbalance that prevents effective restoration of lysosomal homeostasis (Supporting information 2).

### MEV-derived Lipids Elevate Host Phagosomal Membrane Order and Apparent Tension to Inhibit Phagosome-Lysosome Fusion

We next sought to determine whether the inhibition of phagosome–lysosome fusion correlates with the membrane remodeling induced by MEV-derived lipids. To address this, using micropipette aspiration, we quantified the change in tension of GUV membrane doped with MEV-derived lipids in comparison to GUVs mimicking native phagosomal membrane. Notably, MEV-lipid-doped membranes showed phase separation with distinct ordered and disordered domains displaying markedly higher domain mobility compared to native phagosomal membranes. During aspiration, the vesicle geometry - specifically the aspiration length (*Lp*) and vesicle radius (*Rv*) (Fig. 6A-B, Supplementary movie 14-15) was monitored to assess changes in membrane mechanics. Under a constant applied pressure (Fig. 6A-B), MEV-lipid-doped membranes exhibited a significantly greater aspiration length (*Lp*) relative to control membranes indicating increased membrane deformability of predominantly *Ld* region (Fig. 6C). In a phase separated (*Lo-Ld*) membrane *Lo* domains are more ordered resulting in a smaller area per lipid. Consequently, when a substantial fraction of the membrane undergoes *Lo-Ld* phase separation, the average area per lipid of the entire GUV is expected to decrease slightly, effectively reducing the excess membrane area available. In addition *Lo* domains due to its line tension typically protrude slightly resulting out of plane deformation and this again contributes towards the area cost and the increase in global membrane tension. We infer the increase in global tension from the aspiration parameters shown in Fig 6E. We have revised the manuscript to make this explicit. The extended aspiration length therefore reflects selective protrusion of the fluid phase, not a globally reduced tension. At the global scale the membrane tension can remain high owing to the phase separation as evident from the Fig. S15A in line with previous reports^71–74^ . Quantitative analysis revealed that MEV-lipid incorporation enhanced the area expansion by approximately ∼ 1.5-fold compared to control membranes (Fig. 6D). Similarly, the bending modulus of the host membrane decreased by nearly tenfold in the presence of MEV-derived lipids (Fig. 6E). For vesicles where *Lp* = 0, no direct area expansion occurred, preventing the determination of both area stretch modulus and bending modulus. Indeed using phase-separated model membranes as a control system that exhibited a similar increase in protrusion length under identical aspiration conditions (Fig. S15A). To further validate these observations, we performed fluorescence lifetime imaging using the mechanosensitive probe Flipper-TR (Fig. 6F-G). Phagosomal membrane mimicking GUVs and their MEV-lipid–doped counterparts were prepared with varying molar percentages of MEV lipids and labelled with Flipper-TR. Consistent with the previous observations, MEV-derived lipids induced a concentration-dependent increase in fluorescence lifetime, indicative of elevated membrane order (Fig. 6G).

**Figure 6.**
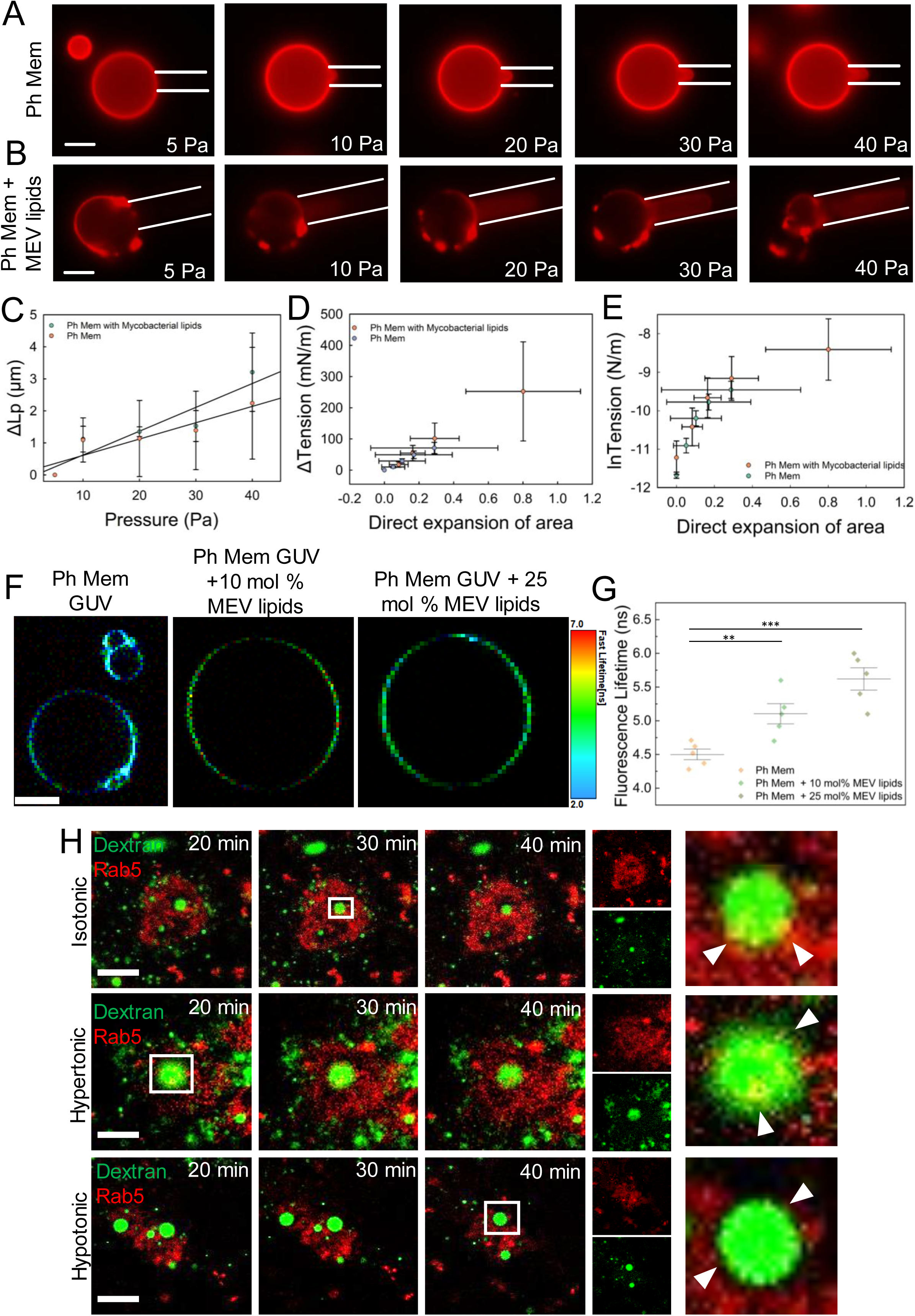
MEV-derived lipids alone are sufficient to increase the phagosome membrane tension. Time-lapse fluorescence imaging of GUVs labeled with 0.1% Rh-PE (red) during micropipette aspiration under increasing negative pressure. (**a)** Phagosomal membrane mimic and **(b)** phagosomal membrane doped with 25 mol% MEV-derived lipids. Scale bar, 5 µm. Images are representative of three independent experiments. **(c)** Plot of protrusion length (ΔL_P_) versus applied pressure (P) for conditions in panels **a** and **b**. **(d)** Quantification of the area stretch modulus (K_a_) derived from the relationship between change in membrane tension (Δτ) and direct area expansion. **(e)** Quantification of the bending modulus (K_b_) obtained from the semi-logarithmic plot of membrane tension versus area expansion for the conditions shown in panels **a** and **b**. Data represent mean ± SD from three independent experiments. **(f)** FLIM images of Flipper-Tr labelled phagosomal membrane mimic GUVs, Phagosomal membrane GUVs doped with 10mol% and 25mol% of MEV derived lipids. Scale bar, 5 µm. **(g)** Quantification of Flipper-Tr fluorescence lifetimes of the conditions as shown in **(f).** Each data point represents each biological replicate. All experiments are conducted in PBS (pH 7.4) at 37°C. **(H)** Time-lapse confocal imaging of macrophages stably expressing mCherry–Rab5 (red) and treated with 70-kDa fluorescent dextran (green) under different osmolarity conditions: isotonic, hypotonic (increased membrane tension), and hypertonic (reduced membrane tension). Images were acquired at 10-minute intervals. All other experimental parameters were kept identical across conditions. (**p<0.01, ***p<0.001 in one-way ANOVA).

Since membrane order and tension are empirically correlated via lateral area expansion, we wondered whether membrane tension directly influences Rab5 recruitment onto the phagosomal membrane. We used THP-1-derived macrophages stably expressing mCherry-tagged Rab5 (mCh-Rab5) and monitored endocytic trafficking through single-cell time-lapse imaging. Cells were pulsed with 70-kDa fluorescent dextran for 20 minutes to label newly formed endocytic vesicles, after which imaging was initiated under controlled osmotic environments (+50 mOsm and - 50 mOsm) to modulate membrane tension ^30,75^. Under isotonic conditions, where membrane tension remains close to physiological levels, Rab5 recruitment onto dextran-containing endosomes was detected within ∼30 minutes. In hypertonic conditions, which reduce membrane tension by causing cellular shrinkage, Rab5 was recruited even earlier within ∼20 minutes indicating that a relaxed membrane facilitates early endosomal maturation. In striking contrast, under hypotonic conditions, where cells experience membrane stretching and elevated membrane tension, Rab5 recruitment was completely abolished, and no detectable association of Rab5 with endosomal membranes occurred till ∼40 minutes (Fig. 6H).These results demonstrate that increased membrane tension impairs early endosomal maturation by blocking Rab5 recruitment, while reduced tension accelerates the process, highlighting membrane tension as a key biophysical regulator of phagosomal progression. This suggests that fusion of MEVs with the phagosomal membrane markedly elevates membrane tension, which in turn hampers the recruitment of Rab5 to the membrane (Fig. 3 B-C).

Interestingly, doping the phagosomal membrane with MEV-derived lipids triggered pronounced liquid–liquid phase separation within the bilayer, a process intrinsically linked to changes in membrane line tension (i.e, the free energy per unit length of the interface between coexisting lipid phases). To investigate this effect, we reconstituted giant unilamellar vesicles (GUVs) mimicking the phagosomal lipid composition and incorporated with 25 mol% mycobacterial lipids (Fig. 7A, Supplementary movie 16-17, S15). The vesicles were co-labelled with TopFluor Cholesterol (Lo phase marker, green) and Rhodamine-PE (Ld phase marker, red) to visualise lateral membrane heterogeneity. Through time-lapse Confocal top-view imaging revealed that well-defined *Lo* and *Ld* domains in lipid-doped GUVs. These domains were separated by sharply delineated boundaries, consistent with high *Lo-Ld* interfacial energy. Over time, the domains underwent coarsening, driven primarily by domain coalescence (fusion of adjacent like-phase domains). Both processes are thermodynamically favoured when line tension is high, because the system minimizes total boundary length to reduce the interfacial free energy. From a biophysical perspective, the line tension (γ) at the domain boundary can be described as:

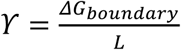

**Figure 7.**
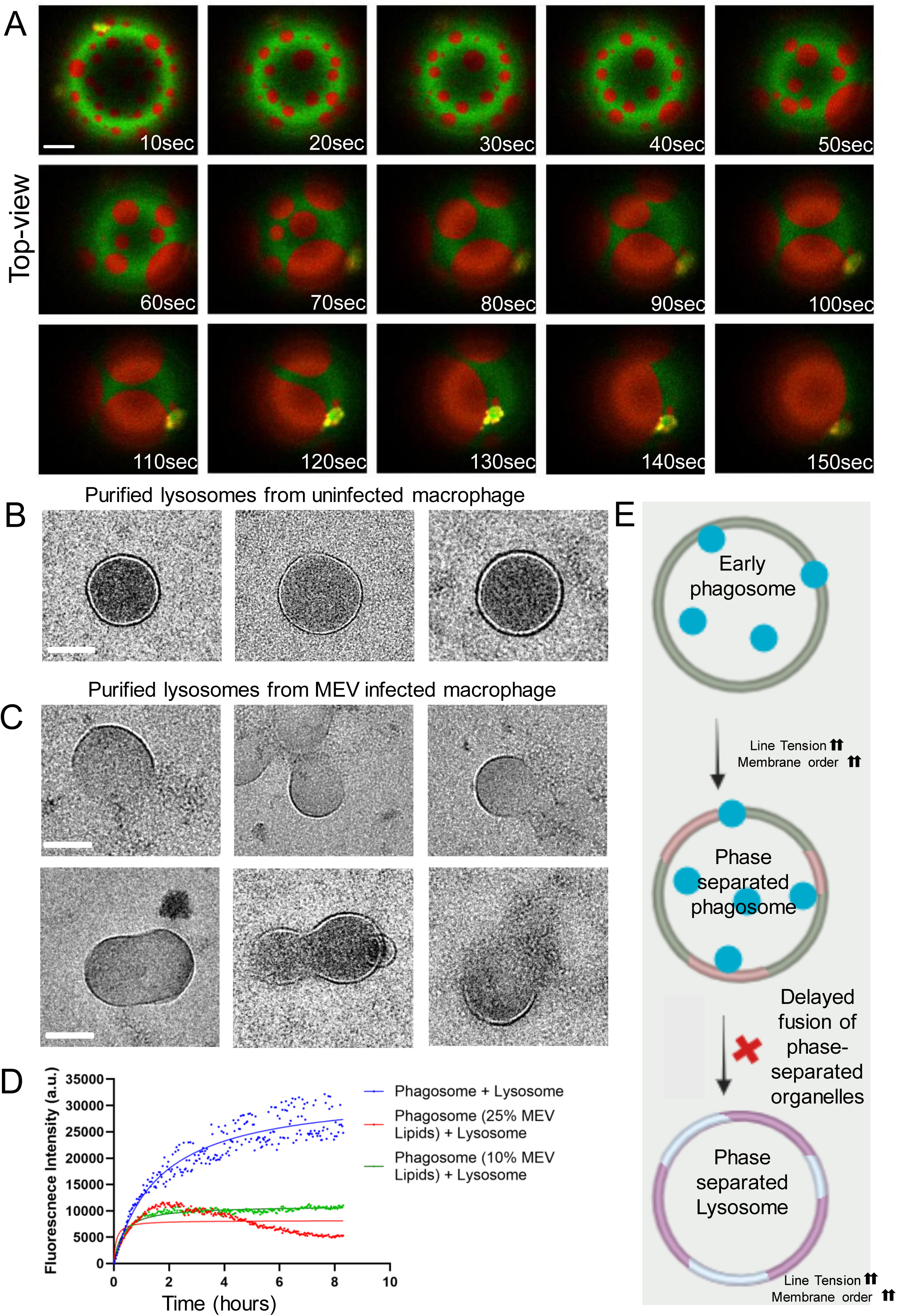
Phase-separated phagosomes and lysosomes exhibit reduced fusion efficiency. **(a)** Time-lapse confocal imaging of phagosomal membrane–mimicking GUVs doped with 25 mol% MEV-derived lipids and labelled with 0.01% Rh-PE (red) and 0.01% TopFluor-Cholesterol (green). Images illustrate the emergence of lipid phase separation upon MEV lipid incorporation. Representative frames from five independent biological replicates are shown. All experiments were performed in PBS (pH 7.4) at 37 °C. Scale bar, 5 μm. Transmission electron micrographs of negatively stained isolated lysosomes from **(b)** uninfected macrophages and **(c)** MEV-treated macrophages. Images illustrate the morphological differences and membrane organization induced upon MEV exposure. Scale bar, 50 nm. **(d)** Förster resonance energy transfer (FRET) assay was performed with lysosomal membrane LUVs labelled with 1% NBD-PE and 1% Rh-PE. Donor fluorescence intensity was monitored as a function of time in the presence of increasing concentrations of MEV lipids. Data points are shown as the means ± S.D. of three independent measurements. All experiments are carried out in PBS pH 7.4 at 37°C. **(e)** Schematic representation illustrating how MEV-driven lipid phase separation increases membrane order and line tension in phagosomal and lysosomal membranes, thereby inhibiting efficient phagosome-lysosome fusion (MEV represented as a blue circle).

Where, ΔG_boundary_ is the excess free energy associated with the phase boundary and L is the boundary length. Increased γ drives the system toward larger, fewer domains, as observed here. In contrast, control GUVs mimicking native phagosomal membranes without MEV-derived lipids displayed uniform fluorescence, indicating the absence of phase separation and negligible line tension under identical imaging conditions. Taken together, these findings suggest that MEV-derived lipids increase *Lo-Ld* interfacial energy, resulting in higher line tension. This physical change promotes domain growth, stabilizes large-scale lateral heterogeneity, and could increase the energetic barrier for membrane mixing, providing a plausible mechanism for MEV-mediated inhibition of phagosome-lysosome fusion. We then wondered whether lysosomes undergo phase separation inside cells? To check this, we purified lysosomes from uninfected as well as MEV infected macrophage. Negative staining transmission electron microscopy revealed a striking difference between the two pools of purified lysosomes (Fig. 7 B-C). Lysosomes purified from uninfected cells exhibited well-defined, continuous circular boundaries with sharp contrast. In contrast, lysosomes from infected cells displayed discontinuous boundaries, with sharp contrast evident only in discrete regions. This discontinuous appearance may be indicative of phase separation within the lysosomal membrane induced by MEV fusion. Phagosomal lysis would typically produce collapsed or irregularly shaped membranes rather than the intact circular profiles observed here. To verify the quality and purity of the isolated lysosomal fraction, we performed western blot analysis using the lysosomal markers LAMP1 and Cathepsin D (CTSD). Both proteins were strongly enriched in the isolated fractions, confirming successful purification of lysosomes (Fig. S14I). Further, we also performed a lysosomal leakage assay using Lucifer Yellow CH, a fluid-phase marker that accumulates within lysosomes. If MEV treatment induced lysosomal membrane permeabilisation or lysis, leakage of Lucifer Yellow from lysosomes into the cytosol would be expected. However, we did not observe any detectable leakage of Lucifer Yellow in MEV-treated cells compared with untreated controls, indicating that lysosomal membranes remain intact and retain their luminal contents following MEV exposure (Fig S14G-H).

Finally, to assess the functional consequence of this mechanically altered state, we examined phagosome-lysosome fusion using a Förster Resonance Energy Transfer (FRET)-based lipid mixing assay (Fig. 7D). Phagosome-mimicking liposomes containing 0, 10, or 25 mol% MEV-derived lipids were mixed with lysosome-mimicking liposomes at a 1:3 molar ratio (lysosome: phagosome) in the presence of 10% (w/v) PEG to simulate cytosolic crowding. In control vesicles, robust lipid mixing was observed, reflected in high FRET efficiency between donor (NBD) and acceptor (Rhodamine) fluorophores. MEV-lipid incorporation caused a dose-dependent reduction in FRET efficiency, accompanied by recovery of donor fluorescence, indicating impaired membrane fusion. The most pronounced inhibition occurred at 25 mol%, correlating with the strongest mechanical alterations. Together, these data suggest that MEV-derived lipids remodel the phagosomal membrane by increasing membrane order and apparent tension thereby reducing its fusogenic potential with lysosomes (Fig. 7E).

### EVs from diverse bacterial species induce conserved membrane remodelling and impair phagosome maturation

To determine whether EV-induced membrane remodelling is conserved across bacterial species, we extended our analysis to extracellular vesicles derived from *Klebsiella pneumoniae* (KEVs; Gram-negative) and *Staphylococcus aureus* (SEVs; Gram-positive). THP-1-derived macrophages treated with SEVs or KEVs were labelled with the mechanosensitive probe Flipper-TR (100 nM), and membrane mechanics were monitored by fluorescence lifetime imaging microscopy (FLIM). Time-resolved measurements over 60 min revealed a rapid and significant increase in Flipper-TR lifetime following EV exposure. KEV treatment increased the lifetime from ∼4.5 ns to ∼5.7 ns, while SEV treatment increased it from ∼4.6 ns to ∼5.4 ns (Fig. 8A-D), indicating a pronounced elevation in host membrane order induced by EVs from both Gram-negative and Gram-positive bacteria.

**Figure 8.**
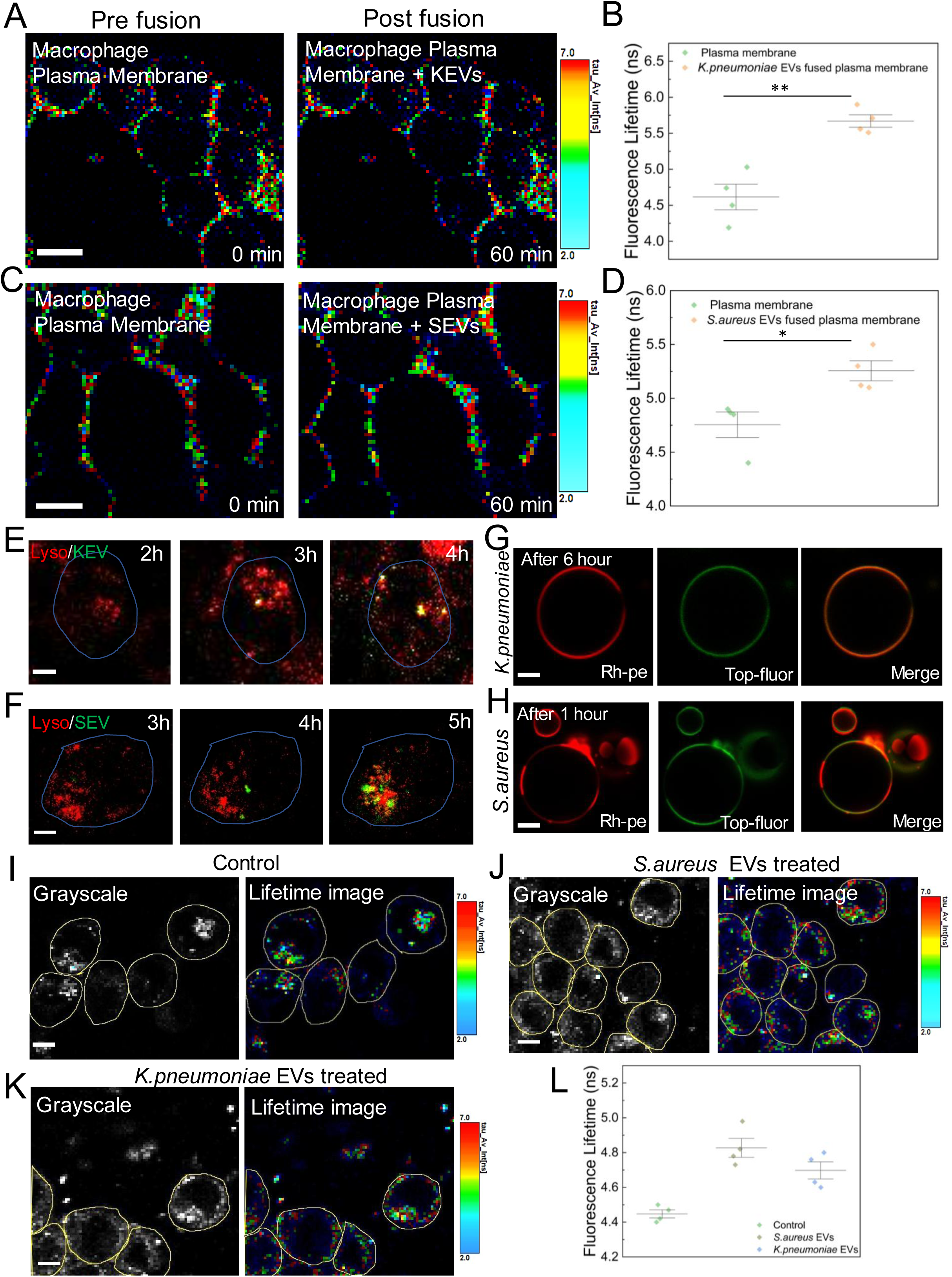
Bacterial extracellular vesicles induced alterations in membrane tension and lipid organisation during phagosome maturation. Fluorescence lifetime imaging microscopy (FLIM) analysis of membrane biophysical changes during KEV and SEV fusion. **(a)** FLIM images of Flipper-Tr labelled live macrophages before and after KEV fusion. **(b)** Quantification of Flipper-Tr fluorescence lifetimes in macrophages as shown in **(a). (c)**FLIM images of Flipper-Tr labelled live macrophages before and after SEV fusion. **(d)** Quantification of Flipper-Tr fluorescence lifetimes in macrophages as shown in **(c).** Data points in **(b)** and **(d)** represent mean ± S.D. from each independent experiment. Live-cell time-lapse confocal imaging of lysosome-phagosome fusion following early endosomal fusion events. (e) *Klebsiella pneumoniae* extracellular vesicles (KEVs), and (f) *Staphylococcus aureus* extracellular vesicles (SEVs) (green) were monitored for subsequent fusion with lysosomes (red). The time-series imaging demonstrates the progressive recruitment and fusion of lysosomes with EV-containing phagosomes. The time interval between frames was 60 minutes. Scale bar, 10 µm. Each condition represents ∼50 cells. Time-lapse confocal imaging of phagosomal membrane–mimicking GUVs incorporating EV-derived lipids.GUVs doped with 25 mol% EV-derived lipids and labeled with 0.01% Rh-PE (red) and 0.01% TopFluor–cholesterol (green) were imaged over time to assess lipid organization. **(g)** KEV-derived lipids and **(h)** SEV-derived lipids. Images demonstrate the emergence of lipid phase separation following EV lipid incorporation. Representative frames from five independent biological replicates are shown. All experiments were performed in PBS (pH 7.4) at 37 °C. Scale bar: 5 µm. FLIM analysis of lysosomal membrane tension in macrophages following treatment with bacterial extracellular vesicles. **(i)** Untreated macrophages, **(j)** Staphylococcus aureus EV– treated macrophages, and **(k)** Klebsiella pneumoniae EV–treated macrophages stained with Lyso-Flipper, a lysosomal membrane tension sensor. All measurements were acquired after vesicle-lysosome fusion, revealing EV-dependent alterations in lysosomal membrane tension. Images are representative of five independent biological replicates (∼150 cells per condition**).(l)** Quantification of Lyso-Flipper fluorescence lifetimes corresponding to the conditions shown in (**i–k**). Each data point represents the mean lifetime from an independent biological replicate. (**p<0.01, ***p<0.001 in one-way ANOVA).

We next examined the kinetics of phagosome-lysosome fusion for EVs from these species. Live-cell imaging showed that KEV-containing phagosomes acquired lysosomal markers at ∼4 h post-internalisation (Fig. 8E), whereas fusion of SEV-containing phagosomes occurred slightly later, at ∼5 h (Fig. 8F). To test whether EV-derived lipids remodel phagosomal membranes in a manner that could influence fusion efficiency, we reconstituted phagosomal membrane mimics doped with 25 mol% KEV- or SEV-derived lipids and monitored lateral membrane organization. KEV-lipid–doped membranes developed clear phase-separated domains after ∼6 h, whereas SEV-derived lipids induced rapid phase separation within ∼1 h (Fig. 8G-H).

Finally, we assessed whether EV fusion perturbs lysosomal membrane mechanics. Using the lysosome-targeted mechanosensitive probe Lyso-Flipper, we observed a significant increase in fluorescence lifetime upon EV fusion, reaching ∼4.7 ns for KEVs and ∼4.85 ns for SEVs (Fig. 8I-L). These data indicate elevated lysosomal membrane order induced by EVs from both species. Collectively, these findings demonstrate that EVs from diverse bacteria induce conserved remodelling of host and lysosomal membranes, generating mechanically stressed compartments that are less permissive to efficient phagosome–lysosome fusion and thereby disrupting lysosomal homeostasis.

## Discussion

Our study identifies Bacterial Extracellular Vesicles (BEVs) as an evolutionarily conserved tools in hijacking phagosome maturation. Using Mycobacteria as a model we first discover that MEV fusion, and notably, MEV-derived lipids alone, is sufficient to remodel the host’s plasma, endosomal and lysosomal membranes by increasing membrane order and stiffness. Such manipulation of host membrane physical parameters, though subtle in magnitude, may have striking functional consequences for intracellular trafficking^76,77^. For example, the modulation of membrane order and tension is known to impact the clathrin-mediated endocytosis as well as intra-luminal vesicle formation and endosome trafficking ^32,39^.

Using a quenching-based assay in vivo, supported by FRET and FLIM-FRET in reconstituted systems, we first established that MEVs spontaneously fuse with the host plasma membrane elevating its membrane order and stiffness (Fig. 1). Mycobacterial virulence lipids such as phthiocerol dimycocerosates, trehalose dimycolate and sulfolipids upon exogenous treatment with cells are known to perturb host signalling by altering the order and stiffness in the plasma membrane ^55,78–81^. However, these minimalist yet elegant studies bypassed the critical question of how these inherently hydrophobic molecules are delivered to their intracellular membrane targets during a natural infection, including the delivery mechanism, dosage, and specific intracellular targeting of these hydrophobic effectors. Our study provides the critical missing link by identifying MEVs as physiological delivery system enabling spatially distributed enrichment of complex envelope lipids and tuning of host membranes. The increased order and apparent tension of the phagosomal membrane delays the critical recruitment of Rab5 (Fig. 3, 6 & 7), an essential GTPase that governs early endosome maturation^5^. Indeed marked increase in the phagosomal membrane tension is expected to alter the lipid packing defects that can impact the Rab binding^82,83^, thus delaying the limiting step for phagosome maturation. Mycobacteria are known to escape out into the host cytosol after rupturing the phagosome by the secretion of virulence factors such as ESAT-6^9,42^ and continue to secrete the MEVs in the host cytosol ^22^. We find that MEV lipids such as derivatives of LAM, PDIM, PIM, and Mycolactone enrich in lysosomes, increasing the lysosomal membrane order and viscosity (Fig. 4 & 5). The enrichment of MEV-derived lipids in the lysosomal membranes results in phase separation and accumulation of line tension at the boundaries owing to the hydrophobic mismatch that might trigger lysis ^84^. Interestingly, the purified lysosomes from the MEV infected cells seem to show a population of both phase separated and lysed lysosomes suggesting lysophagy. The increase in lysosomal viscosity and order induced by MEV lipids could compromise degradative capacity, nutrient recycling, and signalling from lysosomal platforms such as mTORC1 (Fig. 5) ^68^. This mimics the lysosomal dysfunction and metabolic stress reported during live *Mycobacterium tuberculosis* infection ^85^, suggesting that MEV cargo alone can recapitulate core aspects of bacterial virulence thus hijacking not only the pathogen containing host but also bystander cells. The trafficking of MEV lipids to lysosomes prompted us to investigate the fate of early phagosomal compartments. Quantification of membrane tension for early phagosomal compartments is limited by the lack of probes and direct measurements, thus, we resorted to reconstituting phagosomal compartment enriched with MEV-derived lipids *in vitro*. Indeed quantitative measurement by lifetime, time-lapse imaging and micropipette aspiration showed a marked increase in membrane order and tension as a function of MEV-derived lipids enrichment (Fig. 6). Importantly, our FRET-based lipid mixing assay established a direct link between MEV lipids and fusion inhibition. Incorporation of 10-25 mol% MEV lipids into phagosome-mimicking liposomes significantly reduced fusion efficiency with lysosome-mimicking vesicles (Fig. 7). This block correlated with the increased membrane order detected by FLIM-Flipper-TR measurements, consistent with the principle that elevated order and tension increase the energetic barrier for hemifusion and pore opening^38,86,87^. Thus, MEV lipids act as mechanical inhibitors of fusogenicity, rewiring the phagosomal membranes to resist lysosomal fusion.

A striking transcriptional reprogramming of the host reinforces the organelle-wide mechanical rewiring. RNA-seq of MEV-infected macrophages revealed suppression of key phagosomal maturation genes (*Rab7, Syntaxin, VAMP8*) and upregulation of lysosomal stress (*CTSD, CTSL, LGALS3*) and membrane repair (*CHMP4B*) markers. This indicates there is a tug-of-war situation between the MEV induced manipulation and the host response to counter it. Plasma membrane tension is known to activate mTORC2 and PLD pathway ^30,88^, that in turn regulates the expression of TFEB, a master regulator of lysosomal biogenesis. RNA seq data for MEV infected macrophage indicates an upregulation of both mTORC2 and PLD genes and downregulation of TFEB causing the delay. Thus, we propose that MEV mediated rise in the host membrane order and likely apparent tension reprograms host transcription machinery of phagosome maturation via mTORC2/PLD - TFEB axis.

Surprisingly, extracellular vesicles of *S. aureus* and *K. pneumonia* were also observed to delay phagosome maturation by altering the order of cellular membranes (Fig. 8). This establishes bacterial EVs an evolutionary conserved mechanism that enable pathogens to hijack host and bystander cells through mechanical, rather than solely biochemical, manipulation remotely. Despite structurally distinct, the pathogens can exploit a converging biophysical mechanism by delivering more ordered EV lipid cargo that alters membrane order across phagosome and lysosome. Subsequently, the fusion of EVs creates an energy barrier that blocks phago-lysosomal fusion and disrupts homeostasis (Fig. 9). This dual mode of inhibition provides a powerful survival advantage to the bacterium by preventing phagosomal maturation at the site of entry, but also renders lysosomes intrinsically less competent to fuse with incoming phagosomes (Fig. 4A-B). This reveals a broader paradigm in pathogenesis - the exploitation of host membrane mechanics via extracellular vesicles provides a potent, lipid-driven generic strategy to subvert cellular processes, complementing the protein-based effectors.

**Figure 9.**
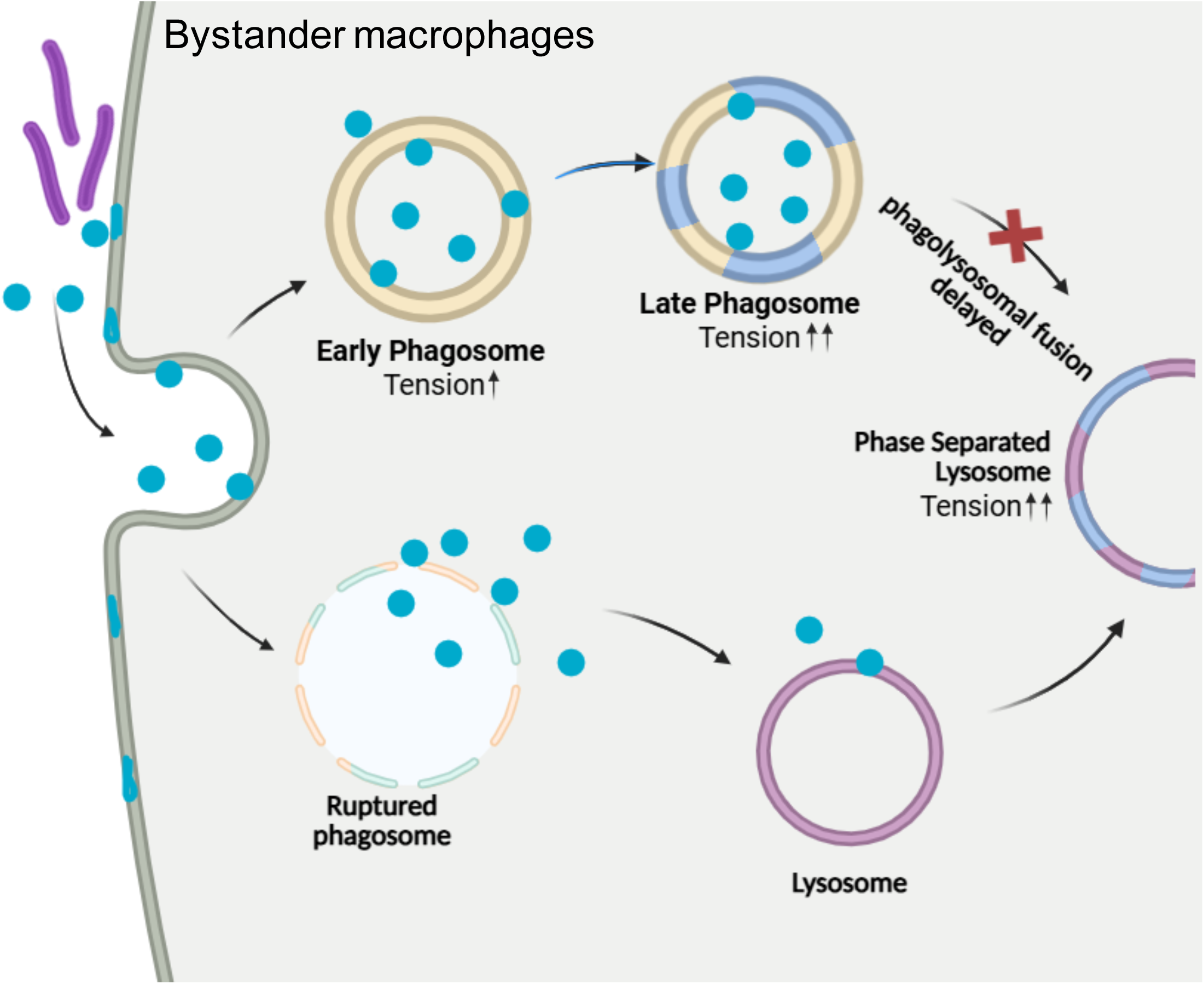
Schematic representation of the proposed mechanism by which MEVs mediate abortion or delay of the phagosomal maturation process in bystander macrophages. Fusion of mycobacterial extracellular vesicles (MEVs) (shown as blue circle with the host membrane remodels its lipid composition, driving membrane ordering and phase separation (depicted as a two-colour membrane). Such mechanical remodelling of membranes effects phagosome maturation, ultimately aborting phagosome-lysosome fusion and promoting bacterial survival within host cells.

### Material and method

### Materials

1,2-dioleoyl-*sn*-glycero-3-phosphocholine (DOPC), 1,2-dioleoyl-*sn*-glycero-3-phosphoethanolamine (DOPE), BSM, di-stearoyl phosphatidyl ethanolamine-PEG (2000)-biotin and cholesterol were purchased from Avanti Polar Lipids. Fluorescent conjugated lipid lissamine rhodamine B sulfonyl (Liss Rhod PE) was also purchased from Avanti Polar Lipids. Avidin coated on the chamber slides was purchased from Sigma-Aldrich. PKH-67 dye used to label MEVs procured from Sigma-Aldrich. Roswell Park Memorial Institute medium (RPMI-1640), 1× Dulbecco’s phosphate buffer saline (D-PBS), sodium bicarbonate, 20× antibiotic–antimycotic solution, amphotericin-B, streptomycin, penicillin, paraformaldehyde, PMA (phorbol 12-myristate-13-acetate), were purchased from Sigma-Aldrich.

### Mammalian cell culture

THP-1 monocytes were cultured in Roswell Park Memorial Institute (RPMI) 1640 medium (Gibco) supplemented with 10% fetal bovine serum (FBS) and maintained to confluence in a 5% CO₂ incubator. Cells were seeded onto appropriate culture dishes and differentiated into macrophages using 50 ng/mL phorbol 12-myristate 13-acetate (PMA; Sigma-Aldrich). After 24 h of PMA treatment, the medium was replaced, and cells were allowed a 24 h resting phase before experiments. These cell lines were not re-authenticated in our laboratory. For overexpression experiments, differentiated THP-1 macrophages were transfected with the respective constructs using Lipofectamine LTX (Invitrogen) following the manufacturer’s protocol. For LysoTracker assays, cells were incubated with 70 nM LysoTracker Red (Invitrogen) for 30 min before imaging.

### Mycobacteria and bacterial culture

Mycobacterium tuberculosis (*Mtb*) H37Ra (avirulent), M. smegmatis were cultured in minimal medium (MM) containing 1 g L⁻¹ KH₂PO₄, 2.5 g L⁻¹ Na₂HPO₄, 0.5 g L⁻¹ asparagine, 50 mg L⁻¹ ferric ammonium citrate, 0.5 g L⁻¹ MgSO₄·7H₂O, 0.5 mg L⁻¹ CaCl₂, and 0.1 mg L⁻¹ ZnSO₄, supplemented with 0.1% (v/v) glycerol (pH 7.0) and with or without 0.05% (v/v) tyloxapol. In certain experiments, cultures were maintained in Middlebrook 7H9 (M7H9) broth supplemented with 10% (v/v) oleic acid-albumin-dextrose–catalase (OADC; BD Microbiology Systems) and 0.5% (v/v) glycerol. Cultures were incubated at 37 °C for up to 10 days before vesicle isolation. Cell viability at day 10 was quantified by enumerating colony-forming units (CFUs) obtained from serial dilutions plated on Middlebrook 7H11 agar.

### Isolation, purification and fluorescent labelling of Mycobacterial Extracellular Vesicles (MEV)

Extracellular vesicles were isolated as previously described ^22^ with minor modifications to enhance yield and purity. Briefly, 1 L bacterial cultures were harvested by centrifugation at 3,450 × g for 15 min at 4 °C, and the cell-free supernatants were passed through 0.45 µm polyvinylidene difluoride (PVDF) filters (Millipore) to remove residual cells. Filtrates were concentrated ∼20-fold using an Amicon Ultrafiltration System (Millipore) equipped with a 100 kDa molecular weight cut-off membrane. The concentrate was sequentially centrifuged at 4,000 × g and 15,000 × g for 15 min each at 4 °C to remove remaining cell debris and aggregates. Vesicles were then pelleted by ultracentrifugation at 100,000 × g for 1 h at 4 °C. The crude vesicle pellet was resuspended in 1 mL of 10 mM HEPES (pH 7.4) containing 150 mM NaCl, and mixed with 2 mL of 35% (w/v) OptiPrep solution (Sigma-Aldrich) prepared in the same buffer. This suspension was overlaid with a discontinuous OptiPrep density gradient (30-35% w/v) and centrifuged at 100,000 × g for 16 h at 4 °C. Following centrifugation, 1 mL fractions were collected from the top of the gradient, dialyzed individually against PBS overnight at 4 °C, and recovered by ultracentrifugation (100,000 × g, 1 h, 4 °C). Purified vesicles were finally resuspended in PBS and stored at 4 °C for immediate use. Vesicle size distribution and polydispersity were determined by dynamic light scattering (DLS), and protein quantification was done using BCA estimation. For intracellular trafficking studies, MEVs were fluorescently labelled with PKH67 or PKH26 membrane dyes according to the manufacturer’s instructions. Unincorporated dye was removed using size-exclusion chromatography to eliminate free dye and dye aggregates before downstream cellular uptake and imaging experiments.

### Nanoparticle Tracking Analysis (NTA)

The size distribution and concentration of isolated membrane vesicles were determined using Nanoparticle Tracking Analysis (NTA, Malvern Panalytical) at 405nm. Vesicle samples were diluted 100-fold in sterile phosphate-buffered saline (PBS) to achieve optimal particle concentration for measurement.PBS alone was used as a negative control to ensure the absence of particulate contaminants and validate instrument performance. NTA provided quantitative data on vesicle size distribution and particle number in the stock solution, confirming the homogeneity and quality of the extracted vesicles.

### Light Microscopy

Eight-well chambered slides (Ibidi) were coated with 10 µL of 1 mg/mL streptavidin, followed by the addition of 200 µL of biotinylated GUVs. The slide was incubated for 15 min to allow immobilization of the GUVs. MEVs were then added into the chamber. Imaging was performed on an Olympus FV3000 confocal microscope using the appropriate excitation lasers. Identical laser power and detector gain settings were maintained for all experimental conditions.

### Fluorescence Recovery After Photobleaching (FRAP)

For FRAP measurements, GUVs were doped with 0.1% Atto-PE and incubated with MEVs for 1 h to allow efficient fusion. Pre-bleach images were acquired at reduced laser intensity. Photobleaching of the selected ROI was performed at full laser power for 30 s at the equatorial plane of the GUV. After bleaching, the laser intensity was returned to the attenuated setting and fluorescence recovery was recorded over time. FRAP measurements for each condition were repeated three times, and recovery curves were normalized before analysis.

### Preparation of GUV

Giant unilamellar vesicles (GUVs) were formulated to mimic the lipid composition of the phagosomal membrane, comprising 40 mol% 1,2-dioleoyl-sn-glycero-3-phosphocholine (DOPC), 5.5 mol% 1,2-dioleoyl-sn-glycero-3-phosphoethanolamine (DOPE), 22 mol% brain sphingomyelin (BSM), 32.5 mol% cholesterol, 0.03 mol% 1,2-distearoyl-sn-glycero-3-phosphoethanolamine-polyethylene glycol-biotin (PEG-biotin), and 0.1 mol% Lissamine Rhodamine PE (Rhod-PE). GUVs were generated via electroformation as described previously^42^, with modifications to optimize yield. Briefly, 15 µL of the lipid mixture (5 mg mL⁻¹ in chloroform) was uniformly deposited onto indium-tin-oxide (ITO) - coated glass slides (Nanion Technologies GmbH) and dried under vacuum for ≥2 h to remove residual solvent. The resulting lipid film was rehydrated in phosphate-buffered saline (PBS, pH 7.4; 300 ± 5 mOsm) and subjected to electroformation under a sinusoidal AC field (2 V, 10 Hz) at 60 °C for 3 h to promote GUV growth.

### R18 dequenching (lipid mixing) fluorescence assay

For R18 labelling of MEVs, MEVs were fluorescently labelled using Octadecyl Rhodamine B chloride (R18 dye). A total of 0.6 μL of R18 (optimised concentration) was added to the MV suspension and sonicated in a water bath sonicator for 30 min, with alternating cycles of 3 min sonication followed by 5 min rest to achieve uniform dye incorporation under a highly quenched state. Unbound dye was removed using G-50 MicroSpin columns (Cytiva, Cat. No. 27533001). The columns were first centrifuged for 3 min to remove the storage buffer and compact the resin. 100 μL of R18-labelled vesicles were then carefully loaded onto the column and centrifuged at 450 × g for 3 min. The eluate containing purified R18-labelled MVs was collected for subsequent use.

The R18 dequenching assay was performed to monitor the membrane fusion kinetics of MEVs as previously described ^28^. Fusion-induced dilution of the self-quenching dye R18 increases in fluorescence intensity. R18-labelled MEVs were mixed with unlabeled plasma membrane-mimicking liposomes (DOPC/DOPE/SM/Chol) at a 1:9 molar ratio in PBS (pH 7.4). Fluorescence emission at 595 nm was recorded over time using a spectrofluorometer (excitation: 545 nm; slit width: 5 nm for both excitation and emission).

The percentage of membrane fusion was calculated using the following equation:

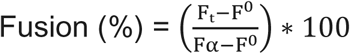

where F₀ is the fluorescence at time zero, Fₜ is the intensity at time t, and Fα represents the maximum fluorescence obtained after adding Triton X-100 to fully disrupt the membranes and achieve infinite dye dilution.

For cellular fusion assays, the fusion efficiency of MVs with host cells was assessed using differentiated THP-1 macrophages. Purified R18-labelled MVs were added at an optimized ratio of 200 vesicles per cell. Membrane fusion was quantified by monitoring R18 fluorescence dequenching, which reflects dilution of the self-quenching dye upon fusion of the labelled vesicles with the host cell membrane.

### Forster Resonance Energy Transfer-based lipid mixing assay

Fusion between plasma membrane-mimicking LUVs and MEVs was quantified by monitoring changes in FRET efficiency between NBD-PE (donor) and Rh-PE (acceptor). Lipid-mixing kinetics were assessed by measuring donor fluorescence dequenching over time, reflecting dilution of the FRET pair upon fusion. Probe-containing LUVs were mixed with probe-free vesicles at a 1:9 ratio, maintaining a total lipid concentration of 200 µM. Donor emission was recorded at 530 nm (excitation: 465 nm) for 4 h using a CLARIOstar microplate reader.

For phagosome-lysosome FRET assays, fluorophores were incorporated into lysosomal membranes, and donor fluorescence changes were monitored following interaction with phagosomes containing varying molar percentages of MEV-derived lipids. All experiments were performed in PBS (pH 7.4) at 37 °C.

### Lipid Monolayer experiments

Monolayer measurements were conducted using a KSV NIMA Langmuir balance equipped with two movable barriers and a Wilhelmy plate microbalance (filter paper plate) for surface pressure detection. Lipid monolayers composed of DOPC:DOPE:SM:Chol were prepared by spreading a lipid–chloroform solution (1 mg mL⁻¹) dropwise onto the autoclaved and filtered double-deionized water maintained at 25 °C. After spreading, the monolayers were allowed to equilibrate for 15 min to enable complete solvent evaporation. For membrane conditions incorporating MEV lipids, the lipids were added to the monolayer mixture at increasing molar percentages. Surface pressure-area (π-A) isotherms were recorded during symmetric compression of the monolayer at a constant barrier speed of 1 mm min⁻¹ until the collapse pressure (πc) was reached. The Langmuir trough was thoroughly cleaned before each experiment by sequential rinsing with ethanol and filtered double-deionized water until the surface pressure returned to zero. All experiments were performed in triplicate.

### Micropipette Aspiration

Micropipette aspiration experiments were performed on an Olympus IX83 epifluorescence microscope (100× objective) equipped with Hoffman modulation contrast optics. Three-axis hydraulic micromanipulators were used to precisely position the micropipette. Borosilicate glass capillaries (10 μm inner diameter) were pulled using a Narashige Micro Forge MF-900 and Sutter Instruments P-87 puller to generate micropipettes. Approximately 100 μL of GUV suspension was deposited on a coverslip for analysis.

A controlled suction pressure (ΔP) was applied to aspirate vesicles into the pipette, generating a uniform membrane tension (τ) given by:

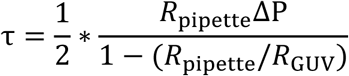

where *_R_*_pipette_ and *_R_*_GUV_ represent the radius of the pipette and GUV, respectively.

The projection length (Lp) inside the pipette was monitored to quantify membrane area changes. The apparent area increase (Δa) was calculated using:

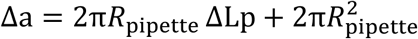

The reference area (a_0_) for a spherical vesicle was defined as:

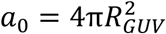

The areal strain was thus determined by:

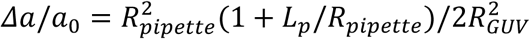

Changes in Lp with varying ΔP were recorded (Fig. 4d). Membrane area expansion was analyzed over a tension range of 76–550 μN/m (Fig. 4e). The area stretch modulus (K_a_) was obtained from the slope of the τ-Δa plot, while the bending modulus (K_b_) was determined from the slope of the semi-log(τ)-Δa plot. All measurements were performed at room temperature.

### Transmission Electron Microscopy

Negative staining was performed as previously described^42^, with minor modifications. Briefly, purified MEVs (10 μL) were adsorbed onto carbon-coated copper grids for 40 min and excess liquid was removed using qualitative filter paper (Himedia). Grids were fixed with 10 μL of 2.5% (v/v) glutaraldehyde, rinsed with deionized water, and subsequently stained with 10 μL of 2% (w/v) uranyl acetate for 3 min. After an additional rinse with deionized water, grids were vacuum-dried overnight. TEM imaging was performed using a JEOL JEM-2100F transmission electron microscope. For vesicle-vesicle fusion assays, MEVs and large unilamellar vesicles (LUVs) were co-incubated on carbon-coated grids before fixation and staining, following the same procedure described above.

### TLR-2 Surface staining and Flow cytometry

Surface staining was performed as previously described^89^, with minor modifications. Briefly, THP-1 cells were seeded in 6-well plates and differentiated into macrophages by treatment with phorbol 12-myristate 13-acetate (PMA) for 12-14 h. Differentiated cells were washed and blocked with 1% BSA for 30 min at 4 °C to prevent nonspecific antibody binding. Cells were then incubated with primary antibodies against TLR2 for 30 min at 4 °C, followed by staining with Alexa Fluor 488-conjugated secondary antibodies for an additional 30 min at 4 °C in the dark. After staining, cells were washed with FACS buffer (1× PBS supplemented with 1% BSA and 0.01% NaN₃) and immediately analyzed by flow cytometry.

### Fluorescence Lifetime Imaging Microscopy (FLIM)

Fluorescence lifetime imaging microscopy was performed on an MT200 time-resolved confocal microscope (PicoQuant, Berlin, Germany) equipped with a time-correlated single-photon counting (TCSPC) unit. Excitation was provided by a 488 nm and 519 nm pulsed laser was used as per the experimental requirement, with power adjusted between 10-100 nW (measured post-dichroic). Samples were mounted in glass-bottom wells positioned on a 60× water-immersion objective. A 488 nm and 519 nm dichroic mirror served as the main beam splitter as per the flurophore. Out-of-focus fluorescence was rejected using a 50 μm pinhole, and in-focus emission was split by a 50/50 beam splitter into two detection paths. Fluorescence photons were detected by single-photon avalanche diodes (SPADs). Data acquisition and lifetime analysis were performed using SymPhoTime 64 software (PicoQuant GmbH, Berlin, Germany). For plasma Membrane tension quantification assays, 100nM of Flipper-tr were incubated for 30 minutes before imaging. For lysosomal membrane tension quantification, 1µM of Lyso-flipper was incubated for 15 minutes before imaging. For Laurdan experiments, 5µM of Laurdan were incubated for 15 minutes before imaging. All FLIM were acquired under optimised conditions and the fitting was performed using the tail-fitting approach, considering the overall frame as previously reported^29^.The fitting was done with a model parameter of 2 and a reduced chi-square value close to 1, with photon counts in the range between 10^2^ to 10^3^ photons per pixel. Furthermore, the apparent pixelation is an intrinsic feature of rapid FLIM acquisition and anisotropy reconstruction and does not affect the quantitative anisotropy measurements used for statistical analysis, consistent with previous literature^29,33,90^.

### Fluorescence Lifetime Imaging Microscopy (FLIM) - FRET

For FLIM–FRET measurements, NBD was used as the donor and R18 as the acceptor. Donor excitation was carried out at 488 nm, and emission was collected above 519 nm, with R18 emission spectrally excluded to ensure single-channel donor lifetime detection. Donor-only (NBD-labeled) and donor–acceptor (NBD+R18) samples were imaged under identical acquisition settings. All measurements were conducted at room temperature, with at least three independent biological replicates.

### Immunofluorescence Staining

THP-1–derived macrophages (5 × 10⁴ cells per coverslip) were seeded onto sterile glass coverslips. Cells were gently rinsed with 1× PBS and fixed with 4% paraformaldehyde (PFA; Thermo Fisher Scientific) for 20 min at room temperature. Excess fixative was removed by three washes with 1× PBS (5 min each). Cells were permeabilised with 0.1% Triton X-100 for 10 min and blocked with 1% BSA for 1 h at room temperature. cells were incubated overnight in a humidified chamber with anti-LAM primary antibody (Thermo Fisher; 1:50), LC3B (Abclonal), EEA1(Abcam), LAMP1 (Abclonal). After three washes with 1× PBS, Alexa Fluor 488-conjugated anti-rabbit secondary antibody (Invitrogen) was applied for 45 min in the dark. Samples were washed three additional times with 1× PBS, and coverslips were mounted for imaging.

### Lysosomal Isolation

THP-1 cells were cultured to ∼90% confluency in 100 mm dishes and differentiated into macrophages using phorbol 12-myristate 13-acetate (PMA). Lysosomes were isolated using a commercial isolation kit (Sigma-Aldrich, cat. no. LYSISO1) following the manufacturer’s instructions with few modifications. Briefly, cells were harvested by centrifugation at 600 × g for 5 min, resuspended in 2.7 volumes of 1× extraction buffer, and homogenized using a glass Dounce homogenizer. Cell disruption was monitored by trypan blue staining until ∼80–85% of cells were broken. Nuclei were removed by centrifugation at 1,000 × g for 10 min, and the post-nuclear supernatant was centrifuged at 20,000 × g for 20 min to obtain a crude lysosomal pellet. This pellet was resuspended in 0.4 mL of 1× extraction buffer per 1 × 10⁸ cells. For enrichment, crude lysosomes were further purified by density gradient centrifugation on a multistep OptiPrep gradient (Sigma-Aldrich) at 150,000 × g for 4 h. Fractions (0.5 mL) were collected sequentially from the top of the gradient for downstream analysis.

### Lipid Extraction from Mycobacterial Extracellular Vesicles (MEVs)

Lipids were extracted from MEVs as previously described ^91^, with minor modifications. Briefly, purified MEVs were washed three times with cold sterile PBS and resuspended in 1 mL PBS in a glass vial. Lipids were extracted by adding 3 mL of chloroform-methanol (2:1, v/v), followed by vortexing and centrifugation at 4,000 rpm for 10 min at 4 °C. The lower organic phase was collected, while the remaining aqueous phase was acidified with 2% formic acid and re-extracted with 2 mL chloroform to enrich phospholipids. After centrifugation under the same conditions, the organic phase was collected and pooled with the initial extract. Combined organic phases were dried under a gentle stream of nitrogen gas at room temperature and stored at -80 °C until further use. Total lipid content was quantified as previously described ^92^.

### MEVs Uptake Assay

THP-1-derived macrophages were pretreated with methyl-β-cyclodextrin (MβCD) (5mM) and cytochalasin D (10μM) for 1 h at 37°C . Following pretreatment, the cells were incubated with PKH-26 labelled MEVs for 2 h. Cells were then washed with PBS and fixed with 4% paraformaldehyde (PFA) for 15 min at room temperature. After fixation, nuclei were stained with Hoechst (1:4000 dilution) for 5 min at room temperature. Images were acquired using a confocal microscope under identical imaging settings for all experimental conditions. MEV uptake was quantified by measuring the number of fluorescent puncta per cell using ImageJ software.

#### MDLV preparation

Estimated MEV-derived lipids were used to prepare thin lipid films in amber vials (Avanti-Sigma) for the generation of MEV-derived lipid vesicles (MDLVs). Briefly, chloroform was evaporated under a gentle nitrogen stream, followed by vacuum drying for 2 h to remove residual solvent. The resulting dry lipid films were rehydrated in 1 mL PBS (pH 7.4). MDLV preparation involved heating the suspension in a hot water bath at 80 °C and vortexing to ensure complete resuspension. The vesicles were then extruded using an Avanti® Mini Extruder through a 100 nm polycarbonate membrane with 10 mm support filters. A minimum of 25 extrusion passes was performed. Following extrusion, vesicle size distribution was confirmed using dynamic light scattering (DLS). Furthermore, the concentration was estimated using the method as mentioned earlier.

#### Atomic Force Spectroscopy

MFP-3D atomic force microscope (Asylum Research, Santa Barbara, CA, USA) with silicon nitride cantilevers (Oxford Instruments) and a nominal resonance frequency and a spring constant of 30 kHz and 0.16 N/m, respectively, were used for force spectroscopy experiments in contact mode. Cantilever calibration was done using the thermal noise method to obtain a spring constant of 0.10-0.19 N/m. The applied load was 1 nN, and the cantilever velocity was fixed at 6 μm/s. Differentiated THP-1 cells were incubated with MVs at ratio of 1:200 (cell: vesicles) for different time points i.e., 2, 6 and 24 h. The force curves were recorded in the region of the cell body, which was between the cell periphery and the nucleus. In each set of independent experiments, at least 100 cells were used for force spectroscopy. Igor software (Asylum Research) was used to calculate the elastic modulus by fitting the force curve using the Hertz model after providing the necessary data about the tip geometry and Poisson’s ratio of the sample. For analysis of membrane tethers (tether force, tether number, and lengths), a reported custom-made MATLAB program was used. Actin tether force is measured using AFM force spectroscopy during the retraction phase of a force-distance (force-extension) experiment. The AFM tip first contacts the cell surface and is allowed to adhere to the plasma membrane. As the cantilever retracts, if the adhesion between the tip and the membrane is stronger than the membrane’s attachment to the underlying actin cortex, a thin membrane tether is pulled instead of the tip detaching immediately. The force required to maintain this tether is referred to as the tether force, which reflects membrane tension, membrane bending rigidity, and membrane-actin cytoskeletal adhesion. The tether force is obtained from the force-distance curve recorded during cantilever retraction. Tether formation appears as a nearly horizontal force plateau (or a series of plateaus and rupture steps) on the retraction curve. AFM analysis software identifies the plateau region, often after smoothing the data, calculates the mean force over this region, and reports this value as the tether force.

### F-actin Immunofluorescence and Actin Isosurface Imaging

Differentiated THP-1 cells were incubated with MVs at ratio of 1:200 (cell: vesicles) for different time points, i.e., 2, 6 and 24 h. Later on, cells were fixed with 4% paraformaldehyde for 10 min at RT and rinsed with 1× PBS for three times. Then, cells were permeabilized using 0.1% Triton-X-100 in 1× PBS, followed by blocking in 1% BSA for 1.5 h and rinsed with PBS/PBST. TRITC-Phalloidin was used to stain F-actin (2 μg/mL) for 30 min in the dark; it was rinsed thoroughly with 1× PBS, and slides were mounted with mounting media. Images were taken on a laser scanning confocal microscope (Olympus FV3000) using a 60×/1.4 NA objective. The 3D reconstruction of the filaments from confocal stacks was carried out with Cellsense (Evident Scientific) software. 60 to 80 cells in each set (n=2; N=3) for various conditions were analyzed. The approximate volumes enclosed by isosurfaces were normalized by the projected area of the cells, which was obtained using 3D Object Counter Fiji plugin. The significance level was assessed by One-way ANOVA using GraphPad Prism software version 10.1.0. Post-acquisition image analysis for F-actin structures was performed using the FiloQuant plugin in Fiji software. Relative quantification was determined by calculating the number of punctae (ITCN plugin, ImageJ), filaments (FiloQuant plugin, Fiji) of F-actin per cell. Actin anisotropy was done using the Fibril Tool Fiji pluginA minimum of 120-150 cells from randomly selected fields was scored per condition.

### Transcriptome profiling

#### RNA Extraction and Quality Assessment

Differentiated THP-1 macrophages treated with MEVs for 2 h, 12 h, or 24 h were harvested for RNA extraction. Total RNA was isolated using TRIzol Reagent (Invitrogen) according to the manufacturer’s protocol. RNA concentration and purity were assessed using a NanoDrop spectrophotometer (Thermo Fisher Scientific).

#### Library Preparation and RNA Sequencing

RNA sequencing and library preparation were performed by Eurofins Genomics Pvt. Ltd. Libraries were generated using the TruSeq Stranded Total RNA Library Prep Kit (Illumina) with rRNA depletion. cDNA was synthesised using random primers. Paired-end sequencing libraries (2 × 150 bp) were prepared following the manufacturer’s instructions and sequenced on an Illumina platform.

#### RNA-seq Data Processing and Analysis

Raw reads from the samples were processed using Trimmomatic v0.39 to remove adapter sequences, ambiguous bases (>5% N), and low-quality reads (>10% bases with Phred QV < 25). Reads shorter than 100 nt after trimming were discarded. High-quality paired-end reads (QV > 25) were retained for alignment.

Reads were mapped to the Homo sapiens GRCh38 reference genome (Ensembl release 111) using STAR v2.7.10a with default parameters. Gene-level quantification was performed with featureCounts. Differential gene expression analysis was carried out using DESeq2, and functional enrichment analysis was performed using ClusterProfiler. All downstream analyses and visualisations were conducted in R. However, the experiments in Fig. 3G and 5Q are the same experiment representing different gene subsets.

### Lipidomics analysis

#### Liquid Chromatography

Extracted lipids were then analysed in both positive and negative ion mode using a 1290 Infinity UHPLC System attached to a 6550 iFunnel Q-TOF system (Agilent Technologies, USA). The lipids were identified using databases available online. An Agilent 1200 HPLC (Agilent Technologies; Palo Alto, CA) with a 2.1 inner diameter (ID) × 150 mm, 3.5 μm XBridge C18 column (Waters Corp.; Milford, MA) heated to 45°C was used with a binary solvent system and a flow rate of 0.175ml/min (Sartain et al., 2011). Equilibration was performed with 95% solvent A (0.1% formic acid in filtered MilliQ water) and 5% solvent B (0.1% formic acid in acetonitrile). A 5 μL aliquot of lipid extract (equivalent to 20 μg of dried extract) was injected onto the column. The gradient began with the initial solvent composition held for 5 minutes, followed by a 24-minute linear increase to 100% solvent B, which was then maintained for an additional 5 minutes.

#### Mass Spectrometry

An Agilent Mass Spectrometer Quadrupole time-of-flight (MS Q-TOF) G6545XT equipped with a Dual AJS ESI mode source was used for accurate mass analysis of the LC eluent. Positive- (+) and negative- (-) ion data were generated by operation of the mass spectrometer with a capillary voltage of 4000 V, nebulizer of 45 psig, drying gas of 8 l/min, gas temperature of 300°C, fragmentor of 175 V, charging voltage of 2000 V, skimmer of 65 V, and octopole radio frequency voltage of 750 V. Mass spectra were acquired at a rate of 1.00 spectra/s and data were collected as profiled spectra over a mass range of 100 to 3,200 Da. Mass calibration was performed with an Agilent tune mix from 100 to 3,200 Da. Data were collected with the Agilent Qualitative Analysis of MassHunter Acquisition Data software, version 10.0. Positive-ion mass spectra were acquired in Auto MS/MS mode, and collision energies with slope of 6.5 V/100 Da and offset 2.0V were used for fragmentation.

#### Quantification of lipids

LC/MS data files were analyzed using MassHunter Qualitative Analysis Software, version B.10.00 (Agilent Technologies, Santa Clara, CA). Molecular features (MFs) were extracted from the raw data using the Molecular Feature Extraction (MFE) algorithm, which identifies covariant ions, such as isotopes and charge states, from accurate mass LC/MS data and consolidates them into a single feature. The parameters for MFE were set to extract small molecules with a minimum peak filter of 1000 counts. Ion species were limited to +H, +Na, +NH4, +K with a peak spacing tolerance of 0.0005 m/z plus 5.0 ppm. The isotope model was configured for common organic molecules, and the charge state range was restricted to 1-2. No compound filters, mass filters, or mass defect constraints were applied. The extracted MFs were then identified using MassHunter through compound screening based on formulas obtained from the Mtb LipidDB and MycoMass databases for Mtb MV and MV fused with lysosome samples and Lipid Maps https://www.lipidmaps.org/ (mammalian lipid database) used for MV fused lysosome and only lysosome samples. The search was conducted with parameters set to match mass values only, using a match tolerance of 5 mDa with a charge state range of 1-2. From the resulting compound list, the quantitative abundance of each identified molecule was calculated using the following formula.

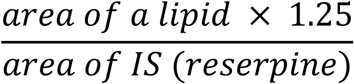

where, 1.25 is the ppm (concentration) of reserpine (internal standard) used.

Molarity of a lipid =

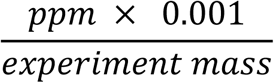

Mol% of each lipid =

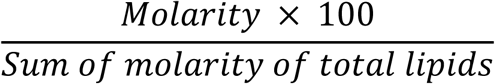

#### Fluorescence Anisotropy

The fluorescence anisotropy measures the rotational flexibility around the fluorophore in a microheterogeneous medium. Fluorescence anisotropy measurements of DPH (1,6-diphenyl-hexa-1,3,5-triene) and its derivative, TMA-DPH (trimethyl ammonium diphenyl hexatriene), were performed using the 96-well Clariostar microplate reader containing host membrane and MEVs. The excitation and emission wavelength were kept at 355 nm and 430 nm, respectively. The milianisotropy of DPH and TMA-DPH in 200 μΜ lipid was monitored by varying the temperature from 25 °C to 42 °C by using the following equation:

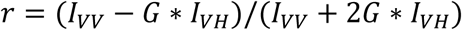

Where G=I_HV_/I_HH,_ (G-factor), I_VV_ and I_VH_ are the measured fluorescence intensities with the excitation polarizer vertically oriented and the emission polarizer vertically and horizontally oriented, respectively.

#### Lysosome leakage assay

THP-1-derived macrophages were incubated with Lucifer Yellow CH in serum-free RPMI medium for 1 h at 37°C to label lysosomes. Following incubation, the cells were washed three times with PBS to remove excess dye, then incubated in fresh complete RPMI medium for an additional 1 h at 37°C to allow dye accumulation within lysosomal compartments. The cells were then treated with MEVs for 12 h. After treatment, the cells were fixed with 4% paraformaldehyde (PFA) for 15 min at room temperature, washed with PBS, and imaged using identical acquisition settings.

#### Immunoblotting

Following lysosome isolation from differentiated THP-1 cells, both the isolated lysosomal fraction and the corresponding whole-cell lysate were lysed in 125 µL of RIPA buffer. Protein concentrations were determined using a BCA Protein Assay Kit (Thermo Scientific) according to the manufacturer’s instructions. Equal amounts of protein (15 µg per sample) were resolved on a 10% SDS-polyacrylamide gel and subsequently transferred onto nitrocellulose membranes. The membranes were blocked with 3% bovine serum albumin (BSA) and incubated overnight at 4°C with primary antibodies against LAMP2 (Abcam) and Cathepsin D (Abclonal). After three washes with TBST (15 min each), the membranes were incubated with HRP-conjugated goat anti-rabbit secondary antibodies. The membranes were washed three additional times (15 min each) to remove unbound secondary antibodies, and immunoreactive bands were detected using enhanced chemiluminescence (ECL, Thermo Scientific) and visualised using a chemiluminescence imaging system.

#### Phago-lysosome fusion assay

THP-1-derived macrophages were incubated with tetramethylrhodamine (TMR)- conjugated dextran (5 mg mL⁻¹) for 1 h to label lysosomes. Cells were subsequently washed extensively with PBS and incubated in fresh complete medium for an additional 2 h (chase period) to allow dextran accumulation within lysosomes. Cells were then treated with MEVs for 12hr, followed by incubation with Alexa Fluor 488-conjugated latex beads (1 µm diameter) for 120 min to allow phagosome formation. Live-cell time-lapse imaging was performed at 37°C using a temperature-controlled microscope stage equipped with a CO₂ incubation chamber. Fusion of lysosomes (red) with phagosomes containing Alexa Fluor 488-labeled latex beads (green) was visualized by the appearance of yellow fluorescence resulting from signal overlap.

For osmotic experiments, Rab5-transfected cells were loaded with fluorescent dextran to label endocytic compartments, and live-cell imaging was performed under isotonic, hypertonic, or hypotonic conditions. Hypertonic conditions were achieved by increasing the medium osmolality by 50 mOsm above isotonic, whereas hypotonic conditions were established by reducing the medium osmolality by 50 mOsm below isotonic through dilution. Rab5 recruitment dynamics to dextran-containing compartments were monitored by live-cell confocal microscopy under each osmotic condition.

For infection studies, macrophages were infected with GFP-*Mycobacterium smegmatis* for 4 h to allow bacterial internalisation and phagosome formation. Following infection, the cells were incubated with PKH26-labelled MEVs for 30 min at 37°C. Cells were subsequently washed with PBS, fixed with 4% paraformaldehyde (PFA) for 15 min at room temperature, and imaged using confocal microscopy.

#### In-Vivo fluorescence imaging

All in vivo experiments involved 6- to 8-week-old BALB/c mice. PKH-26 labelled MEVs were injected in BALB/c mice of 6–8 weeks of age via intranasal (IN) administration. After 48hours GFP-*M.smegmatis* bacteria were injected via intranasal administration. After 7^th^ day, the mice were sacrificed, the organs were resected, and fluorescence images of the organs were captured using NEWTON FT–30 Fluorescence and Bioluminescence Imaging System.

## Ethical statement

All animal experiments were conducted in accordance with the guidelines of the Committee for the Purpose of Control and Supervision of Experiments on Animals (CPCSEA) (Registration No. 1634/GO/ReRcBiBt/S/12/CPCSEA. The experimental protocols were approved by the Institutional Animal Ethics Committee (IAEC) of the National Institute of Science Education and Research (NISER), Bhubaneswar, India, and all procedures were performed in compliance with institutional and national ethical guidelines (Ethical Approval No. NISER/SBS/AH-355).

## Image & Statistical analysis

All data are representative of at least three independent experiments (n = 3), unless otherwise specified in the figure legends. Image analysis and processing were performed using ImageJ. Flow cytometry data were analysed using BD FACS DIVA (BD Biosciences, USA). Data normality was assessed using the Shapiro-Wilk test before statistical analysis. Statistical significance was assessed using one-way ANOVA with Tukey’s test, with p < 0.05 considered significant. All statistical analyses were performed using OriginPro software and SigmaPlot.

## Acknowledgements

We thank Sandeep Choubey, Swagata Ghatak and Bibhu Ranjan Sarangi for critical comments on the manuscript. We thank Rohan Dhiman for the kind gift of the *M. smegmatis and M. tb H37Ra strains*. We thank Apurba Koner for the JIND-Mor probe. We acknowledge all the members of the M.S. lab for their critical reading of the manuscript. MS gratefully acknowledges extramural support from the DBT–Wellcome Trust India Alliance Intermediate Fellowship (Grant IA/I/20/2/505212), the Department of Science and Technology-SERB (Grant EMR/2017/004513) and the Department of Biotechnology (DBT), Government of India (BT/PR21226/MED/122/41/2016 and BT/INF/22/SP33046/2019) and intramural funding from the Department of Atomic Energy. SK gratefully acknowledges support from the DBT-Wellcome Trust India Alliance Intermediate Fellowship (Grant IA/I/21/1/505624).

