## Supplementary dataset for "Pathogenic Bacteria Remotely Remodel Host Membrane Mechanics via Extracellular Vesicles to subvert Phagosome Maturation and Promote infection"

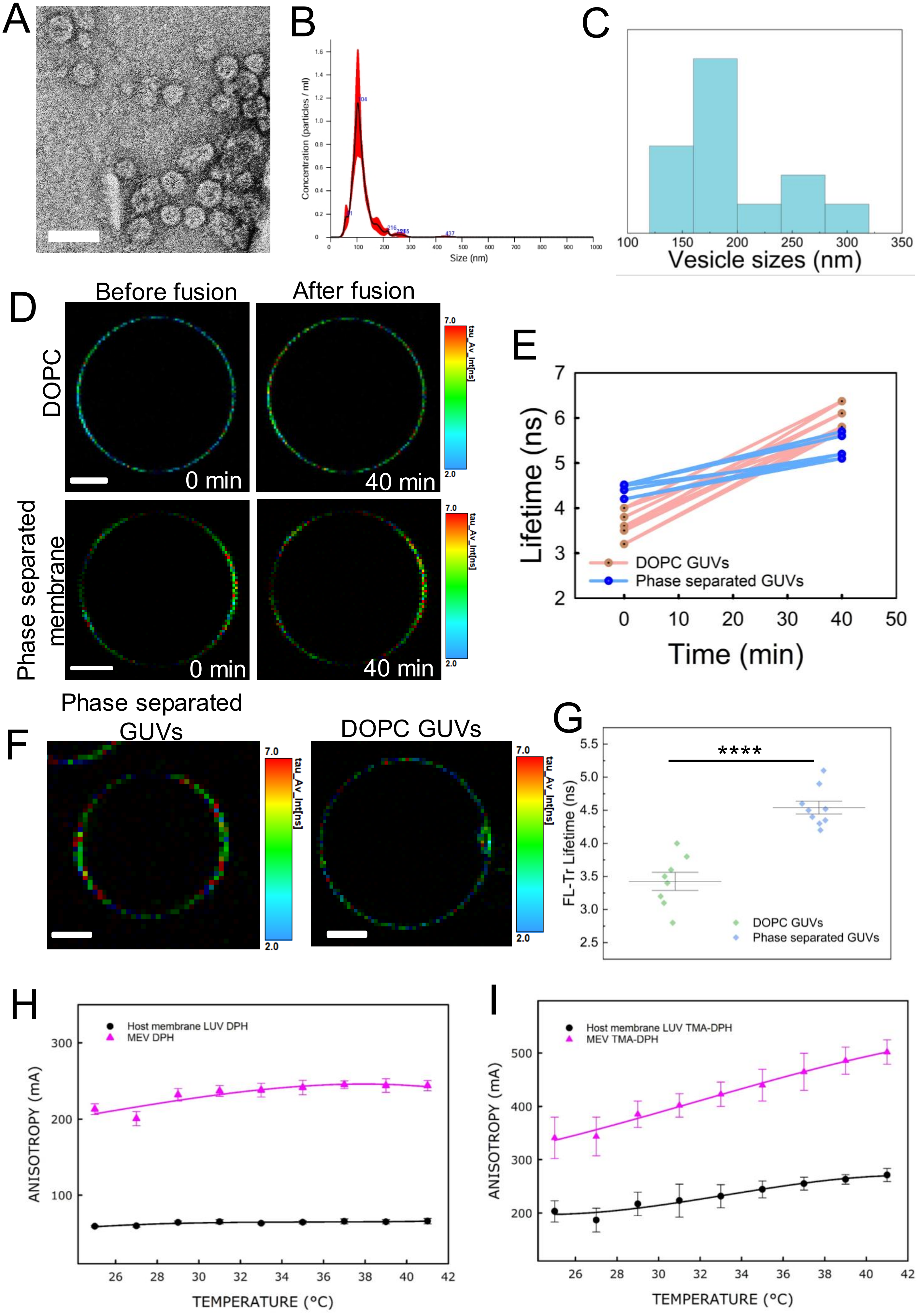

A
B
Concentration (particles / ml)
Size (nm)
C
Vesicle sizes (nm)
D
Before fusion
After fusion
DOPC
Phase separated membrane
0 min
40 min
tau_Av_Int[ns]
7.0
2.0
E
Lifetime (ns)
Time (min)
DOPC GUVs
Phase separated GUVs
F
Phase separated GUVs
DOPC GUVs
G
FL-Tr Lifetime (ns)
****
DOPC GUVs
Phase separated GUVs
H
Host membrane LUV DPH
MEV DPH
ANISOTROPY (mA)
TEMPERATURE (°C)
I
Host membrane LUV TMA-DPH
MEV TMA-DPH
ANISOTROPY (mA)
TEMPERATURE (°C)

**Fig.S1.**
**(A)**Ultra-high-resolution imaging of isolated MEVs confirms their uniform spherical morphology. Scale bar, 200 nm.

**(B)**Nanoparticle tracking analysis of MEV was done using NTA version 3.4. Samples were diluted up to an estimated concentration of $10^{8}$–$10^{9}$ vesicles $ml^{-1}$ to match NTA's compatibility.

**(C)** Dynamic light scattering (DLS) analysis showing the size distribution profile of isolated MEVs, demonstrating a predominantly nanoscale vesicle population (major peak ~150- 200 nm).

**(D)** FLIM images of Flipper-TR incorporated DOPC GUVs and phase-separated GUVs (PC:SM: CHOL-3:3:4) before and after exposure to DOPC LUVs. Scale bar: 5 µm

**(E)** Corresponding fluorescence lifetime measurements of GUVs as shown in (a). Data points represent mean ± S.D. from each independent experiment (n~20GUVs at each condition).

**(F)** FLIM images of Flipper-TR incorporated DOPC GUVs (Homogenous membrane) and phase-separated GUVs (PC:SM: CHOL-3:3:4). Scale bar: 5 µm.

**(G)** Corresponding fluorescence lifetime measurements of GUVs as shown in (c). Data points represent mean ± S.D. from each independent experiment (n~30GUVs at each condition). (**p<0.01, ***p<0.001 in one-way ANOVA).

Fluorescence anisotropy of Mycobacterial extracellular vesicles and host membrane as a function of time.
**(H)** DPH (probes the hydrophobic region of the lipid bilayer)

**(I)** TMA-DPH (probes the hydrophilic region of the lipid bilayer). Data points represent at least three independent experiments with mean ± S.D.

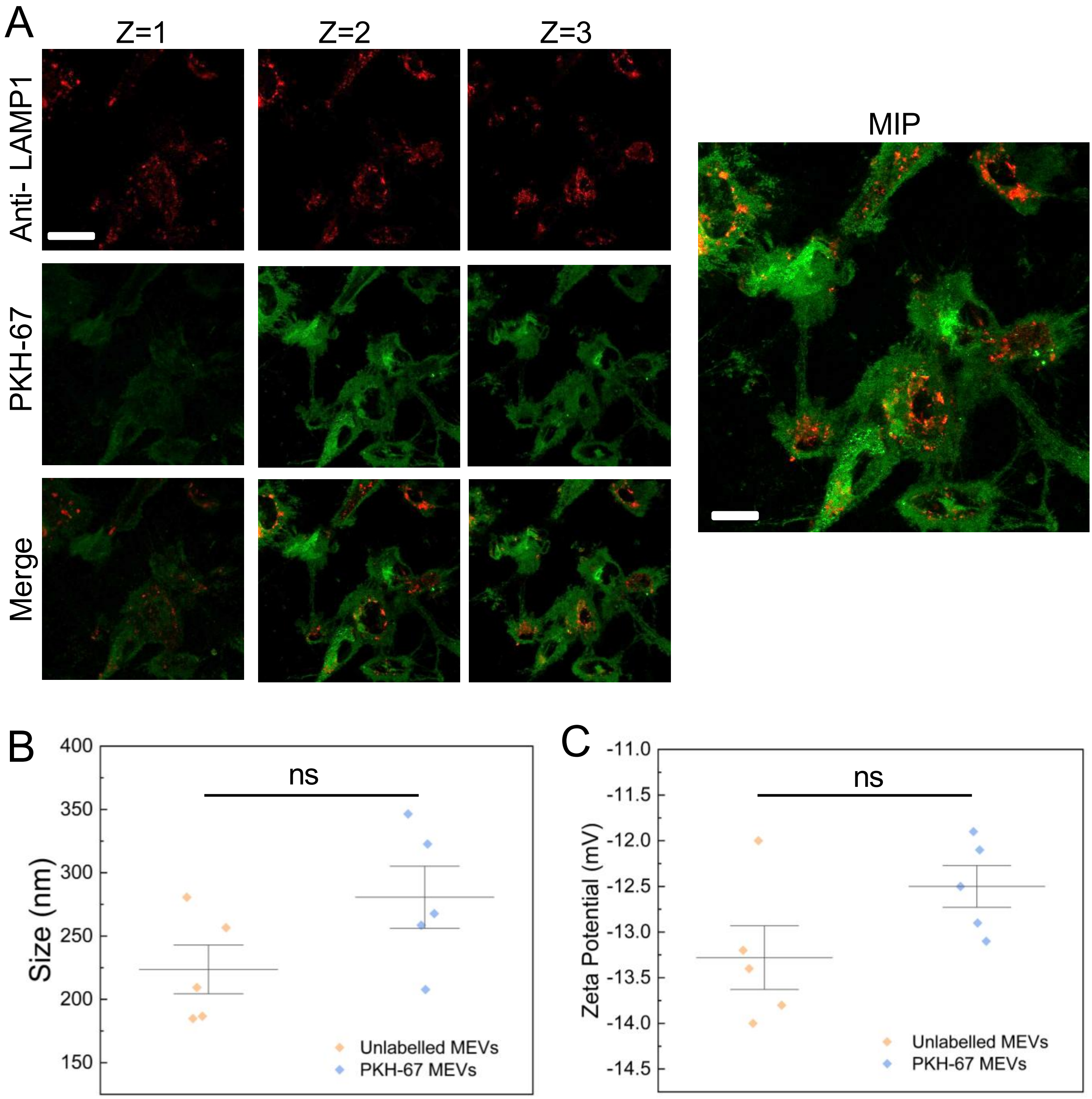

A
Z=1
Z=2
Z=3
Anti- LAMP1
PKH-67
Merge
MIP
B
Size (nm)
400
350
300
250
200
150
ns
Unlabelled MEVs
PKH-67 MEVs
C
Zeta Potential (mV)
-11.0
-11.5
-12.0
-12.5
-13.0
-13.5
-14.0
-14.5
ns
Unlabelled MEVs
PKH-67 MEVs

**Fig. S2.**

**(A)** Z-stack confocal imaging of THP-1–derived macrophages treated with only PKH67 dye and labelled lysosomes with Anti-LAMP1. The PKH-67 dye seems to be diffuse completely inside the cells and colocalises with lysosomes within 30 min of treatment. These are representative images from three independent replicates. Scale bar, 10 μm

**(B)** Size distribution analysis and **(C)** zeta potential analysis of PKH-labelled and unlabelled MEVs. Data points represent the mean of each biological replicate.

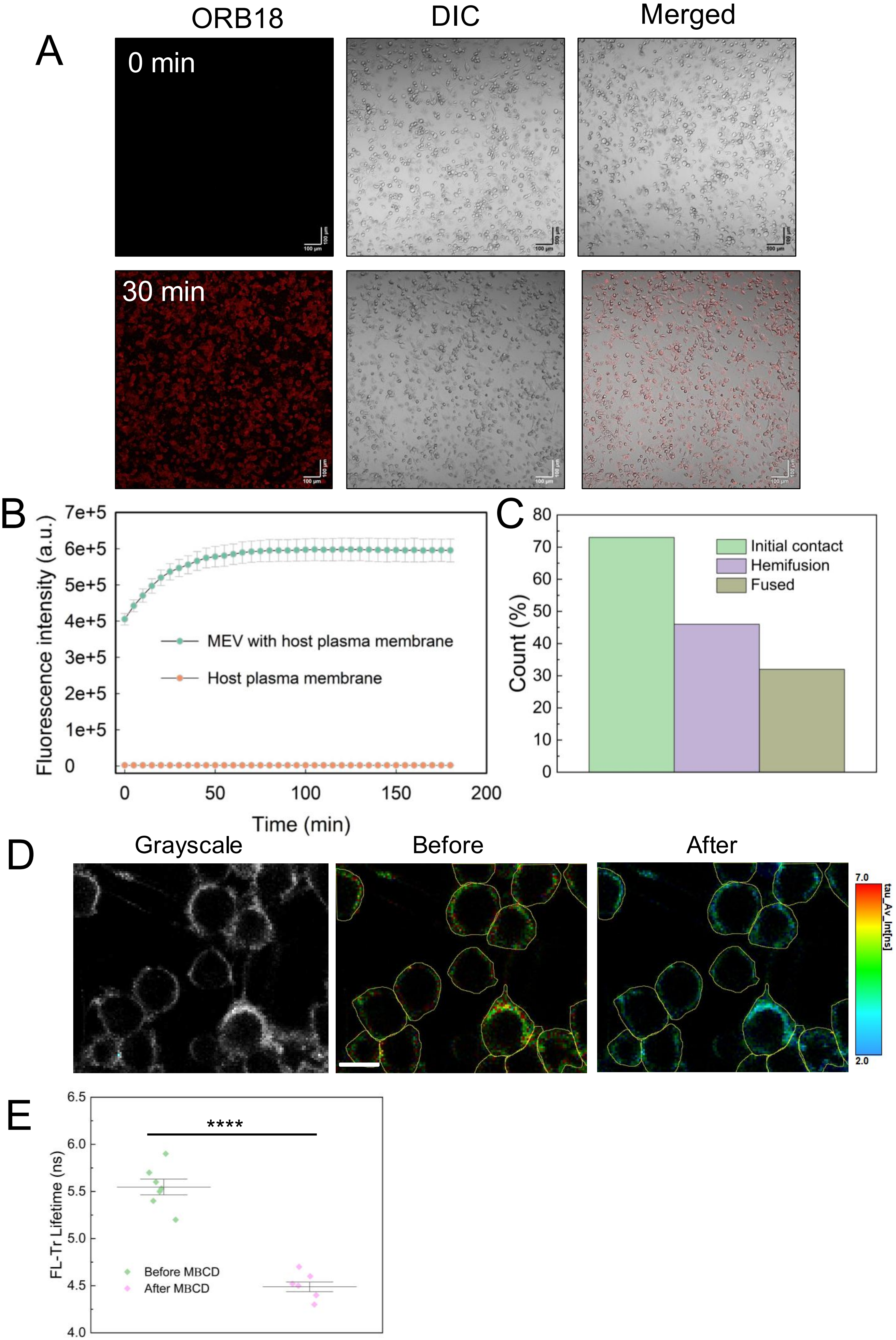
ORB18
DIC
Merged
A
0 min
30 min
100 μm
B
Fluorescence intensity (a.u.)
7e+5
6e+5
5e+5
4e+5
3e+5
2e+5
1e+5
0
MEV with host plasma membrane
Host plasma membrane
0
50
100
150
200
Time (min)
C
Count (%)
80
70
60
50
40
30
20
10
0
Initial contact
Hemifusion
Fused
D
Grayscale
Before
After
7.0
2.0
tau_Av_Int[ns]
E
FL-Tr Lifetime (ns)
6.5
6.0
5.5
5.0
4.5
4.0
****
Before MBCD
After MBCD

**Fig. S3.**

**(A)**Time-lapse confocal imaging of THP-1–derived macrophages incubated with R18-labelled MEVs for 30 minutes. Representative images showing progressive MEV uptake dynamics over the 30 minutes. Scale bar, 100 µm. Images are representative of five independent biological experiments.

**(B)** R18 dequenching assay of plasma membrane mimicking LUVs labelled with self-quenching lipophilic dye R18, and fluorescence intensity was monitored as a function of time in the absence and presence of MEV. Data points are shown as the means ± S.D. of three independent measurements. All experiments are carried out in PBS pH 7.4 at 37°C.

**(C)** Size distribution of the stages of the MEV fusion with the plasma membrane mimicking LUV in fig 1g.

**(D)** FLIM images of Flipper-Tr-incorporated plasma membrane before and after exposure to MβCD.

**(E)** Corresponding fluorescence lifetime measurements of macrophages as shown in (D). Data points shown represents mean ± S.D. from each independent experiment. (**$p<0.01$, ***$p<0.001$ in one-way ANOVA).

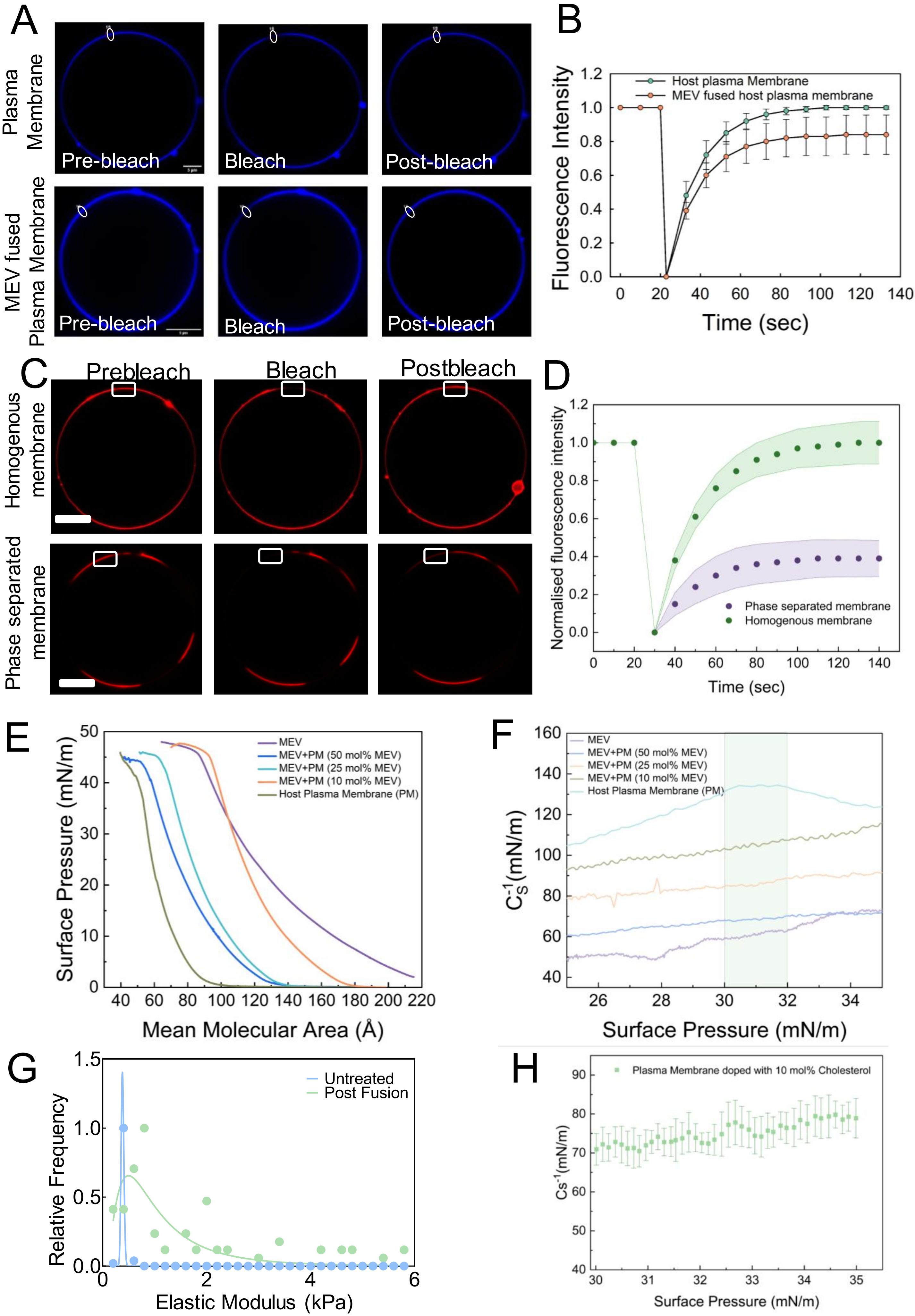
A
Plasma Membrane
MEV fused Plasma Membrane
Pre-bleach
Bleach
Post-bleach
B
Fluorescence Intensity
Time (sec)
Host plasma Membrane
MEV fused host plasma membrane
C
Prebleach
Bleach
Postbleach
Homogenous membrane
Phase separated membrane
D
Normalised fluorescence intensity
Time (sec)
Phase separated membrane
Homogenous membrane
E
Surface Pressure (mN/m)
Mean Molecular Area (Å)
MEV
MEV+PM (50 mol% MEV)
MEV+PM (25 mol% MEV)
MEV+PM (10 mol% MEV)
Host Plasma Membrane (PM)
F
$C_S^{-1}$(mN/m)
Surface Pressure (mN/m)
MEV
MEV+PM (50 mol% MEV)
MEV+PM (25 mol% MEV)
MEV+PM (10 mol% MEV)
Host Plasma Membrane (PM)
G
Relative Frequency
Elastic Modulus (kPa)
Untreated
Post Fusion
H
Cs-1(mN/m)
Surface Pressure (mN/m)
Plasma Membrane doped with 10 mol% Cholesterol

**Fig. S4.**

**(A)** Representative FRAP images of plasma membrane mimicking GUVs before and after MEV fusion, showing differences in post-bleach recovery. Scale bar: 5 μm.

**(B)** Mean fluorescence recovery curves comparing MEV-fused and untreated (only PBS treated) GUV membranes. Data points represent mean values from three independent experiments. Langmuir monolayer analysis of concentration-dependent MEV fusion effects on phagosomal membrane.

**(C)** Representative FRAP images of Homogenous membrane and phase-separated membrane. Scale bar: 5 μm.

**(D)** Mean fluorescence recovery curves where data points represent mean values from three independent experiments. (n~15 GUVs in each condition).

**(E)** Surface pressure (π)-mean molecular area (A) isotherms of phagosomal membrane monolayers incorporating increasing mol% of MEVs.

**(F)** Compressibility modulus ($Cs^{-1}$) values quantified at a surface pressure of 30–32 mN/m (bilayer-equivalent pressure, indicated by the shaded region) for the conditions shown in (g). All monolayer measurements were conducted on an autoclaved dd$H_2O$ subphase at 25 °C. Data represent mean values from three independent experiments.

**(G)** Normalised distribution of dTHP-1 macrophage surfaces measured by atomic force microscopy (AFM) show an increase in cellular stiffness following MV fusion relative to untreated (only PBS) macrophages. Data are presented as mean ± SEM from ≥ 60 cells per condition.

**(H)** Compressibility modulus ($Cs^{-1}$) values quantified at a surface pressure of 30–32 mN/m for the plasma membrane mimicking membrane doped with 10mol% of cholesterol. All monolayer measurements were conducted on an autoclaved dd$H_2O$ subphase at 25 °C. Data represent mean values from three independent experiments.

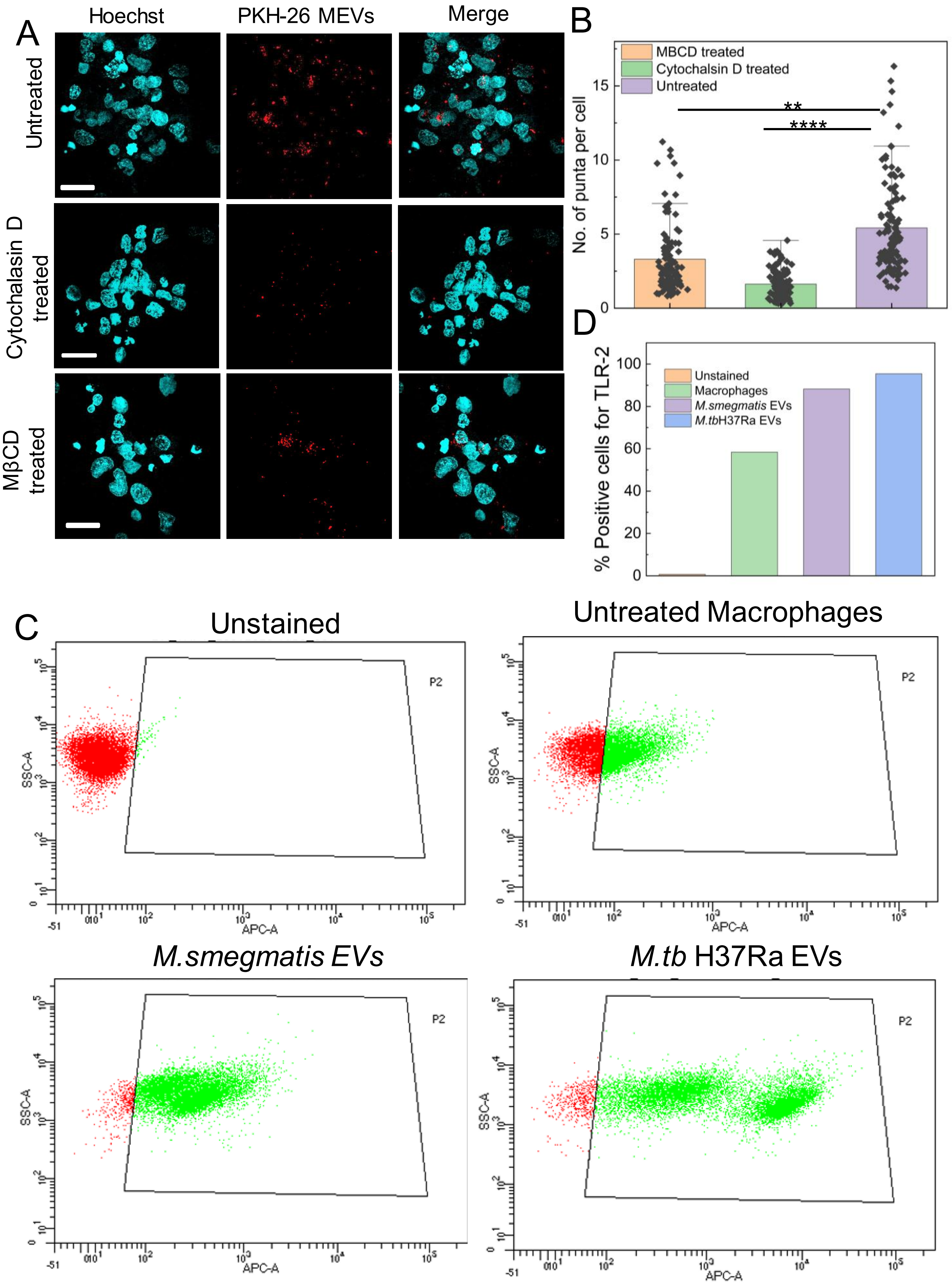

A
Hoechst
PKH-26 MEVs
Merge
Untreated
Cytochalasin D treated
MβCD treated
B
MBCD treated
Cytochalasin D treated
Untreated
No. of punta per cell
**
****
D
% Positive cells for TLR-2
Unstained
Macrophages
M.smegmatis EVs
M.tbH37Ra EVs
C
Unstained
Untreated Macrophages
M.smegmatis EVs
M.tb H37Ra EVs
P2
SSC-A
APC-A

**Fig. S5.**

**(A)** Confocal images of the fixed macrophages pre-treated with Cytochalasin D and MβCD followed by the MEV treatment. These are representative images from three independent experiments.

**(B)** Quantification of the number of puncta per cell corresponding to the conditions. (**$p<0.01$, ***$p<0.001$ in one-way ANOVA).

**(C)** Flow cytometric analysis of TLR2 surface expression in THP-1–derived macrophages following MEV treatment. Flow cytometry profiles showing an increase in TLR2 surface expression upon exposure to MEVs compared with untreated (only PBS) macrophages.

**(D)** Bar plots represent mean fluorescence intensity (MFI), indicating TLR2 upregulation. Data are representative of three independent biological experiments.

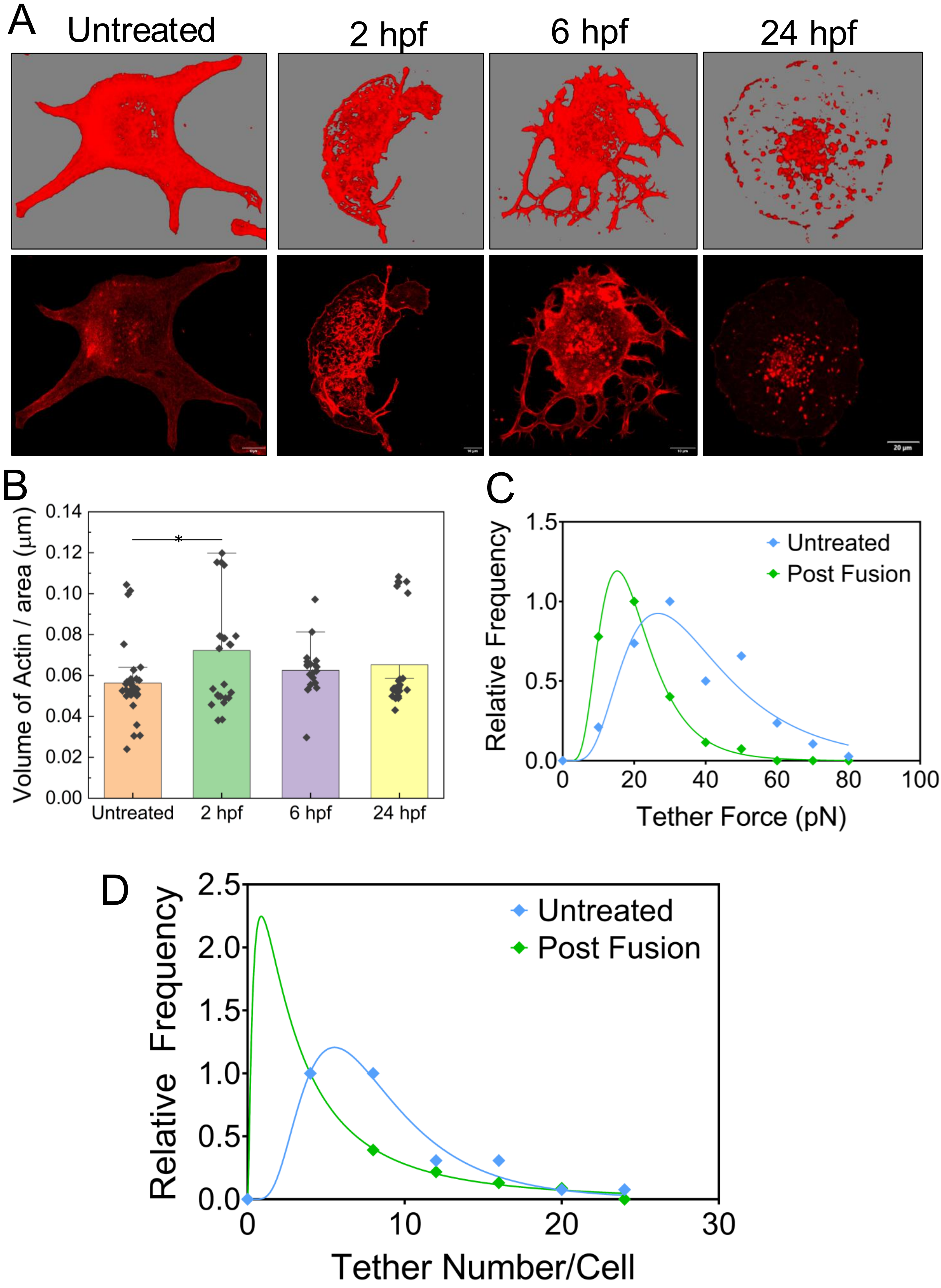
A
Untreated
2 hpf
6 hpf
24 hpf
20 μm
B
Volume of Actin / area (μm)
0.14
0.12
0.10
0.08
0.06
0.04
0.02
0.00
*
Untreated
2 hpf
6 hpf
24 hpf
C
Relative Frequency
1.5
1.0
0.5
0.0
0
20
40
60
80
100
Tether Force (pN)
Untreated
Post Fusion
D
Relative Frequency
2.5
2.0
1.5
1.0
0.5
0.0
0
10
20
30
Tether Number/Cell
Untreated
Post Fusion

**Fig.S6.**
**(A)** Three-dimensional (3D) isosurface renderings (top) and corresponding two-dimensional (2D) confocal projections (bottom) of dTHP-1 cells show increased surface complexity and membrane protrusions after MEVs fusion.

**(B)**The quantified surface volume confirms significant morphological expansion compared to control. Scale bars: 20 μm. Data are shown as mean ± SEM; statistical significance determined using one-way ANOVA.

**(C)** Normalised distribution of tether-force measurements reveals altered membrane-cytoskeleton coupling following fusion, as reflected by changes in the force required to extract membrane tethers from the plasma membrane.

**(D)** Normalised distribution of tether number per cell reveals membrane deformability and cell cortex tension upon MEV fusion. Data are presented as mean ± SEM from ≥ 60 cells per condition.

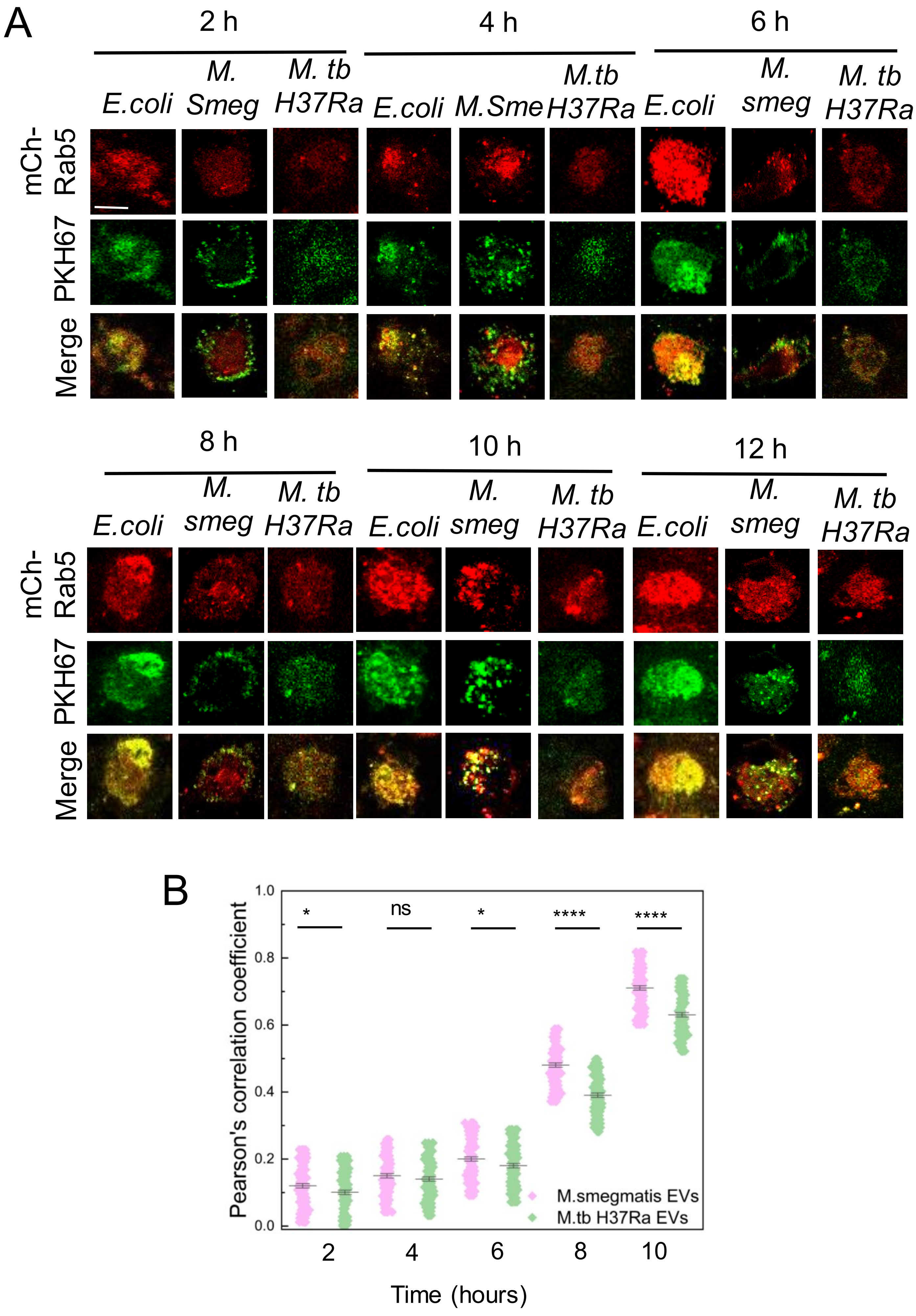
A
2 h
4 h
6 h
E.coli
M. Smeg
M. tb H37Ra
M.Sme
M.tb H37Ra
M. smeg
mCh-Rab5
PKH67
Merge
8 h
10 h
12 h
B
Pearson's correlation coefficient
1.0
0.8
0.6
0.4
0.2
0.0
*
ns
*
****
****
M.smegmatis EVs
M.tb H37Ra EVs
2
4
6
8
10
Time (hours)

**Fig. S7.**

**(A)** Time-lapse confocal imaging of THP-1–derived macrophages incubated with PKH67-labelled MEVs.Representative images of fixed macrophages showing the delayed recruitment of RAB5 to MEV-containing phagosomes over time. Scale bar, 10 μm. Images are representative of five independent biological experiments.

**(B)** Data points represent the quantitative colocalization analysis of Rab5 recruitment at multiple time points using Pearson's correlation coefficient. (n=75 cells were quantified for each condition at a single time point). (**p<0.01, ***p<0.001 in one-way ANOVA).

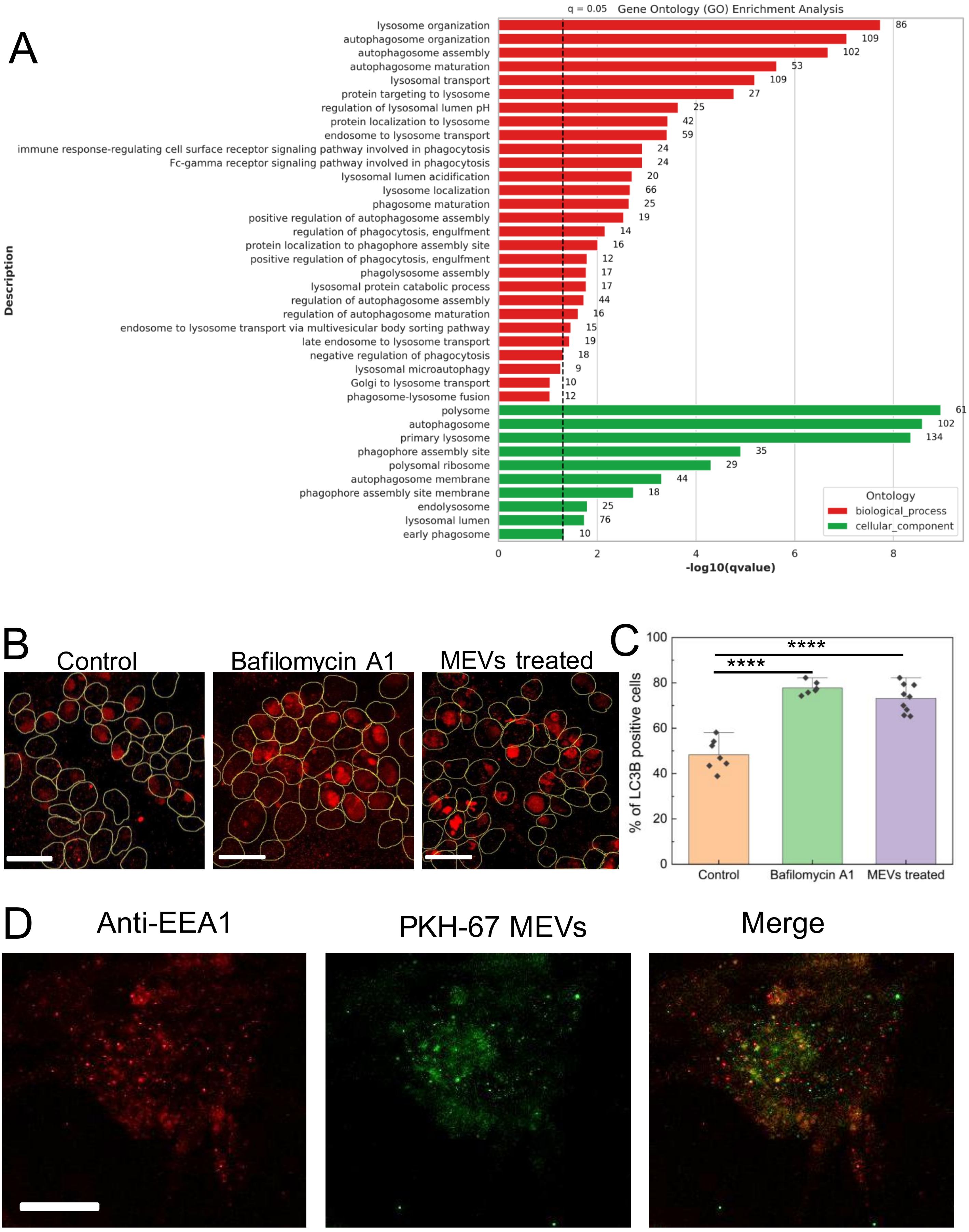
A
q = 0.05 Gene Ontology (GO) Enrichment Analysis
lysosome organization 86
autophagosome organization 109
autophagosome assembly 102
autophagosome maturation 53
lysosomal transport 109
protein targeting to lysosome 27
regulation of lysosomal lumen pH 25
protein localization to lysosome 42
endosome to lysosome transport 59
immune response-regulating cell surface receptor signaling pathway involved in phagocytosis 24
Fc-gamma receptor signaling pathway involved in phagocytosis 24
lysosomal lumen acidification 20
lysosome localization 66
phagosome maturation 25
positive regulation of autophagosome assembly 19
regulation of phagocytosis, engulfment 14
protein localization to phagophore assembly site 16
positive regulation of phagocytosis, engulfment 12
phagolysosome assembly 17
lysosomal protein catabolic process 17
regulation of autophagosome assembly 44
regulation of autophagosome maturation 16
endosome to lysosome transport via multivesicular body sorting pathway 15
late endosome to lysosome transport 19
negative regulation of phagocytosis 18
lysosomal microautophagy 9
Golgi to lysosome transport 10
phagosome-lysosome fusion 12
polysome 61
autophagosome 102
primary lysosome 134
phagophore assembly site 35
polysomal ribosome 29
autophagosome membrane 44
phagophore assembly site membrane 18
endolysosome 25
lysosomal lumen 76
early phagosome 10
Description
Ontology
biological_process
cellular_component
0 2 4 6 8
-log10(qvalue)
B
Control
Bafilomycin A1
MEVs treated
C
****
****
% of LC3B positive cells
0 20 40 60 80 100
Control
Bafilomycin A1
MEVs treated
D
Anti-EEA1
PKH-67 MEVs
Merge

**Fig. S8.**

**(A)** Gene ontology (GO) enrichment analysis of whole-genome transcriptomic profiling of macrophages following MEV treatment. MEV exposure leads to a transcriptional reprogramming of host cells, with significant enrichment of pathways associated with vesicle trafficking, membrane dynamics, and phagosome-lysosome fusion. These changes highlight the broad impact of MEVs on host endomembrane regulation.

**(B)** Immunofluorescence analysis of LC3B in untreated (only PBS treated), bafilomycin-treated, and drug-treated cells. Representative confocal images showing LC3B puncta (red) in untreated macrophages, macrophages treated with bafilomycin A1 (autophagy flux inhibitor), and MEV-treated macrophages. Scale bar, 20 µm.

**(C)** Quantification of %of LC3B positive cell (>20 puncta per cell) is shown in the right panel, expressed as mean ± SD from n = 3 independent experiments. (**p<0.01, ***p<0.001 in one-way ANOVA).

**(D)** Representative confocal images of macrophages showing colocalization of PKH-67 MEVs with EEA1 (Rab5 effector protein), detected by immunofluorescence staining using an anti-EEA1 antibody. Images are representative of three independent experiments. Scale bar, 10 µm

A

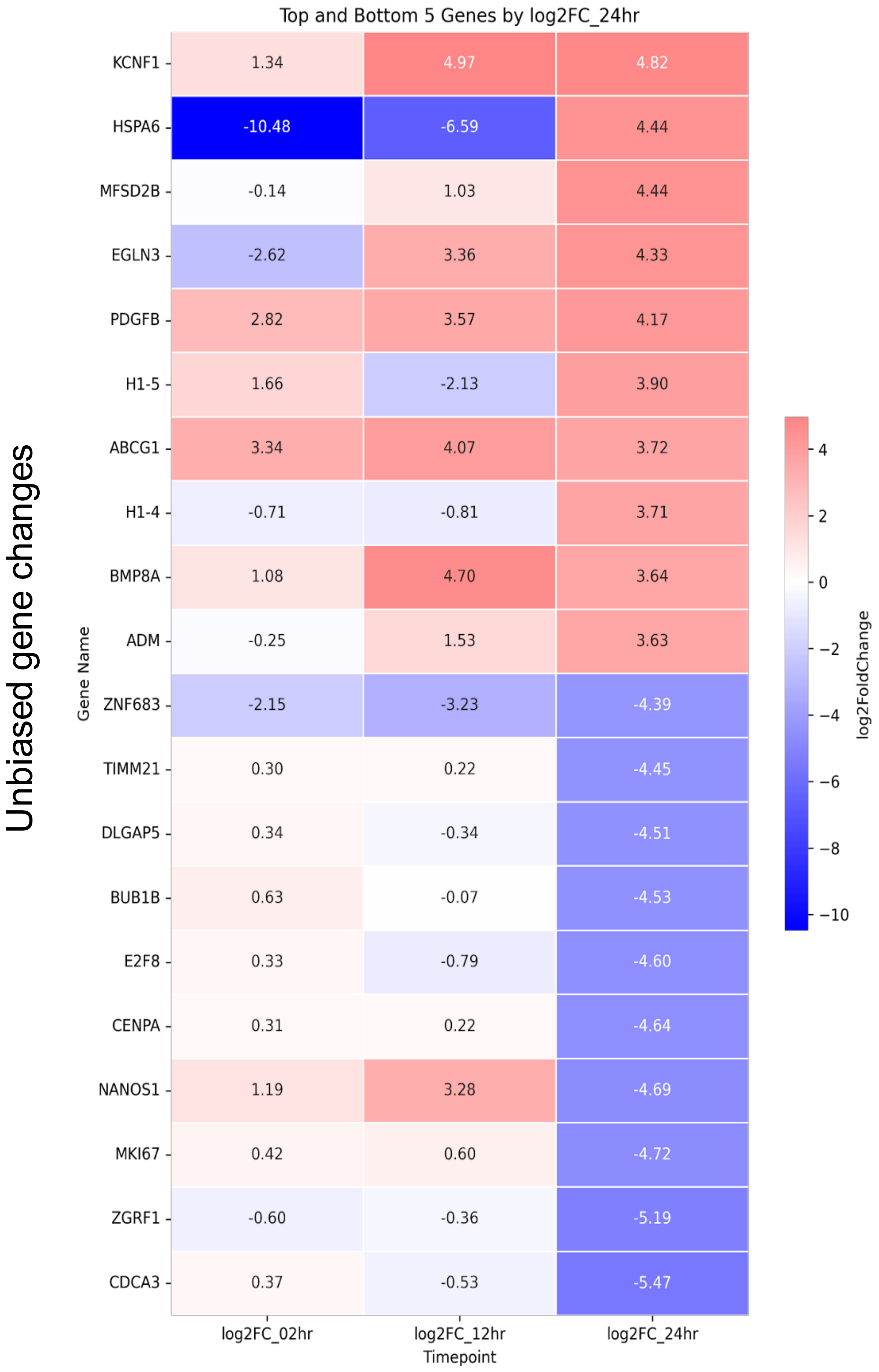
Unbiased gene changes
Top and Bottom 5 Genes by log2FC_24hr
Gene Name
KCNF1 1.34 4.97 4.82
HSPA6 -10.48 -6.59 4.44
MFSD2B -0.14 1.03 4.44
EGLN3 -2.62 3.36 4.33
PDGFB 2.82 3.57 4.17
H1-5 1.66 -2.13 3.90
ABCG1 3.34 4.07 3.72
H1-4 -0.71 -0.81 3.71
BMP8A 1.08 4.70 3.64
ADM -0.25 1.53 3.63
ZNF683 -2.15 -3.23 -4.39
TIMM21 0.30 0.22 -4.45
DLGAP5 0.34 -0.34 -4.51
BUB1B 0.63 -0.07 -4.53
E2F8 0.33 -0.79 -4.60
CENPA 0.31 0.22 -4.64
NANOS1 1.19 3.28 -4.69
MKI67 0.42 0.60 -4.72
ZGRF1 -0.60 -0.36 -5.19
CDCA3 0.37 -0.53 -5.47
log2FC_02hr
log2FC_12hr
log2FC_24hr
Timepoint
log2FoldChange
4
2
0
−2
−4
−6
−8
−10

**Fig. S9.**

**(A)** Heatmaps showing most affected gene expression profiles of macrophage genes at different time intervals (2,12,24 hours) following MEV treatment.

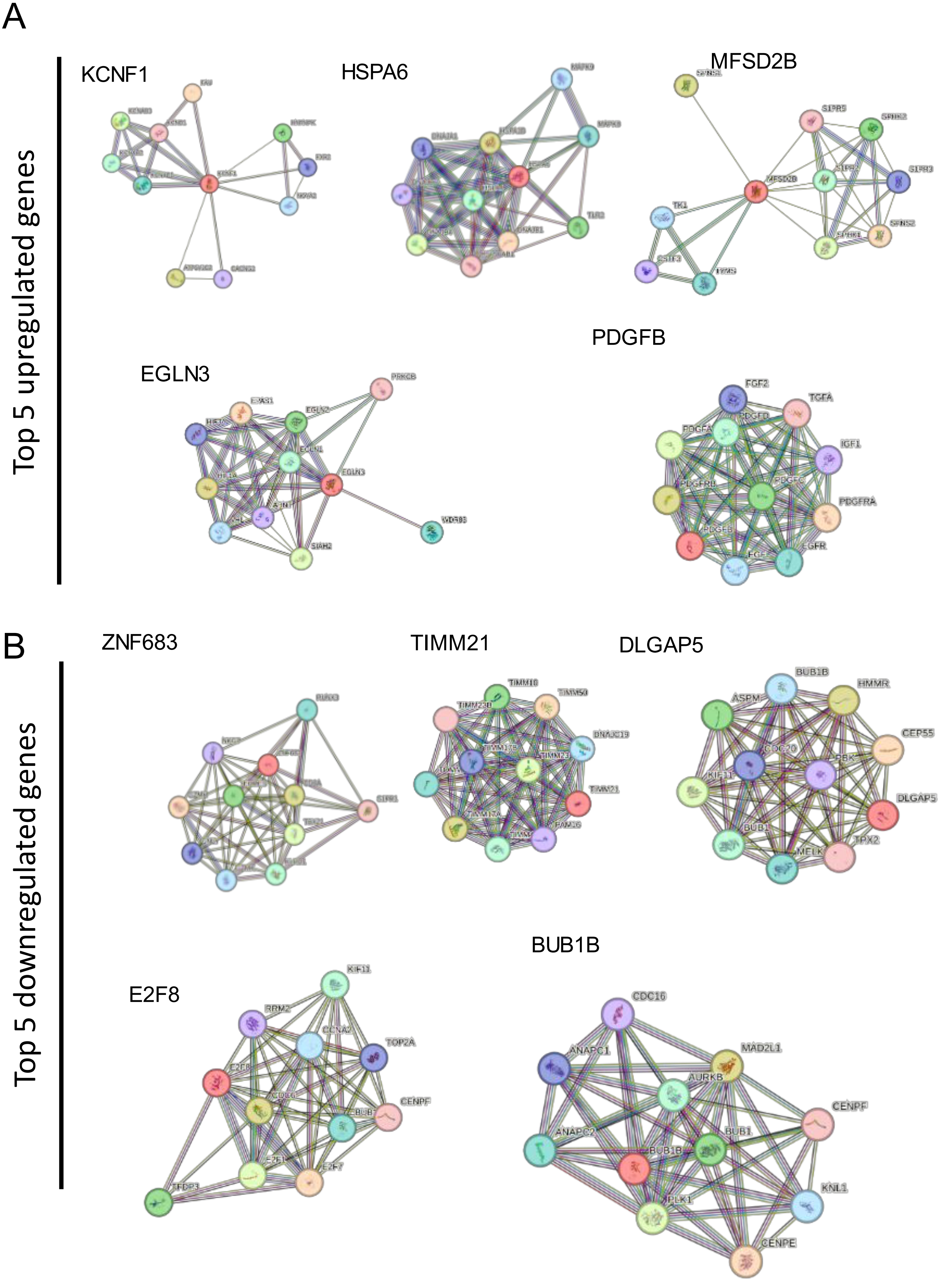
A
Top 5 upregulated genes
KCNF1
HSPA6
MFSD2B
EGLN3
PDGFB
FGF2
TGFA
PDGFD
PDGFA
IGF1
PDGFRB
PDGFC
PDGFRA
PDGFB
EGFR
B
Top 5 downregulated genes
ZNF683
TIMM21
DLGAP5
BUB1B
ASPM
HMMR
CEP55
CDC20
PBK
KIF11
DLGAP5
BUB1
MELK
TPX2
E2F8
KIF11
RRM2
CCNA2
TOP2A
E2F8
CENPF
E2F1
E2F7
TFDP3
BUB1B
CDC16
ANAPC1
MAD2L1
AURKB
CENPF
ANAPC2
BUB1
BUB1B
PLK1
KNL1
CENPE

**Fig. S10.**

Signalling cascades associated with the **(A)** top five upregulated and **(B)** top five downregulated genes identified from the unbiased transcriptomic heatmap analysis. This highlights the major cellular pathways most significantly affected in macrophages following MEV treatment, revealing extensive transcriptional reprogramming of host signalling networks.

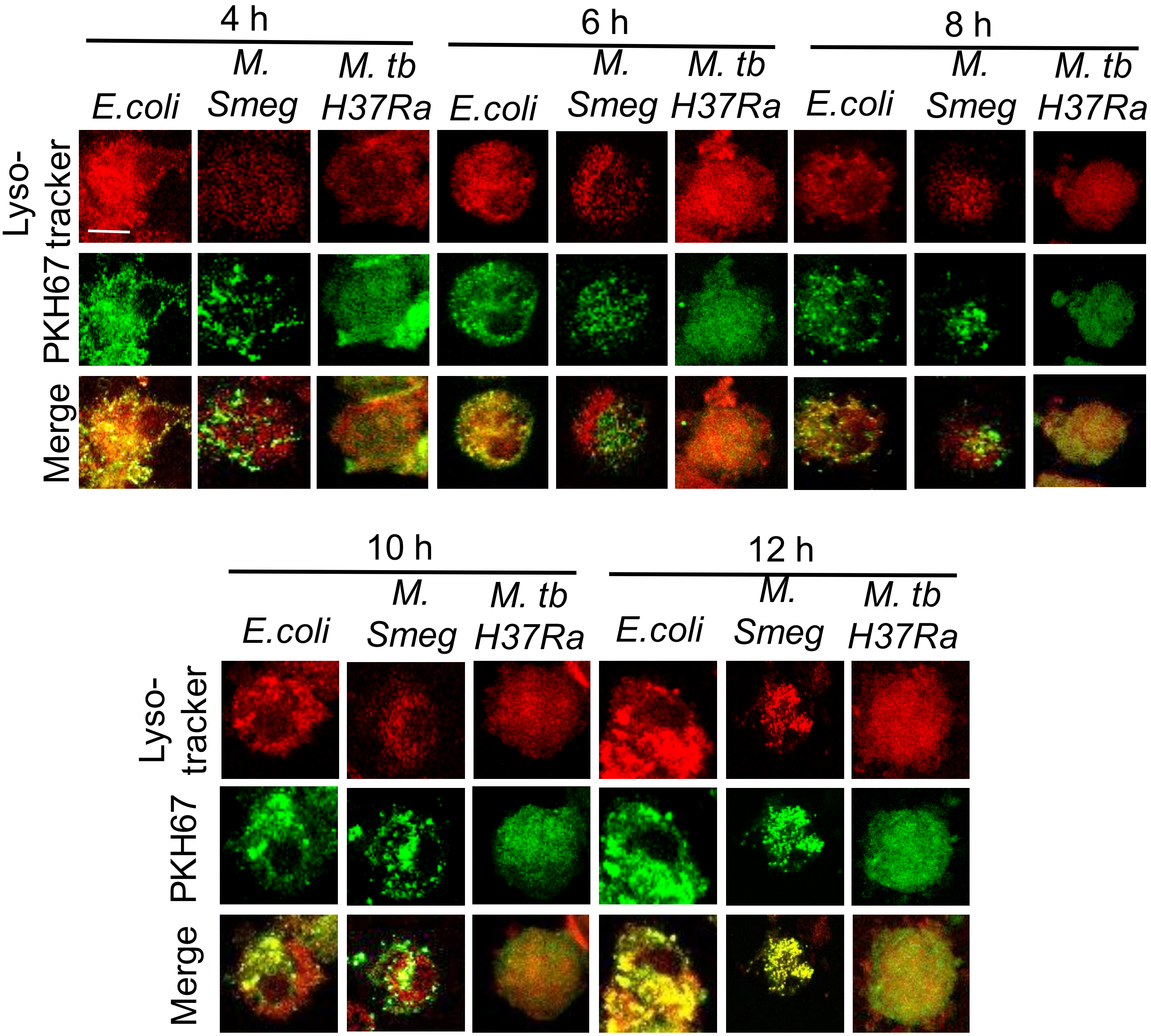

4 h
6 h
8 h
10 h
12 h
E.coli
M. Smeg
M. tb H37Ra
Lyso- tracker
PKH67
Merge

**Fig. S11.**

Time-lapse confocal imaging of THP-1–derived macrophages incubated with PKH67-labelled MEVs and lysotracker Red. Representative images of fixed macrophages showing the delayed fusion of MEV-containing phagosomes with lysosomes over time. Scale bar, 10 µm. Images are representative of five independent biological experiments.

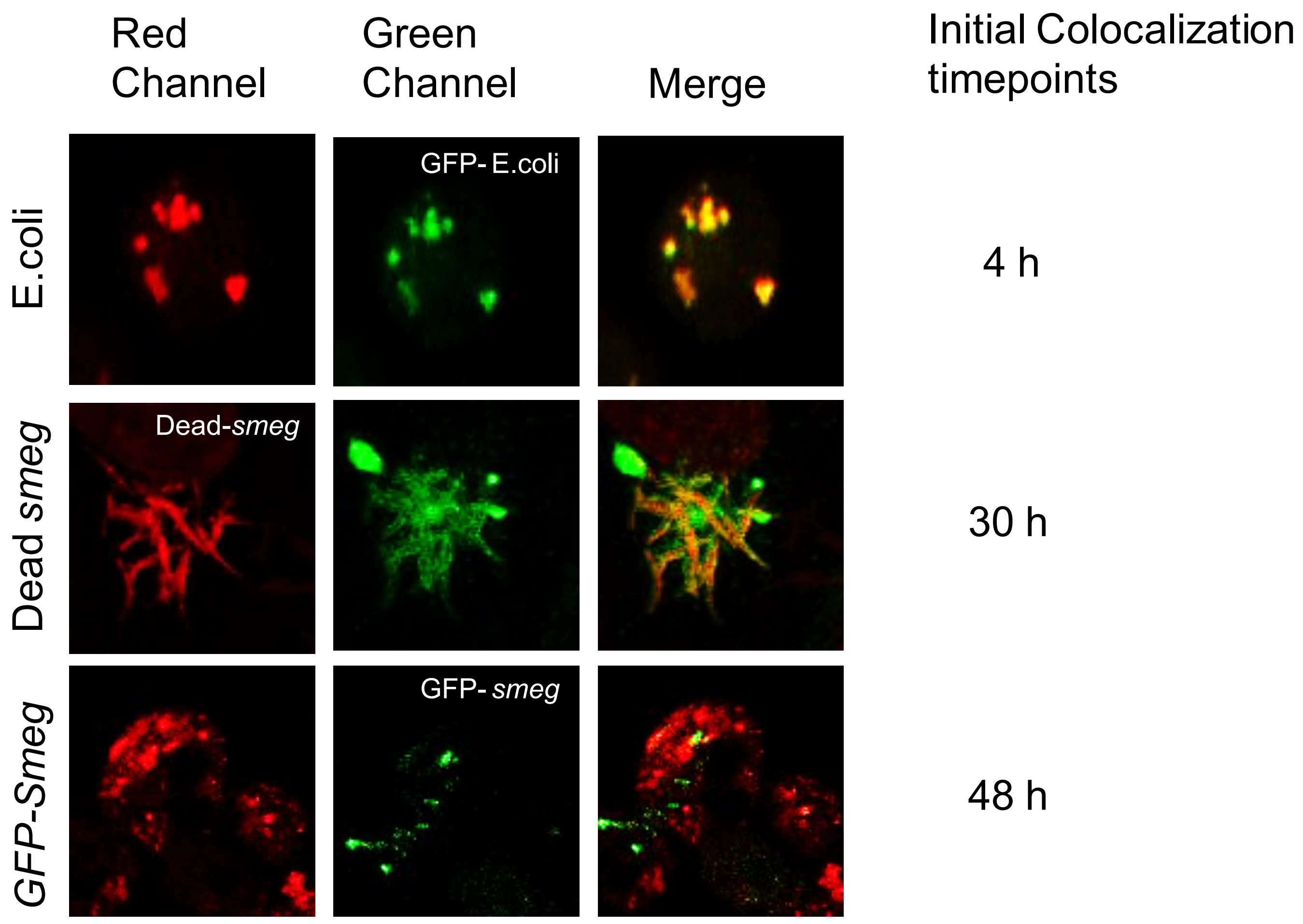

Red Channel
Green Channel
Merge
Initial Colocalization timepoints
E.coli
GFP- E.coli
4 h
Dead smeg
Dead-smeg
30 h
GFP-Smeg
GFP- smeg
48 h

**Fig. S12.**

Fixed-cell imaging of THP-1–derived macrophages infected with *E. coli*, GFP-*M. smegmatis*, or heat-killed *M. smegmatis*. Colocalisation with lysosomal markers is observed at 4 h for *E. coli*, 48 h for live GFP-*M. smegmatis*, and 30 h for heat-killed *M. smegmatis*. The delayed lysosomal fusion in live *M. smegmatis*–infected macrophages highlights the role of MEVs in impairing phagosome–lysosome fusion. Representative images from at least three independent biological experiments.

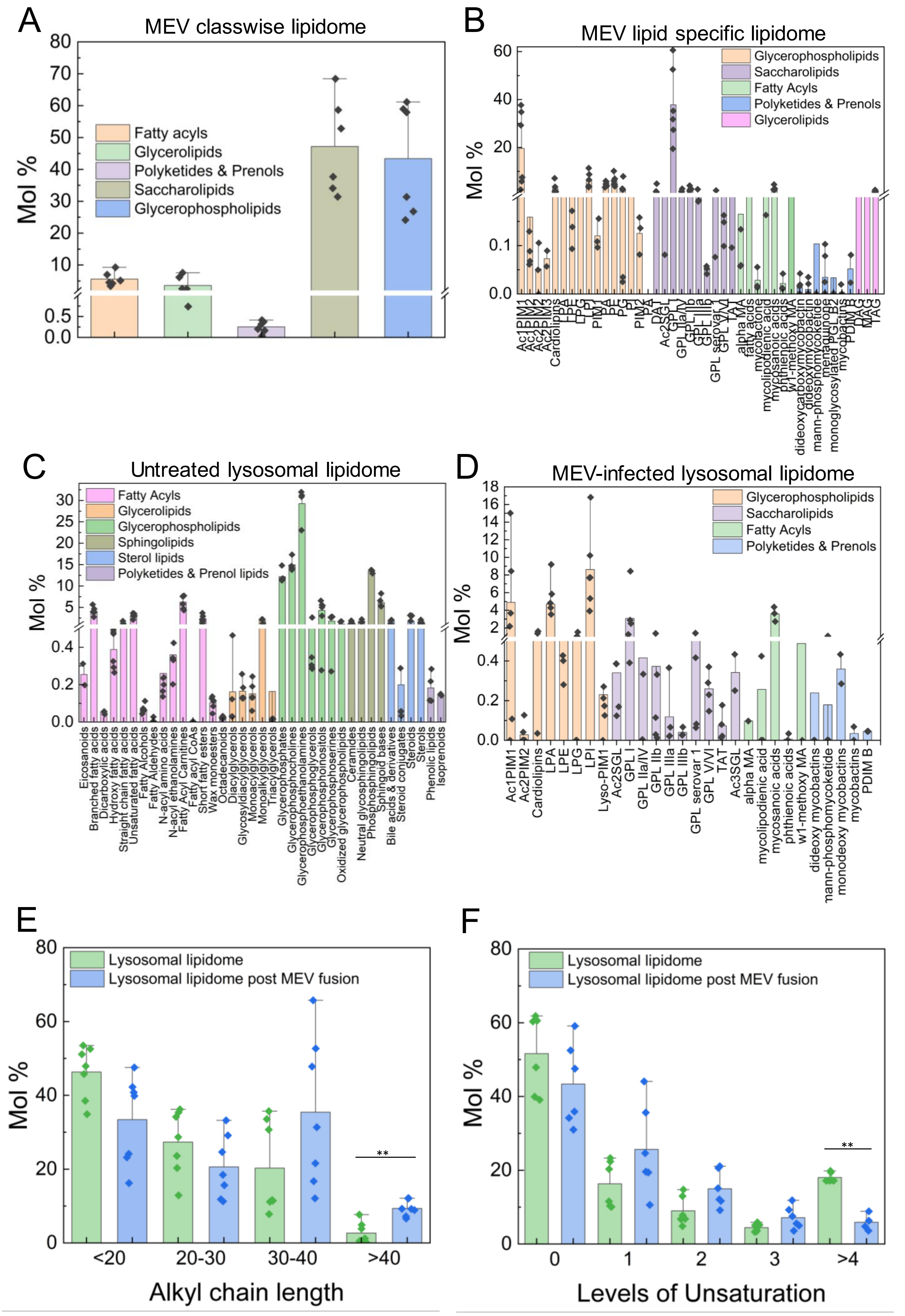
A
MEV classwise lipidome
Mol %
Fatty acyls
Glycerolipids
Polyketides & Prenols
Saccharolipids
Glycerophospholipids
B
MEV lipid specific lipidome
Mol %
Glycerophospholipids
Saccharolipids
Fatty Acyls
Polyketides & Prenols
Glycerolipids
Ac1PIM1
Ac1PIM2
Ac2PIM2
Ac2PIM3
Cardiolipins
LPA
LPE
LPG
LPI
PIM1
PA
PE
PG
PI
PIM2
PS
DAT
Ac2SGL
GPL I
GPL IIa/IV
GPL IIb
GPL IIIa
GPL IIIb
GPL serovar 1
GPL V/VI
TAT
alpha MA
fatty acids
mycolactone
mycolipodienic acids
mycosanoic acids
phthienoic acids
w1-methoxy MA
dideoxycarboxymycobactin
dideoxymycobactin
mann-phosphomycoketide
menaquinone
monoglycosylated PGL B2
mycobactins
PDIM B
DAG
MAG
TAG
C
Untreated lysosomal lipidome
Mol %
Fatty Acyls
Glycerolipids
Glycerophospholipids
Sphingolipids
Sterol lipids
Polyketides & Prenol lipids
Eicosanoids
Branched fatty acids
Dicarboxylic acids
Hydroxy fatty acids
Straight chain fatty acids
Unsaturated fatty acids
Fatty Alcohols
Fatty Aldehydes
N-acyl amino acids
N-acyl ethanolamines
Fatty Acyl Carnitines
Fatty acyl CoAs
Short fatty esters
Wax monoesters
Octadecanoids
Diacylglycerols
Glycosyldiacylglycerols
Monoacylglycerols
Monoalkylglycerols
Triacylglycerols
Glycerophosphates
Glycerophosphocholines
Glycerophosphoethanolamines
Glycerophosphoglycerols
Glycerophosphoinositols
Glycerophosphoserines
Oxidized glycerophospholipids
Ceramides
Neutral glycosphingolipids
Phosphosphingolipids
Sphingoid bases
Bile acids & derivatives
Steroid conjugates
Steroids
Sterols
Phenolic lipids
Isoprenoids
D
MEV-infected lysosomal lipidome
Mol %
Glycerophospholipids
Saccharolipids
Fatty Acyls
Polyketides & Prenols
Ac1PIM1
Ac2PIM2
Cardiolipins
LPA
LPE
LPG
LPI
Lyso-PIM1
Ac2SGL
GPL I
GPL IIa/IV
GPL IIb
GPL IIIa
GPL IIIb
GPL serovar 1
GPL V/VI
TAT
Ac3SGL
alpha MA
mycolipodienic acid
mycosanoic acids
phthienoic acids
w1-methoxy MA
dideoxy mycobactins
mann-phosphomycoketide
monodeoxy mycobactins
mycobactins
PDIM B
E
Mol %
Lysosomal lipidome
Lysosomal lipidome post MEV fusion
**
<20
20-30
30-40
>40
Alkyl chain length
F
Mol %
Lysosomal lipidome
Lysosomal lipidome post MEV fusion
**
0
1
2
3
>4
Levels of Unsaturation

**Fig. S13.**

**Lipidomic characterisation of MEVs and MEV-fused lysosomes. reveals enrichment of MEV-derived lipids in host lysosome.**

**(a, b)** Lipidomic profiling of MEVs annotated using the MtbLipidDB database. Class-wise distribution of lipid species in MEV extracts reveals a predominance of fatty acyls, followed by glycerophospholipids, saccharolipids, and additional minor lipid classes. The compositional landscape highlights the presence of *Mycobacterium*-specific lipid families, underscoring the distinct biochemical signature of MEVs derived from mycobacterial membranes. Data are presented as mean ± SEM from six independent replicates.

**(c)** Class-wise lipidomic profile of untreated (only PBS treated) lysosomal lipid extracts, providing a baseline for comparison with MEV-fused lysosomes.

**(d)** Lipid signatures characteristic of *Mycobacterium tuberculosis* (Mtb), identified in MEV-fused purified lysosomes and annotated using the MtbLipidDB database. Each bar represents mean lipid abundance (± SEM) from six technical replicates. The detection of Mtb-specific lipid classes in fused lysosomes indicates successful transfer and persistence of bacterial lipids within host lysosomal compartments following MEV fusion.

**(e)** Distribution of acyl chain lengths and **(f)** degree of unsaturation of lipid species identified in isolated lysosomes (green) and MEV-fused lysosomal fractions (blue). Data represent mean ± SEM from six technical replicates. (**p<0.01, ***p<0.001 in one-way ANOVA).

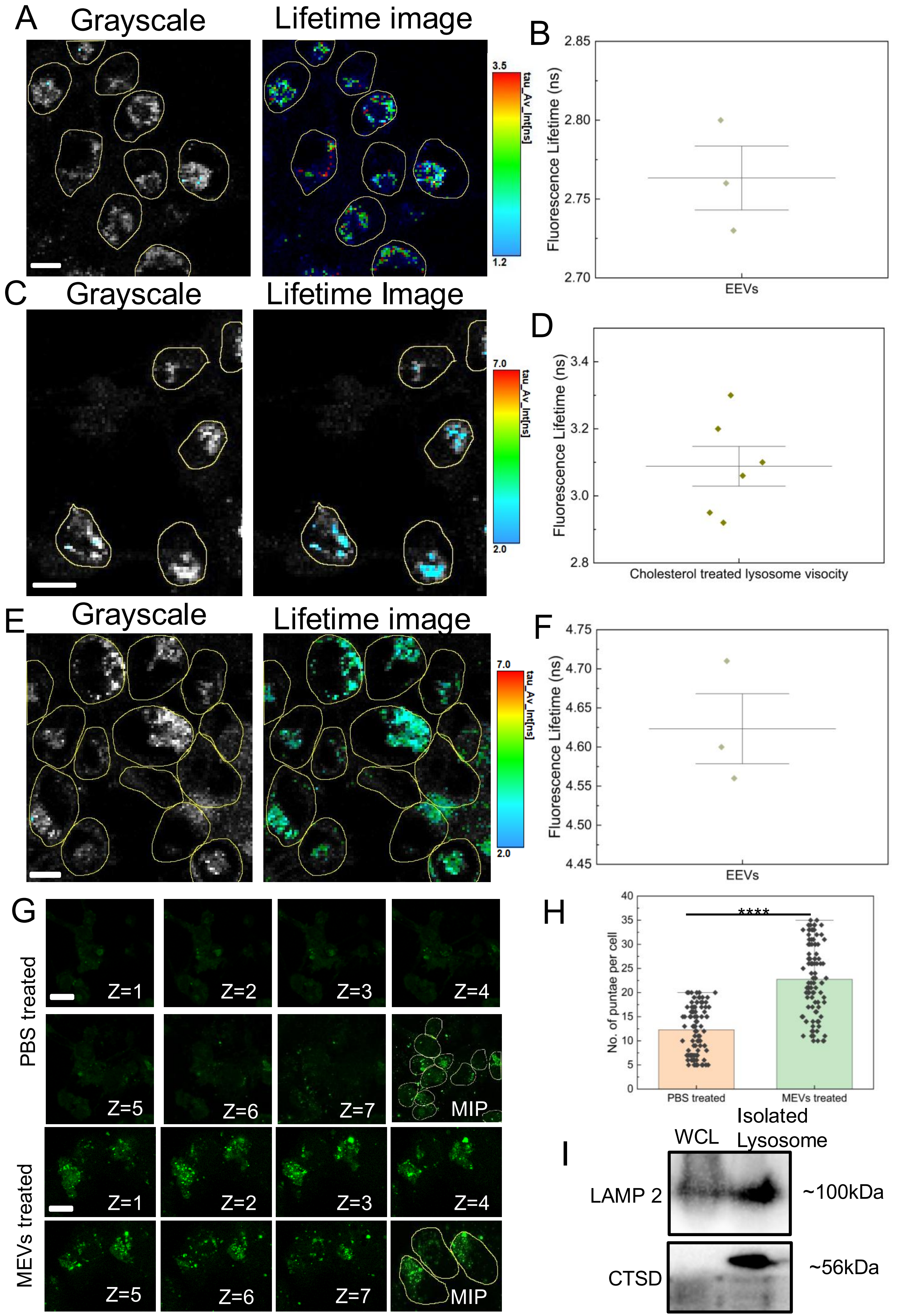
A
Grayscale
Lifetime image
3.5
1.2
tau_Av_Int[ns]
B
Fluorescence Lifetime (ns)
2.85
2.80
2.75
2.70
EEVs
C
Grayscale
Lifetime Image
7.0
2.0
tau_Av_Int[ns]
D
Fluorescence Lifetime (ns)
3.4
3.2
3.0
2.8
Cholesterol treated lysosome visocity
E
Grayscale
Lifetime image
7.0
2.0
tau_Av_Int[ns]
F
Fluorescence Lifetime (ns)
4.75
4.70
4.65
4.60
4.55
4.50
4.45
EEVs
G
PBS treated
Z=1
Z=2
Z=3
Z=4
Z=5
Z=6
Z=7
MIP
MEVs treated
Z=1
Z=2
Z=3
Z=4
Z=5
Z=6
Z=7
MIP
H
No. of puntae per cell
****
40
35
30
25
20
15
10
5
0
PBS treated
MEVs treated
I
WCL
Isolated Lysosome
LAMP 2
~100kDa
CTSD
~56kDa

**Fig. S14.**
Fluorescence lifetime imaging microscopy (FLIM) analysis of lysosomal viscosity in macrophages treated with EEVs. **(A)** EEV-treated macrophages, macrophages stained with JIND-Mor, a lysosomal viscosity sensor. All measurements were acquired 4 hours post-treatment. Images are representative of three independent biological replicates (~150 cells).

**(B)** Quantification of fluorescence lifetimes corresponding to the conditions shown in (a). Each data point represents the mean lifetime from an independent biological replicate.

**(C)** Representative FLIM images of cholesterol–treated macrophages stained with JIND-Mor. Images are representative of six independent biological replicates (~100 cells).

**(D)** Quantification of fluorescence lifetimes corresponding to the conditions shown in (c). Each data point represents the mean lifetime from an independent biological replicate.

**(E)** FLIM analysis of lysosomal membrane tension in macrophages treated with different MEVs. EEVs-treated macrophages, stained with Lyso-Flipper, a lysosomal membrane tension sensor. All measurements were acquired 4 hours post-treatment. Images are representative of three independent biological replicates (~145 cells).

**(F)** Quantification of fluorescence lifetimes corresponding to the conditions. Each data point represents the mean lifetime from an independent biological replicate.

**(G)** Z-stack confocal imaging of THP-1–derived macrophages loaded with lucifer yellow CH lysosomes in the presence and absence of MEVs. Representative images of live macrophages showing a greater number of puncta in the MEV-treated macrophages. Scale bar, 10 μm (n~100 cells in each condition). Images are representative of three independent biological experiments.

**(H)** Quantification of the number of puncta per cell corresponding to the conditions. (**p<0.01, ***p<0.001 in one-way ANOVA).

**(I)** Representative images of western blot of LAMP2 (~100 kDa) and CTSD (~56 kDa) proteins showing the purity of isolated lysosome fractions compared to the remaining whole cell lysate. These are representative blots of three independent replicates.

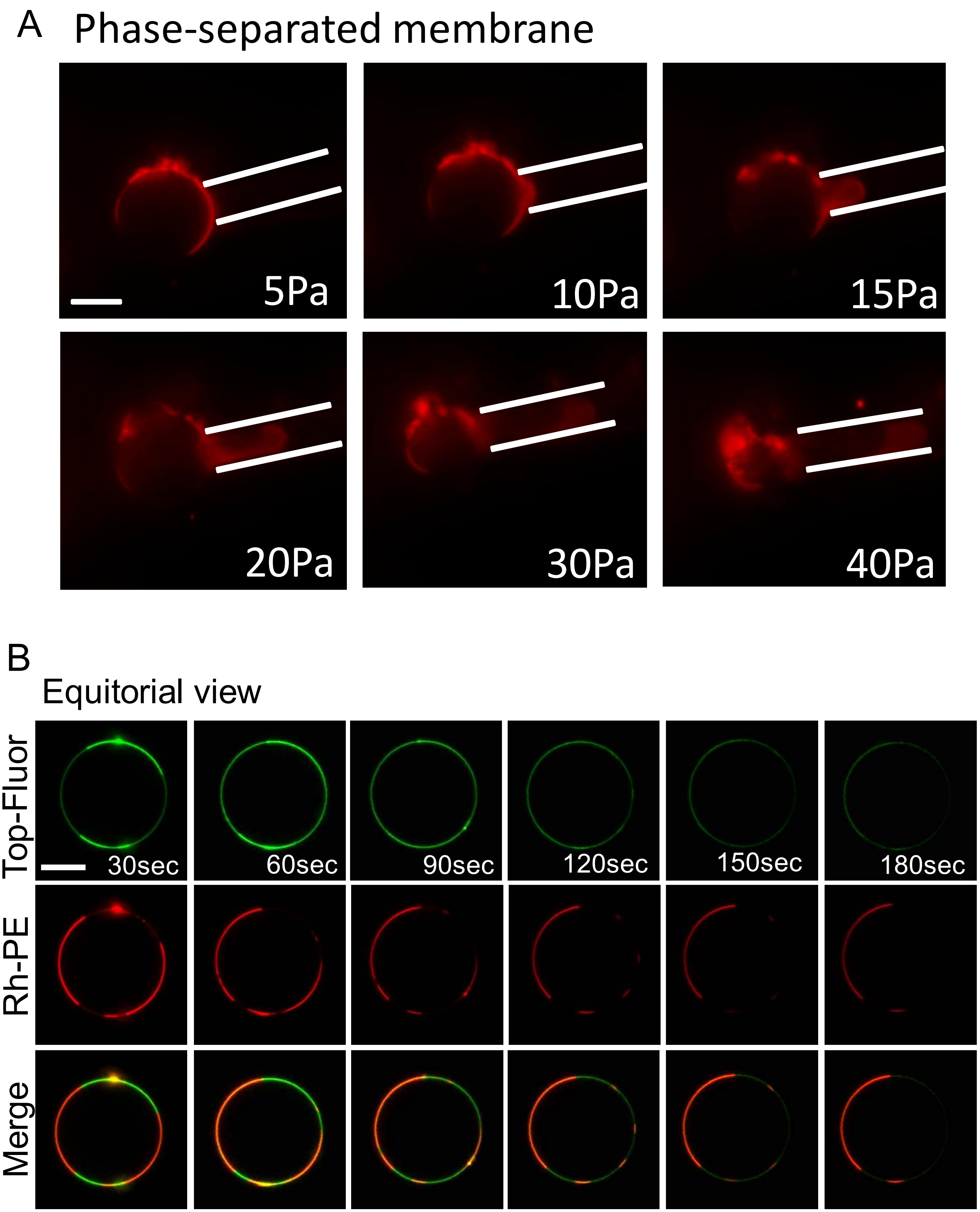
A
Phase-separated membrane
5Pa
10Pa
15Pa
20Pa
30Pa
40Pa
B
Equitorial view
Top-Fluor
Rh-PE
Merge
30sec
60sec
90sec
120sec
150sec
180sec

**Fig. S15.**

**(A)** Time-lapse fluorescence imaging of phase-separated GUVs labelled with 0.1% Rh-PE (red) during micropipette aspiration under increasing negative pressure. Scale bar, 5 µm. Images are representative of three independent experiments

**(B)** Time-lapse confocal imaging of phagosomal membrane–mimicking GUVs doped with 25 mol% MEV-derived lipids and labelled with 0.01% Rh-PE (red) and 0.01% TopFluor-Cholesterol (green). Images (equatorial plane) illustrate the emergence of lipid phase separation upon MEV lipid incorporation. Representative frames from five independent biological replicates are shown. All experiments were performed in PBS (pH 7.4) at 37 °C. Scale bar, 5 µm.
